## Supporting Information Folder for "Characterizing the landscape of gene expression variance in humans": readme.pdf (from another version of this preprint: version 2, 2023-01-09)

### Description of Study-Wise Metadata Table Columns

| Column ID | Column Title | Description |
|---|---|---|
| id | Study ID | Study ID |
| group | Study Group | Study group identifying whether the study belongs to GTEx, TCGA, or neither. |
| tissue | Tissue | Tissue samples were generated from |
| date | Datetime of Row Generation | Datetime of row creation -- note that this is not the datetime the study was generated. |
| raw_counts_genes | Number of Genes in Raw Data | Number of genes in raw count data before any filtering occurs. |
| raw_counts_individuals | Number of Individuals in Raw Data | Number of individuals in raw count data before any filtering occurs. |
| raw_metadata_individuals | Number of Individuals in Raw Metadata | Number of individuals in raw metadata. This number should always be the same as raw_counts_individuals. |
| raw_metadata_features | Number of Features in Raw Metadata | Number of features in raw metadata. |
| filtered_counts_genes | Number of Genes after Filtering | Number of genes after filtering |
| filtered_counts_individuals | Number of Individuals after Filtering | Number of individuals after filtering |
| filtered_metadata_individuals | Number of Individuals after Filtering | Number of individuals after filtering -- this should be the same as filtered_counts_individuals. |
| filtered_metadata_features | Number of Features in Metadata after Filtering | Number of features left in the metadata after filtering. |
| to_ignore_manually_curated | Metadata Features to Ignore -- Manually Curated List by Study | The manually curated list of metadata columns to drop from this study. |

| Column ID | Column Title | Description |
|---|---|---|
| to_ignore_redundant_features | Metadata Features Identified as Redundant and Removed | Features identified as a duplicate of another feature. When this happens, we drop all except a single representative feature. Here we report the columns dropped from this process. |
| to_ignore_large_factors | Metadata Features Removed because number of Levels was above Threshold | Features that are factors and have more than 20 levels are dropped. Here we report features dropped in this process. |
| final_features_removed | Metadata Features Removed According to a Final Set of Filters | We use a final set of filters including high missingness, no levels, small levels, other known bad columns. Here we report the columns dropped by this process. |
| final_metadata_colnames | Remaining Metadata Features | Here we report the resulting features kept for the dataset. |
| replicate_col | Column Marking Replicate Samples | When replicates appear in a dataset, we sum technical replicates. Here we list the column that capture technical replicates in the dataset when one exists. |
