## Supporting Information Folder for "Characterizing the landscape of gene expression variance in humans": readme.pdf (from another version of this preprint: version 2, 2023-01-09)

### Description of Gene Ontology Enrichment Results for High and Low Variance Genes

| Column ID | Description |
| --- | --- |
| ID | Gene Ontology ID |
| Description | Description of Gene Ontology term |
| GeneRatio | Ratio of genes enriched in Gene Ontology term in quantile over total count of genes in quantile |
| BgRatio | Background ratio of genes enriched in Gene Ontology term over total count of genes |
| p.adjust | Adjusted p-value |
| qvalue | q-value |
| geneID | Enriched Gene IDs found in the quantile |
| Count | Count of enriched genes in the quantile |
| varRank | Category of quantile (high/low) |
