## Supporting Information Folder for "Characterizing the landscape of gene expression variance in humans": SI_appendix1_diagnostic_plots.pdf (from another version of this preprint: version 2, 2023-01-09)

ADIPOSE_TISSUE

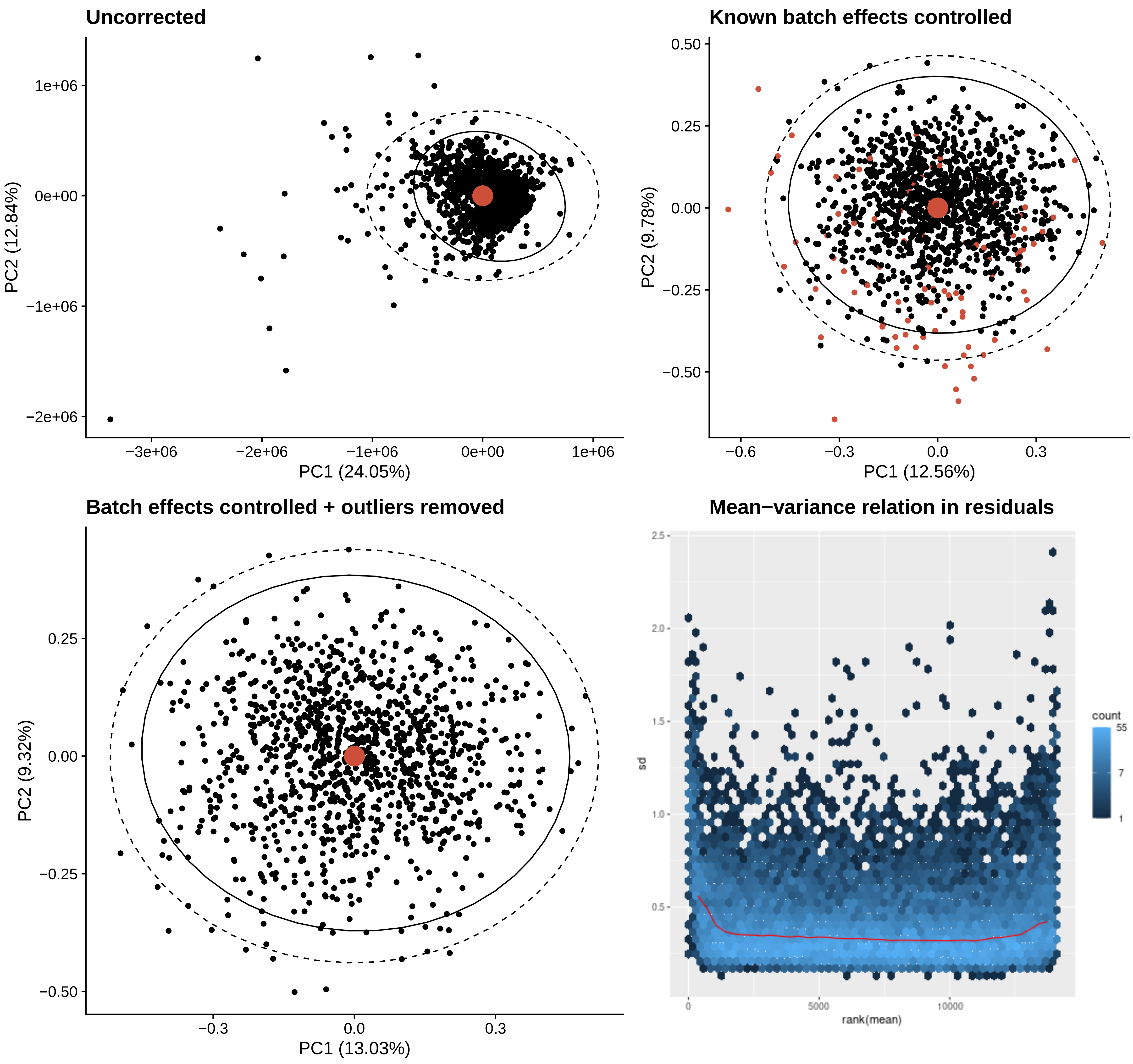

ADRENAL_GLAND

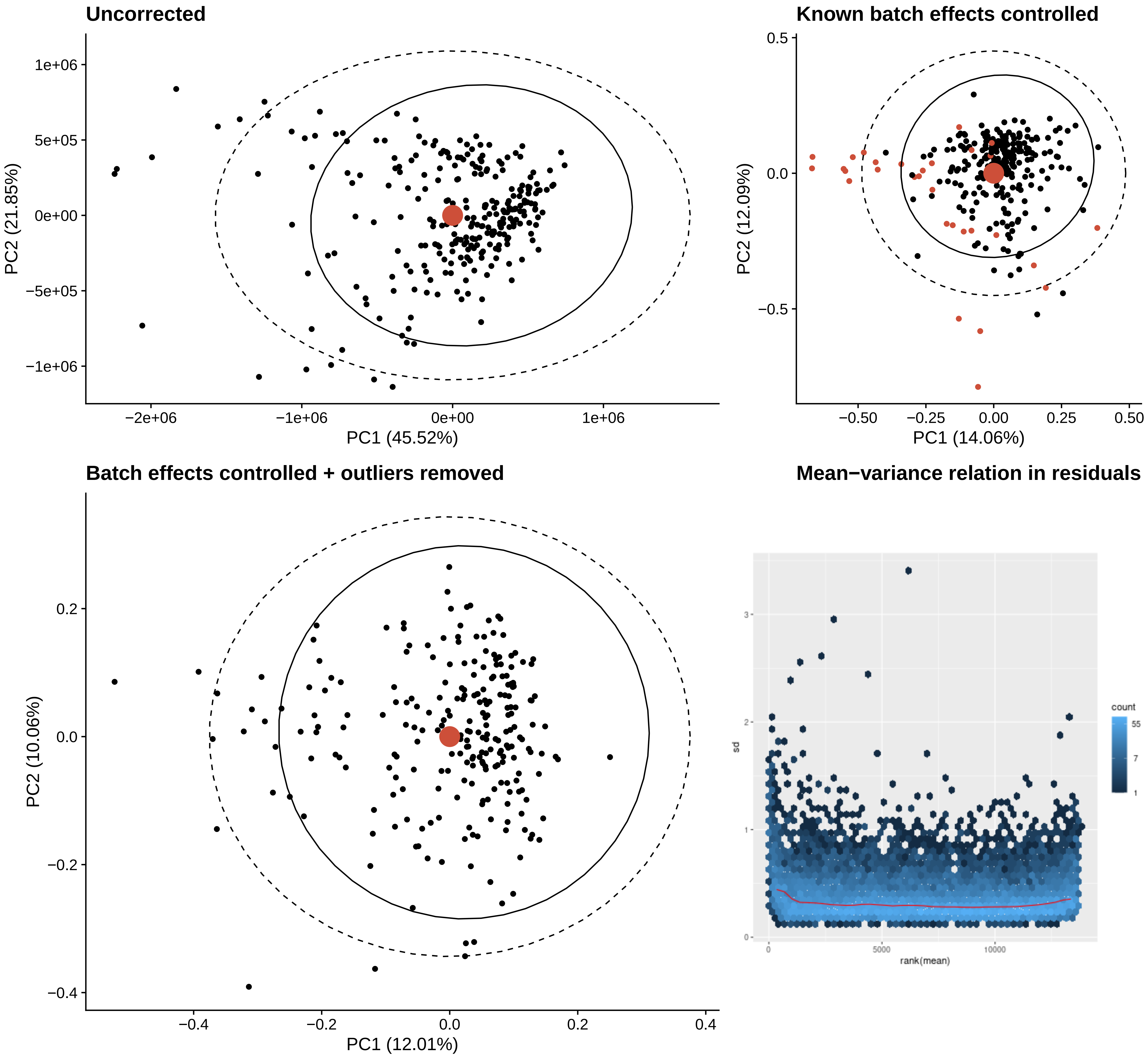

BLOOD

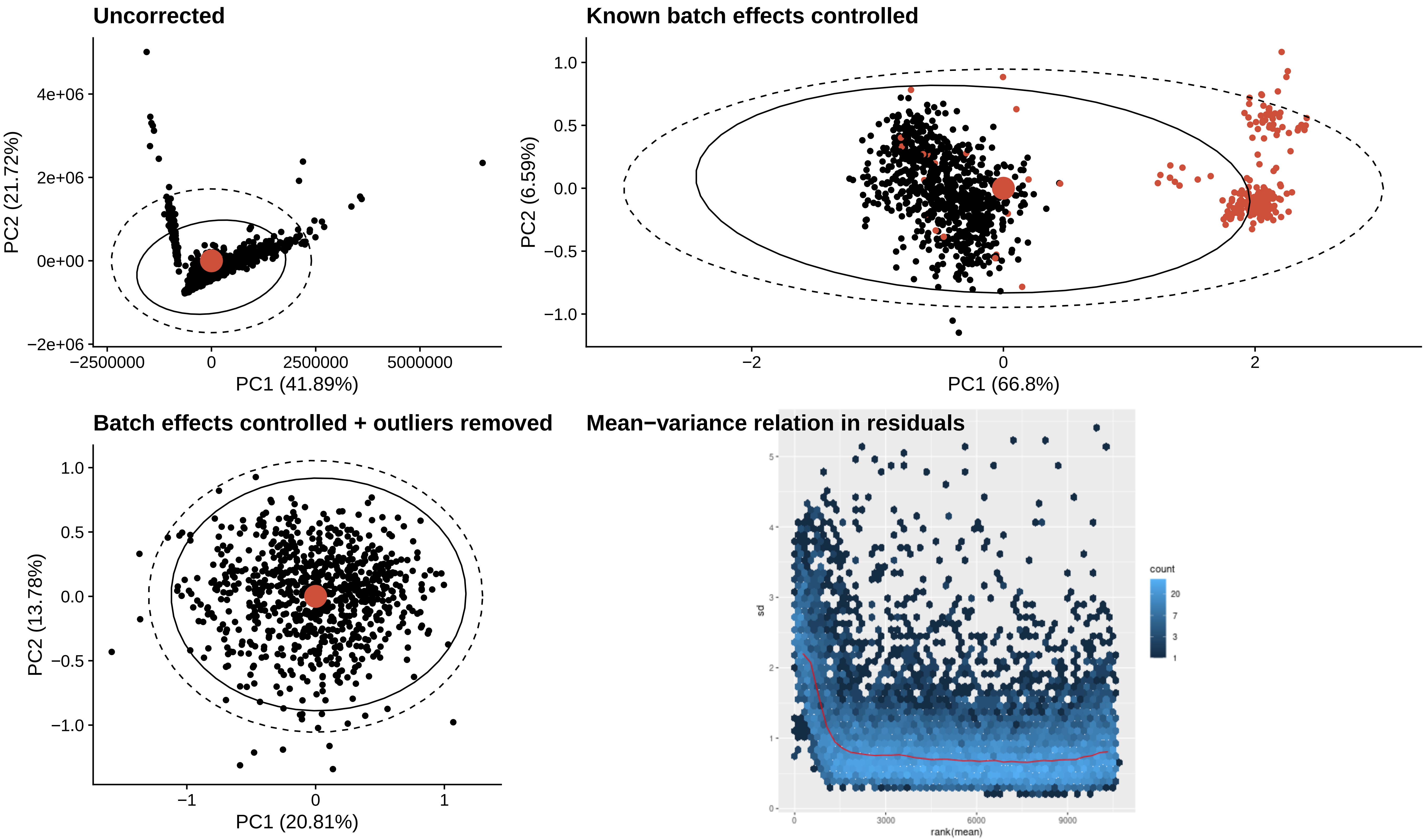

Uncorrected
4e+06
2e+06
0e+00
−2e+06
−2500000
0
2500000
5000000
PC1 (41.89%)
PC2 (21.72%)
Known batch effects controlled
1.0
0.5
0.0
−0.5
−1.0
−2
0
2
PC1 (66.8%)
PC2 (6.59%)
Batch effects controlled + outliers removed
1.0
0.5
0.0
−0.5
−1.0
−1
0
1
PC1 (20.81%)
PC2 (13.78%)
Mean−variance relation in residuals
sd
rank(mean)
0
3000
6000
9000
count
20
7
3
1

BLOOD_VESSEL

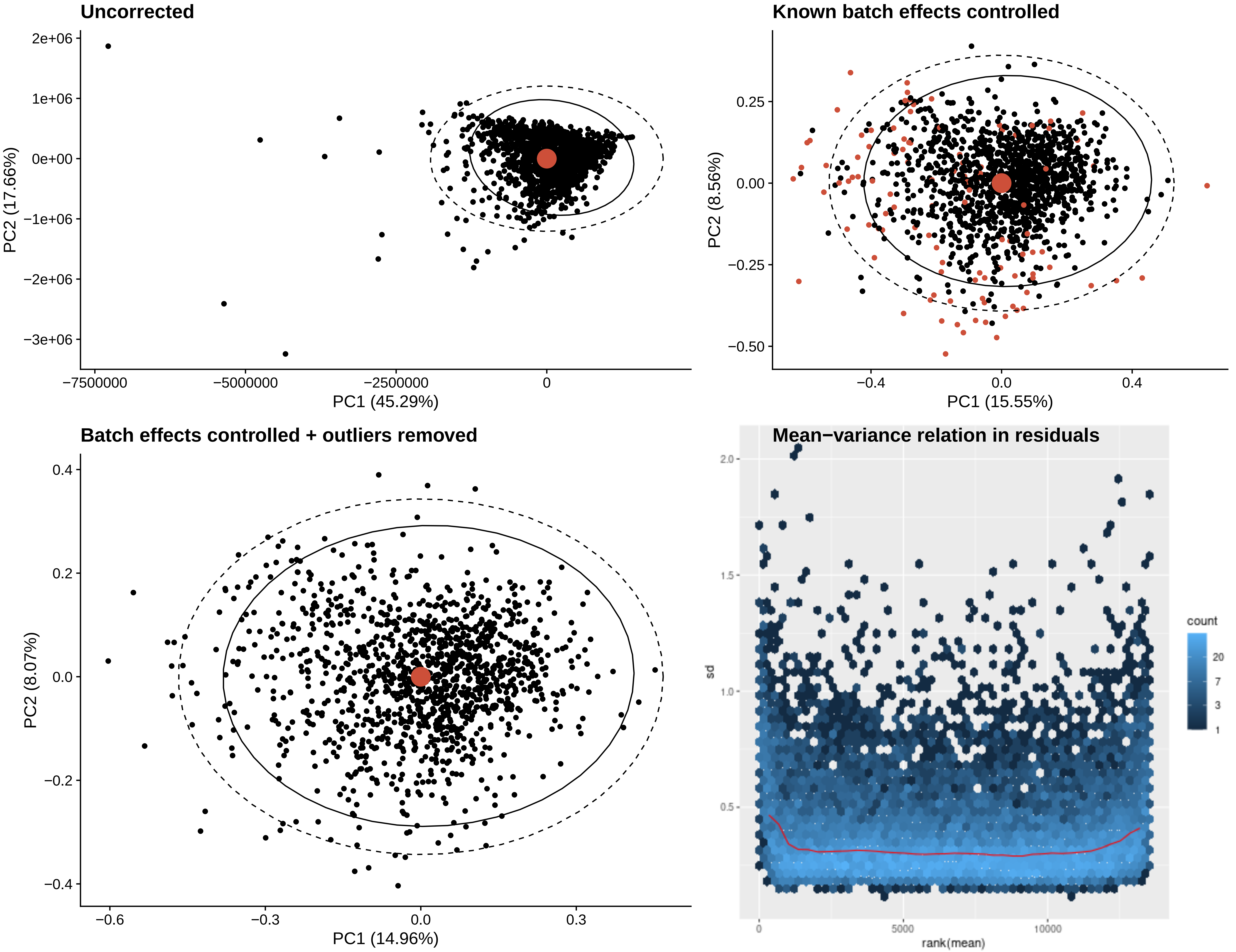

BONE_MARROW

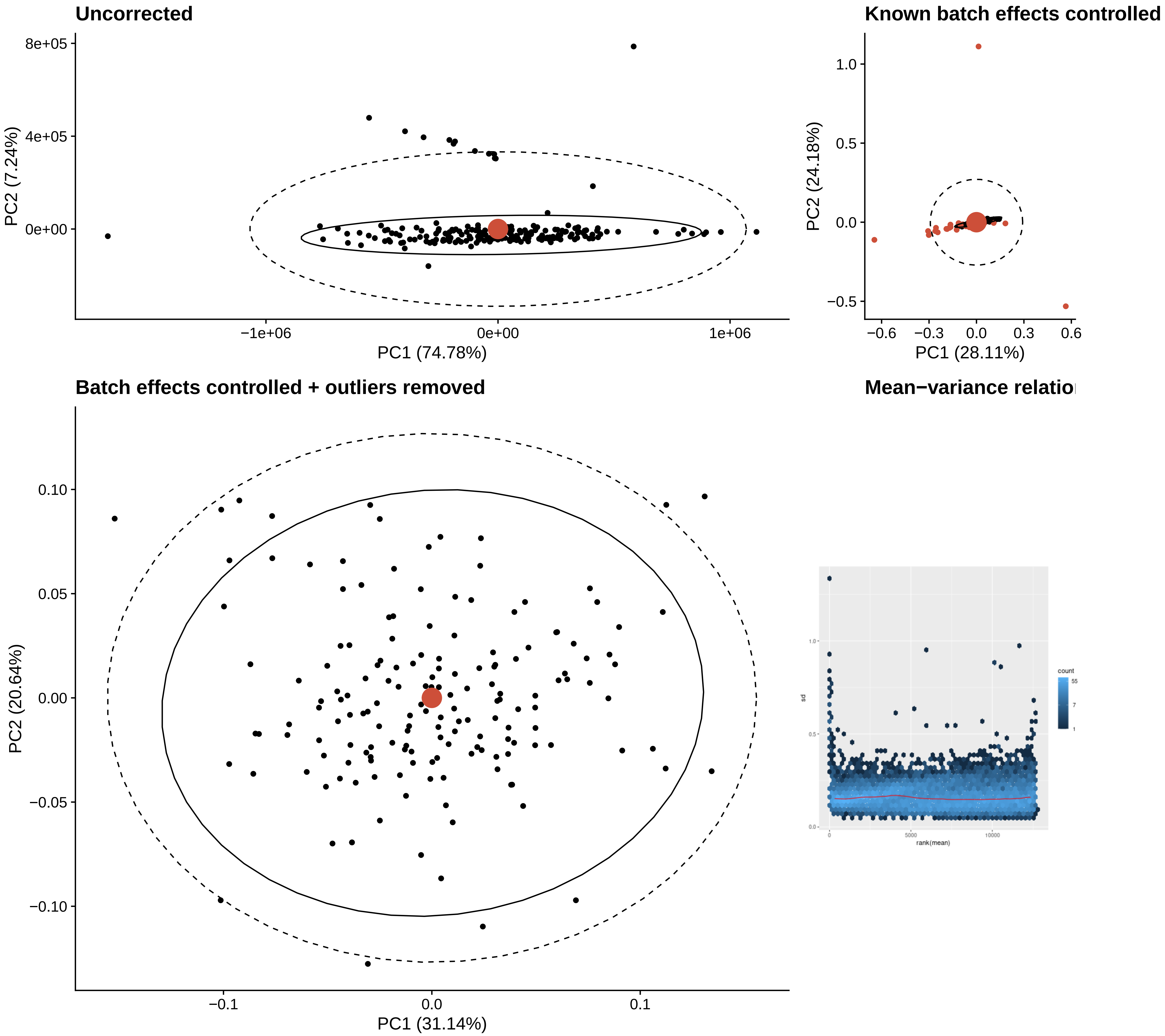

Uncorrected
8e+05
4e+05
0e+00
PC2 (7.24%)
−1e+06
0e+00
1e+06
PC1 (74.78%)
Known batch effects controlled
1.0
0.5
0.0
−0.5
PC2 (24.18%)
−0.6
−0.3
0.0
0.3
0.6
PC1 (28.11%)
Batch effects controlled + outliers removed
0.10
0.05
0.00
−0.05
−0.10
PC2 (20.64%)
−0.1
0.0
0.1
PC1 (31.14%)
Mean−variance relatio
count
sd
rank(mean)

BRAIN

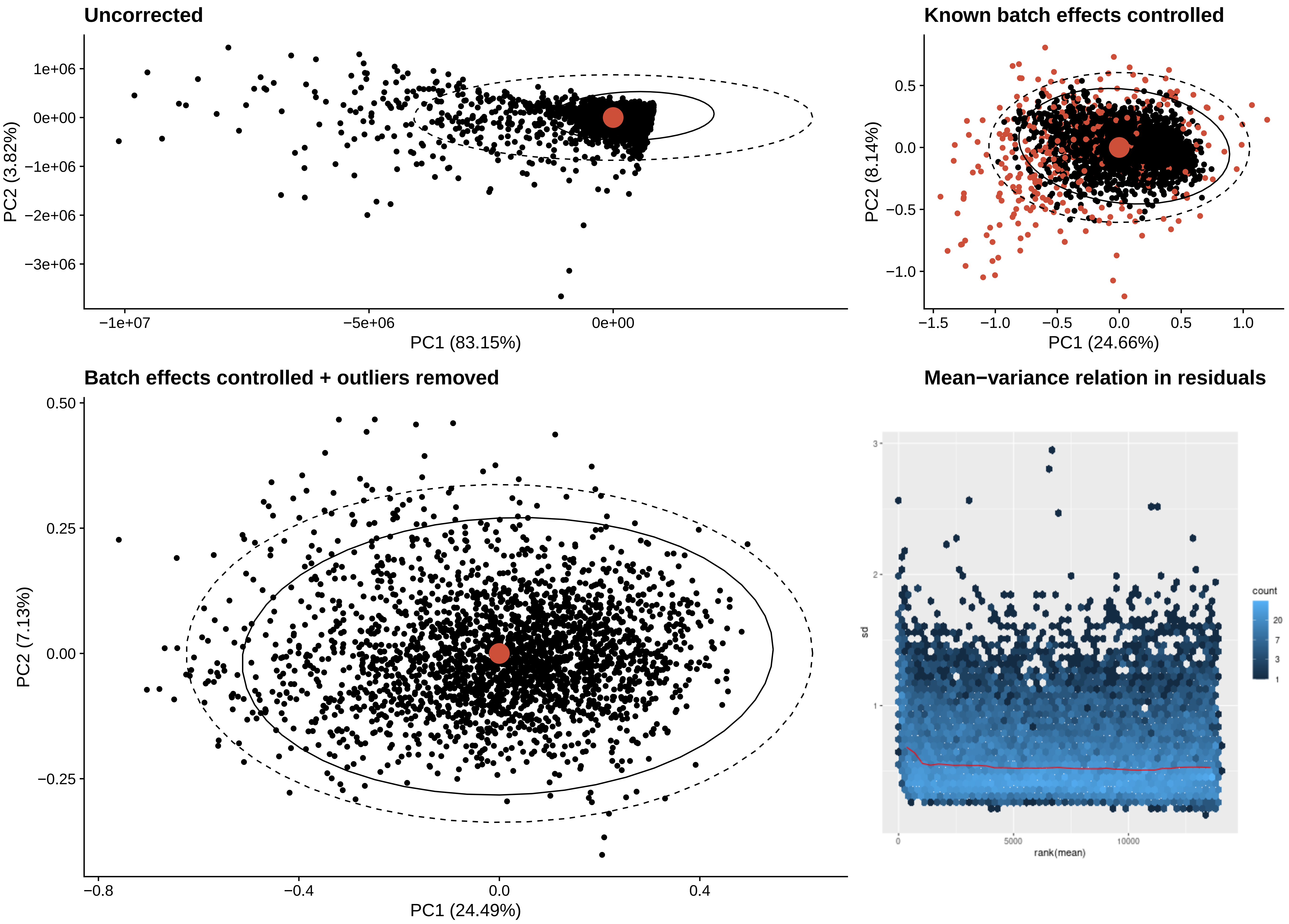

Uncorrected
1e+06
0e+00
−1e+06
−2e+06
−3e+06
PC2 (3.82%)
−1e+07
−5e+06
0e+00
PC1 (83.15%)
Known batch effects controlled
0.5
0.0
−0.5
−1.0
PC2 (8.14%)
−1.5
−1.0
−0.5
0.0
0.5
1.0
PC1 (24.66%)
Batch effects controlled + outliers removed
0.50
0.25
0.00
−0.25
PC2 (7.13%)
−0.8
−0.4
0.0
0.4
PC1 (24.49%)
Mean−variance relation in residuals
sd
rank(mean)
count

BREAST

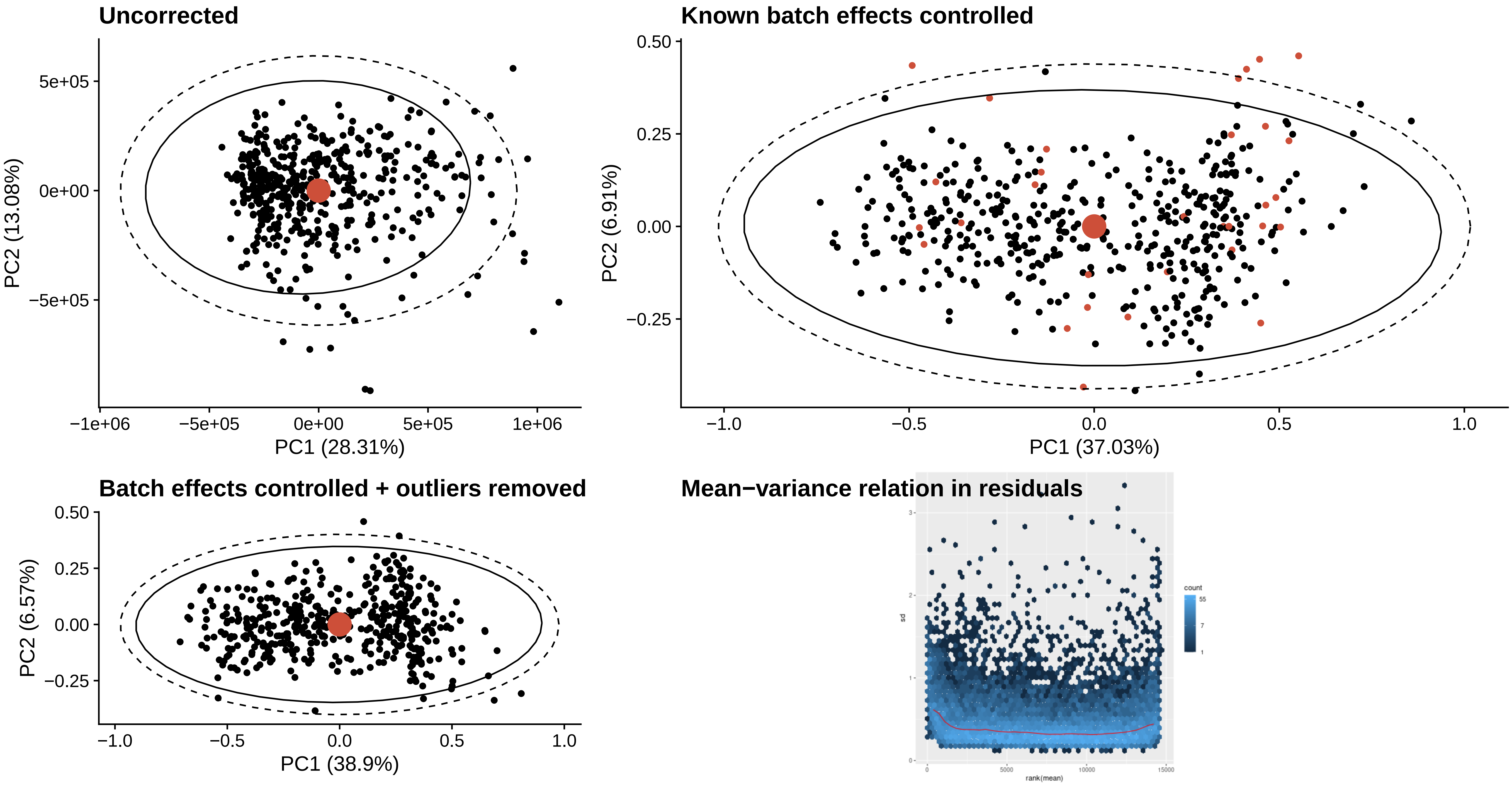

Uncorrected
5e+05
0e+00
−5e+05
PC2 (13.08%)
−1e+06
−5e+05
0e+00
5e+05
1e+06
PC1 (28.31%)
Known batch effects controlled
0.50
0.25
0.00
−0.25
PC2 (6.91%)
−1.0
−0.5
0.0
0.5
1.0
PC1 (37.03%)
Batch effects controlled + outliers removed
0.50
0.25
0.00
−0.25
PC2 (6.57%)
−1.0
−0.5
0.0
0.5
1.0
PC1 (38.9%)
Mean−variance relation in residuals
sd
rank(mean)
count

COLON

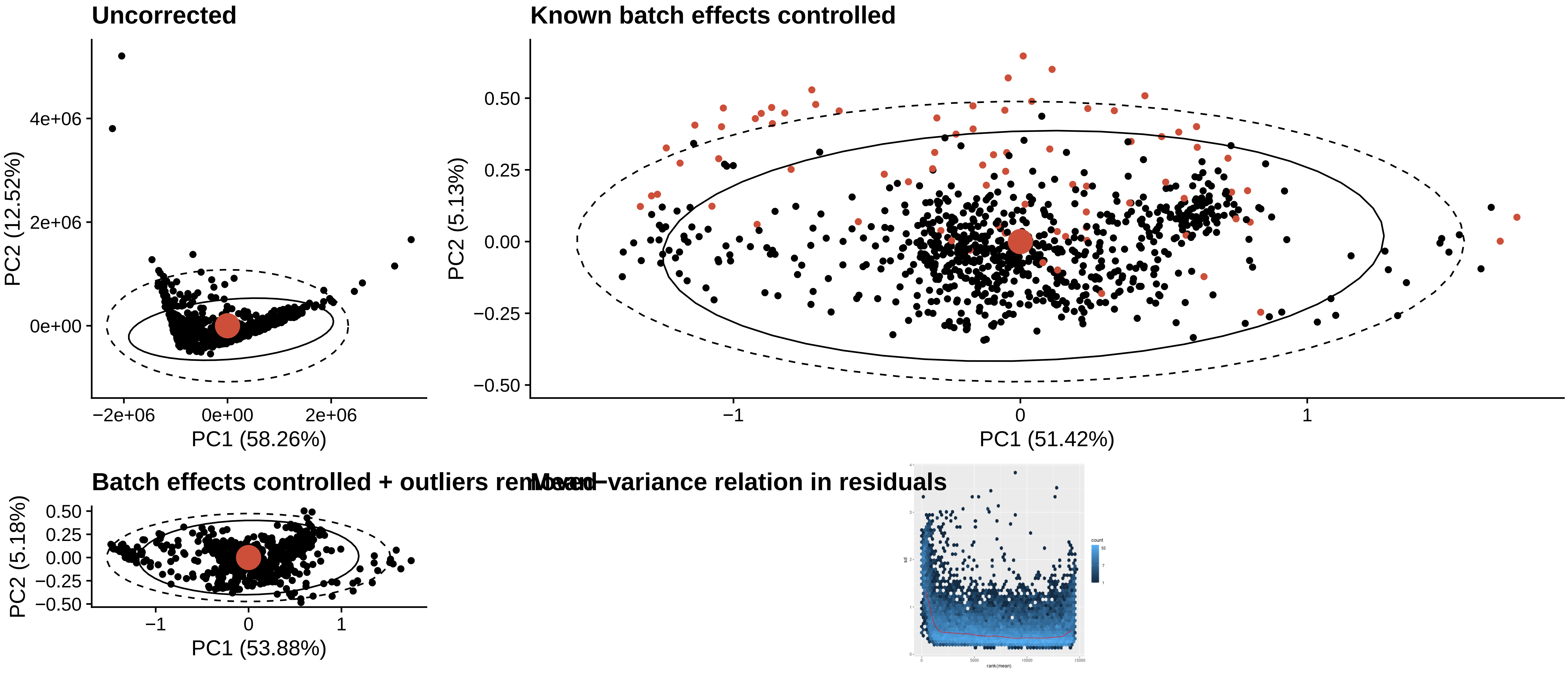
Uncorrected
4e+06
2e+06
0e+00
PC2 (12.52%)
−2e+06
0e+00
2e+06
PC1 (58.26%)
Known batch effects controlled
0.50
0.25
0.00
−0.25
−0.50
PC2 (5.13%)
−1
0
1
PC1 (51.42%)
Batch effects controlled + outliers removed
0.50
0.25
0.00
−0.25
−0.50
PC2 (5.18%)
−1
0
1
PC1 (53.88%)
Mean−variance relation in residuals

ESOPHAGUS

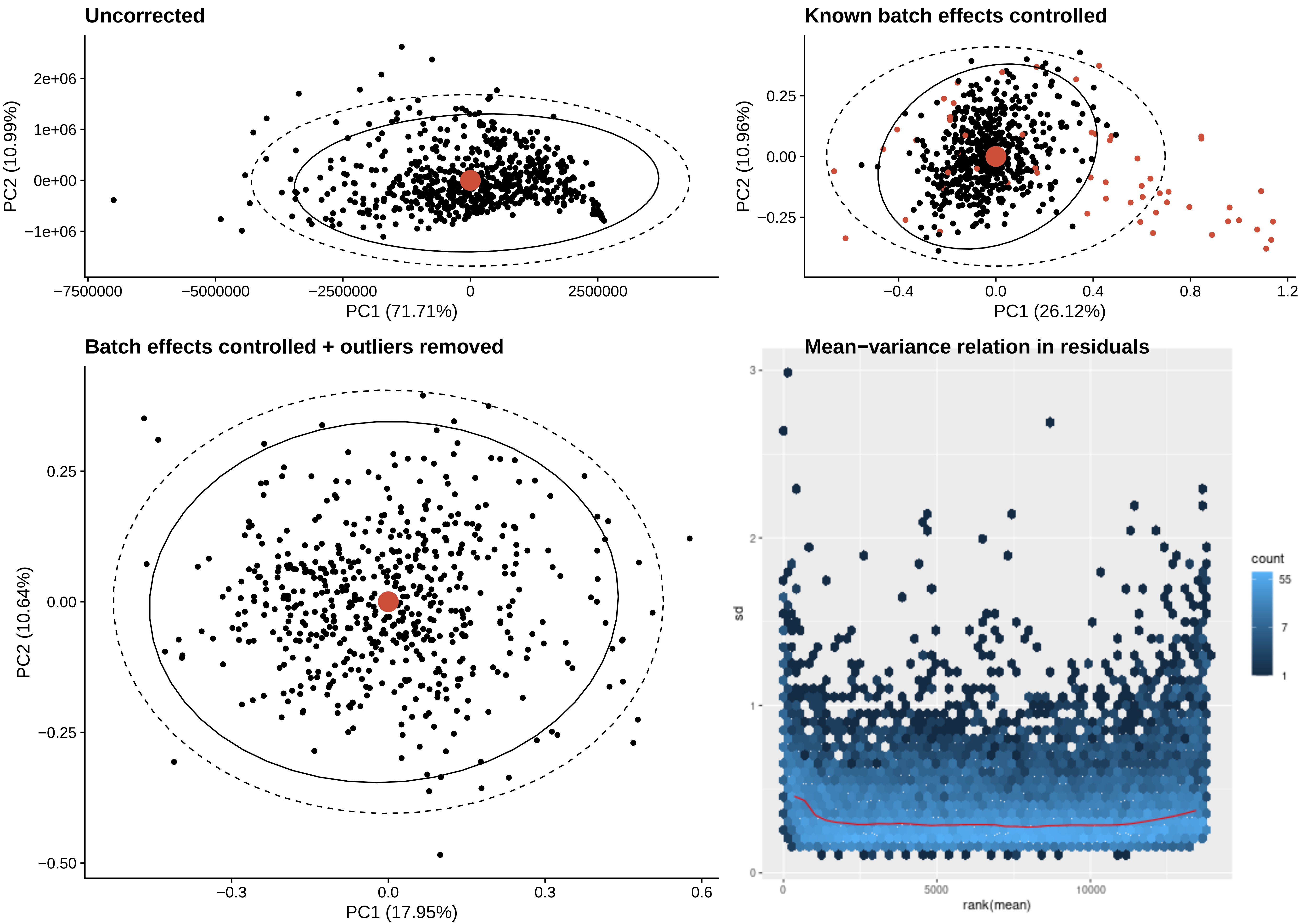

Uncorrected
PC2 (10.99%)
PC1 (71.71%)
Known batch effects controlled
PC2 (10.96%)
PC1 (26.12%)
Batch effects controlled + outliers removed
PC2 (10.64%)
PC1 (17.95%)
Mean−variance relation in residuals
sd
rank(mean)
count

HEART

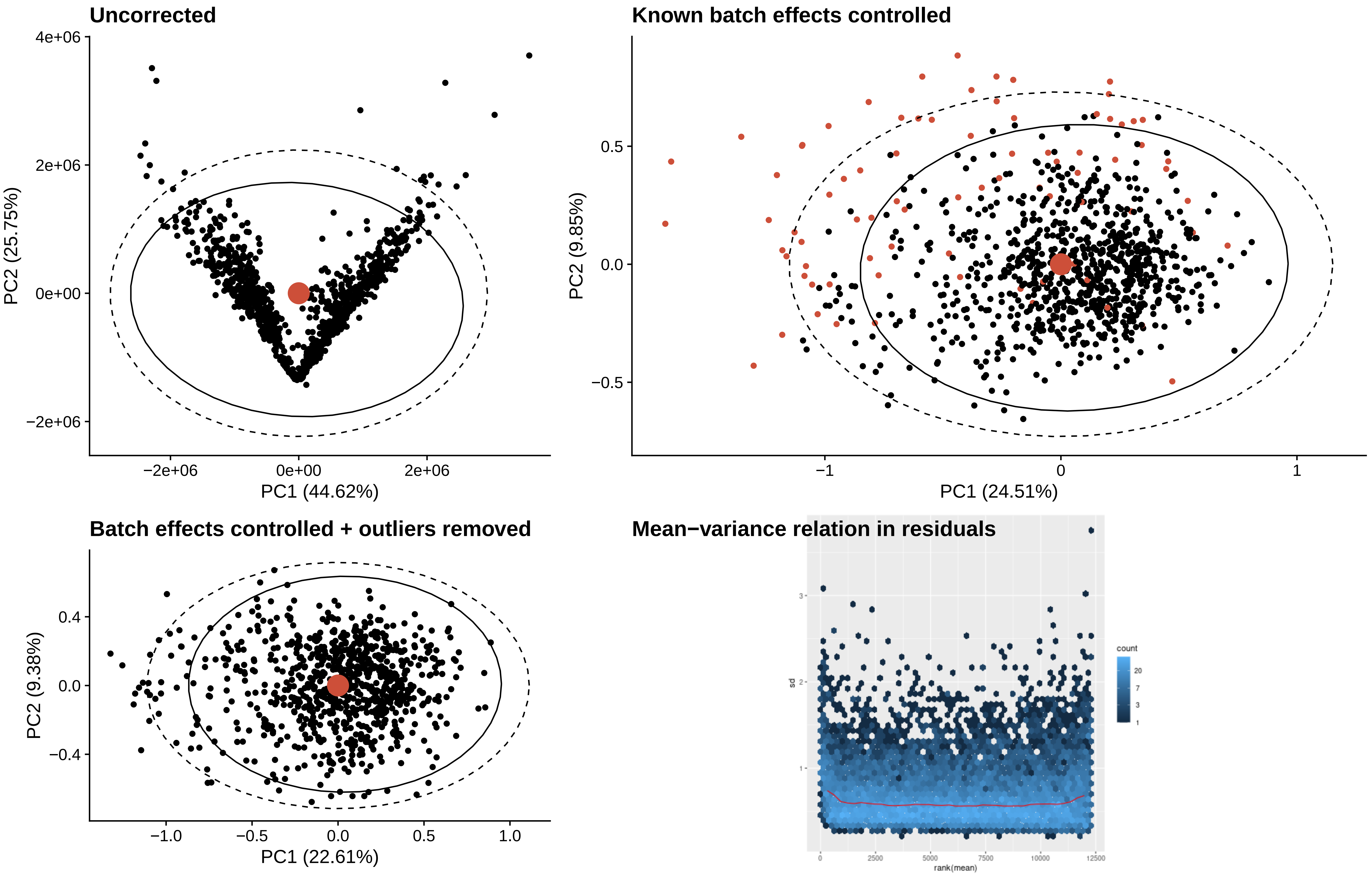
Uncorrected
4e+06
2e+06
0e+00
−2e+06
PC2 (25.75%)
−2e+06
0e+00
2e+06
PC1 (44.62%)
Known batch effects controlled
0.5
0.0
−0.5
PC2 (9.85%)
−1
0
1
PC1 (24.51%)
Batch effects controlled + outliers removed
0.4
0.0
−0.4
PC2 (9.38%)
−1.0
−0.5
0.0
0.5
1.0
PC1 (22.61%)
Mean−variance relation in residuals
sd
3
2
1
0
2500
5000
7500
10000
12500
rank(mean)
count
20
7
3
1

LIVER

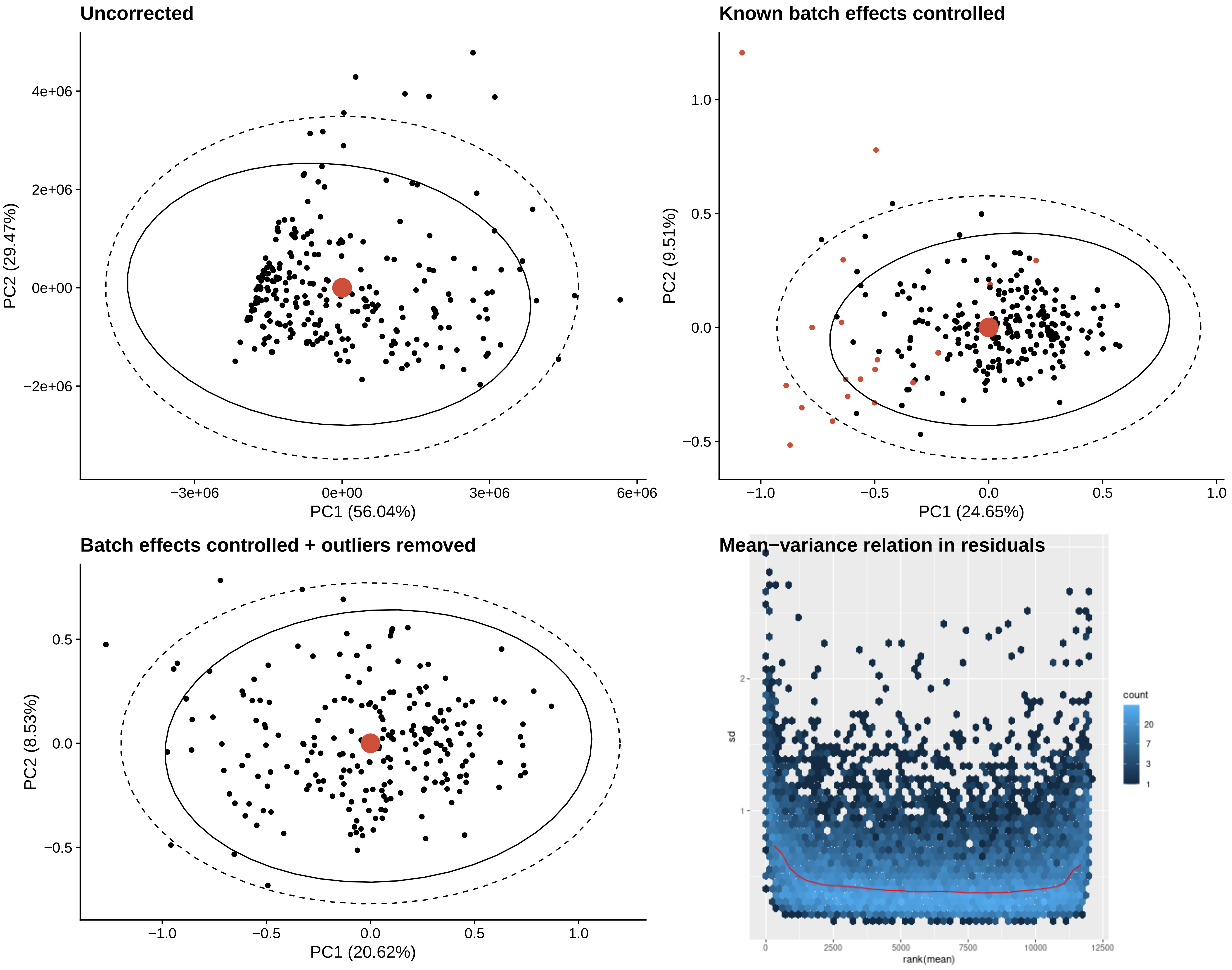

Uncorrected
4e+06
2e+06
0e+00
−2e+06
PC2 (29.47%)
−3e+06
0e+00
3e+06
6e+06
PC1 (56.04%)
Known batch effects controlled
1.0
0.5
0.0
−0.5
PC2 (9.51%)
−1.0
−0.5
0.0
0.5
1.0
PC1 (24.65%)
Batch effects controlled + outliers removed
0.5
0.0
−0.5
PC2 (8.53%)
−1.0
−0.5
0.0
0.5
1.0
PC1 (20.62%)
Mean−variance relation in residuals
sd
2
1
0
2500
5000
7500
10000
12500
rank(mean)
count
20
7
3
1

LUNG

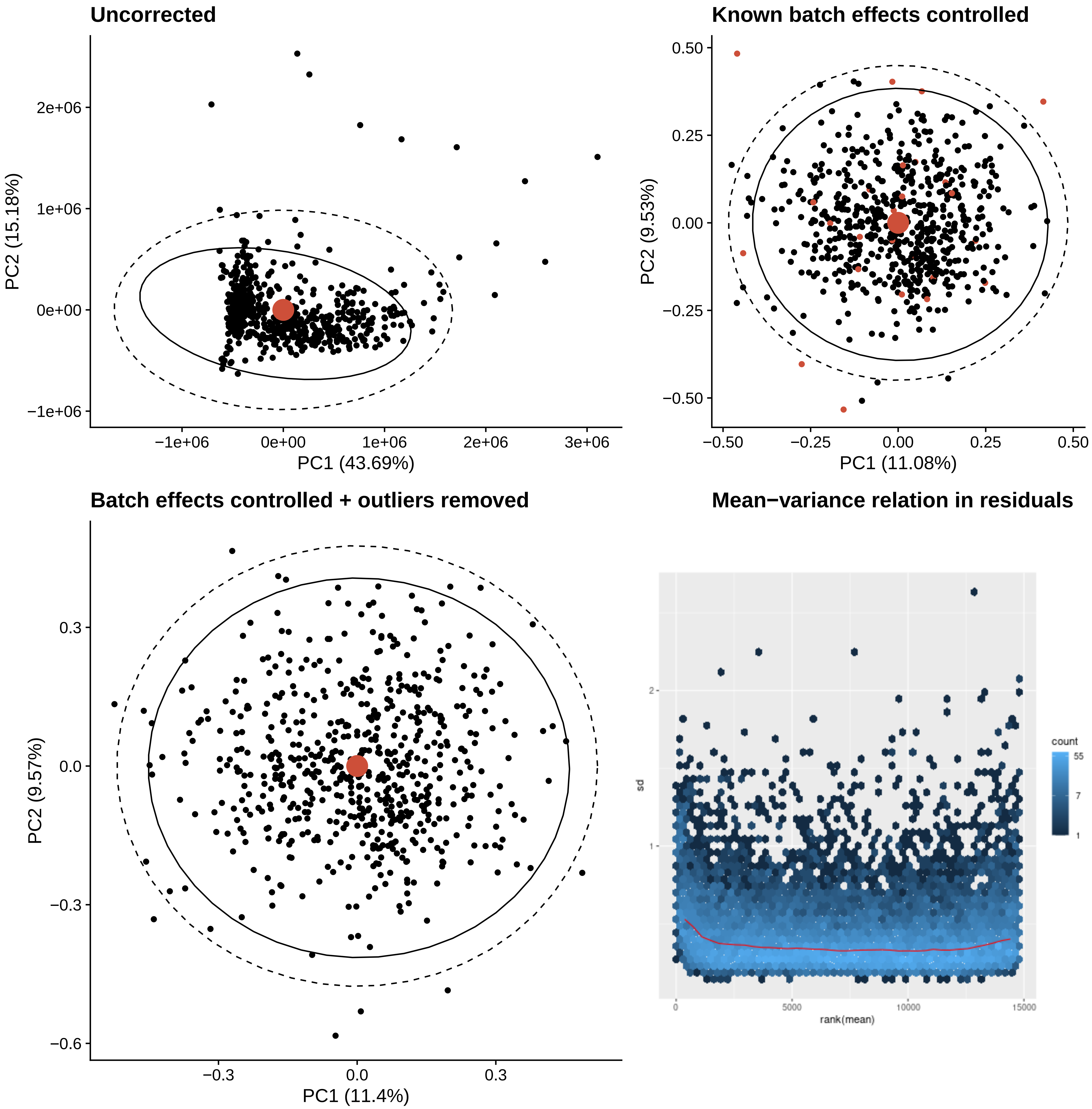

Uncorrected
PC2 (15.18%)
2e+06
1e+06
0e+00
−1e+06
−1e+06
0e+00
1e+06
2e+06
3e+06
PC1 (43.69%)
Known batch effects controlled
PC2 (9.53%)
0.50
0.25
0.00
−0.25
−0.50
−0.50
−0.25
0.00
0.25
0.50
PC1 (11.08%)
Batch effects controlled + outliers removed
PC2 (9.57%)
0.3
0.0
−0.3
−0.6
−0.3
0.0
0.3
PC1 (11.4%)
Mean−variance relation in residuals
sd
2
1
0
5000
10000
15000
rank(mean)
count
55
7
1

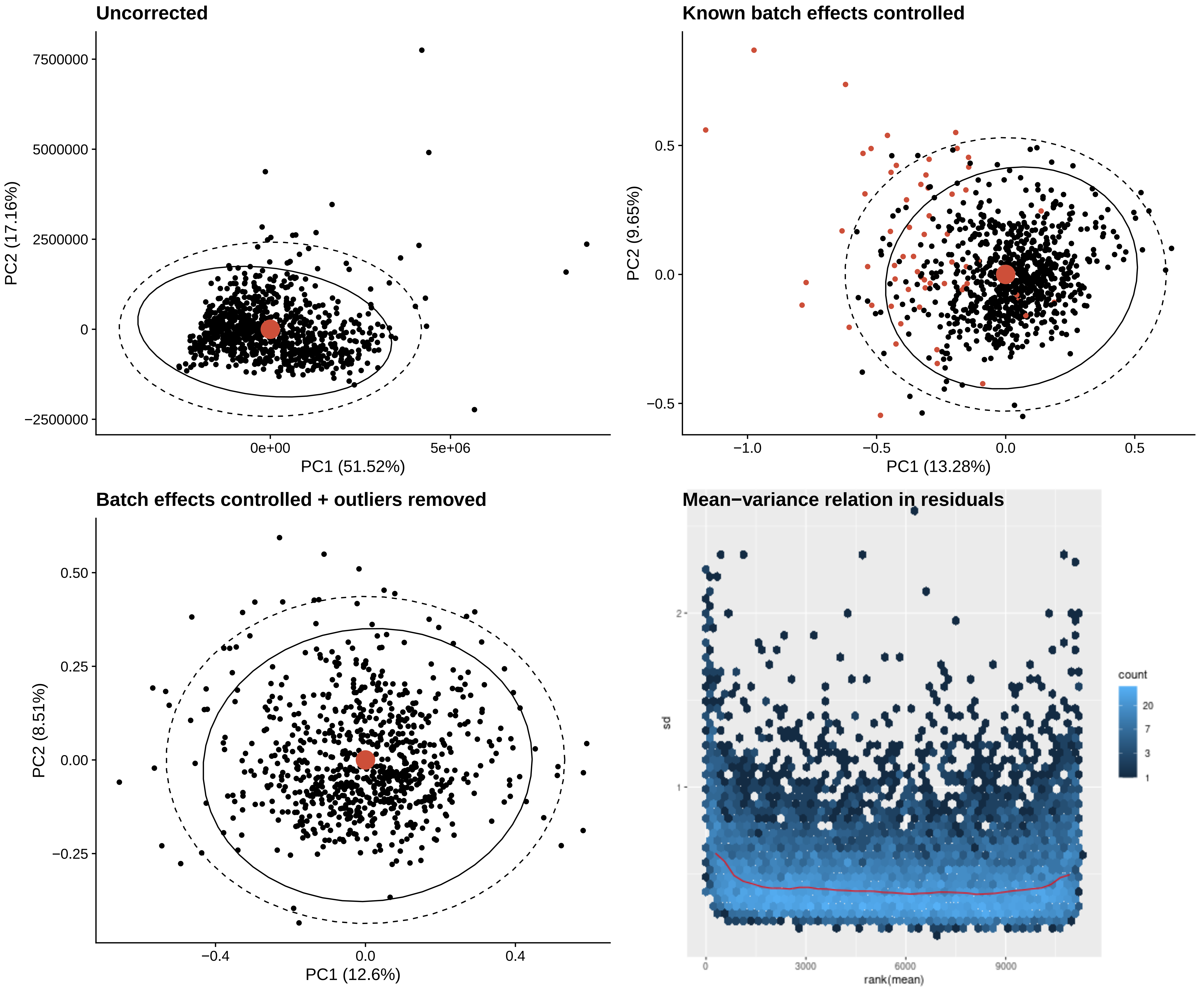
Uncorrected
7500000
5000000
2500000
0
−2500000
PC2 (17.16%)
0e+00
5e+06
PC1 (51.52%)
Known batch effects controlled
0.5
0.0
−0.5
PC2 (9.65%)
−1.0
−0.5
0.0
0.5
PC1 (13.28%)
Batch effects controlled + outliers removed
0.50
0.25
0.00
−0.25
PC2 (8.51%)
−0.4
0.0
0.4
PC1 (12.6%)
Mean−variance relation in residuals
2
1
sd
0
3000
6000
9000
rank(mean)
count
20
7
3
1

NERVE

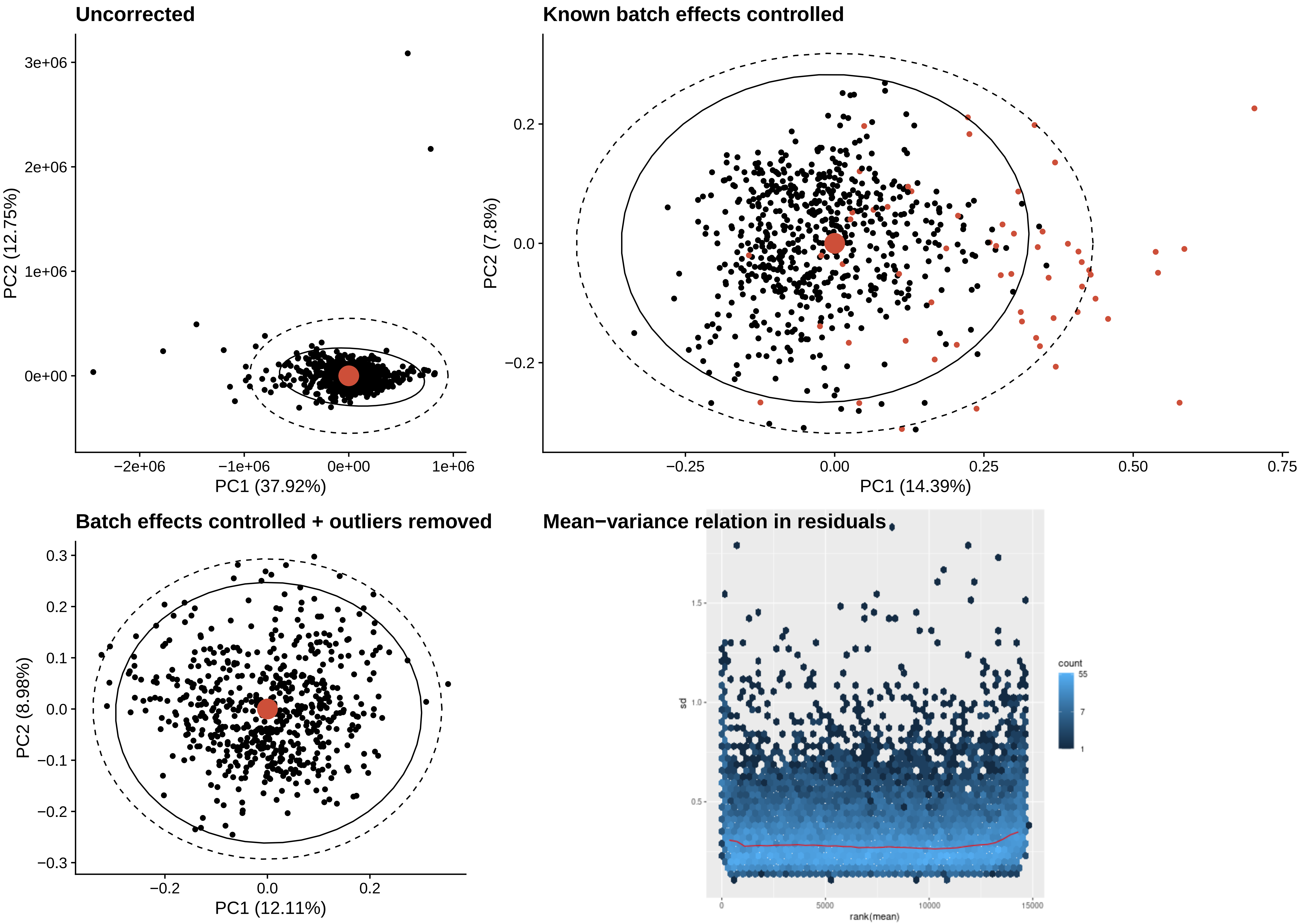
Uncorrected
PC2 (12.75%)
PC1 (37.92%)
3e+06
2e+06
1e+06
0e+00
−2e+06
−1e+06
0e+00
1e+06
Known batch effects controlled
PC2 (7.8%)
PC1 (14.39%)
0.2
0.0
−0.2
−0.25
0.00
0.25
0.50
0.75
Batch effects controlled + outliers removed
PC2 (8.98%)
PC1 (12.11%)
0.3
0.2
0.1
0.0
−0.1
−0.2
−0.3
−0.2
0.0
0.2
Mean−variance relation in residuals
sd
rank(mean)
1.5
1.0
0.5
0
5000
10000
15000
count
55
7
1

OVARY

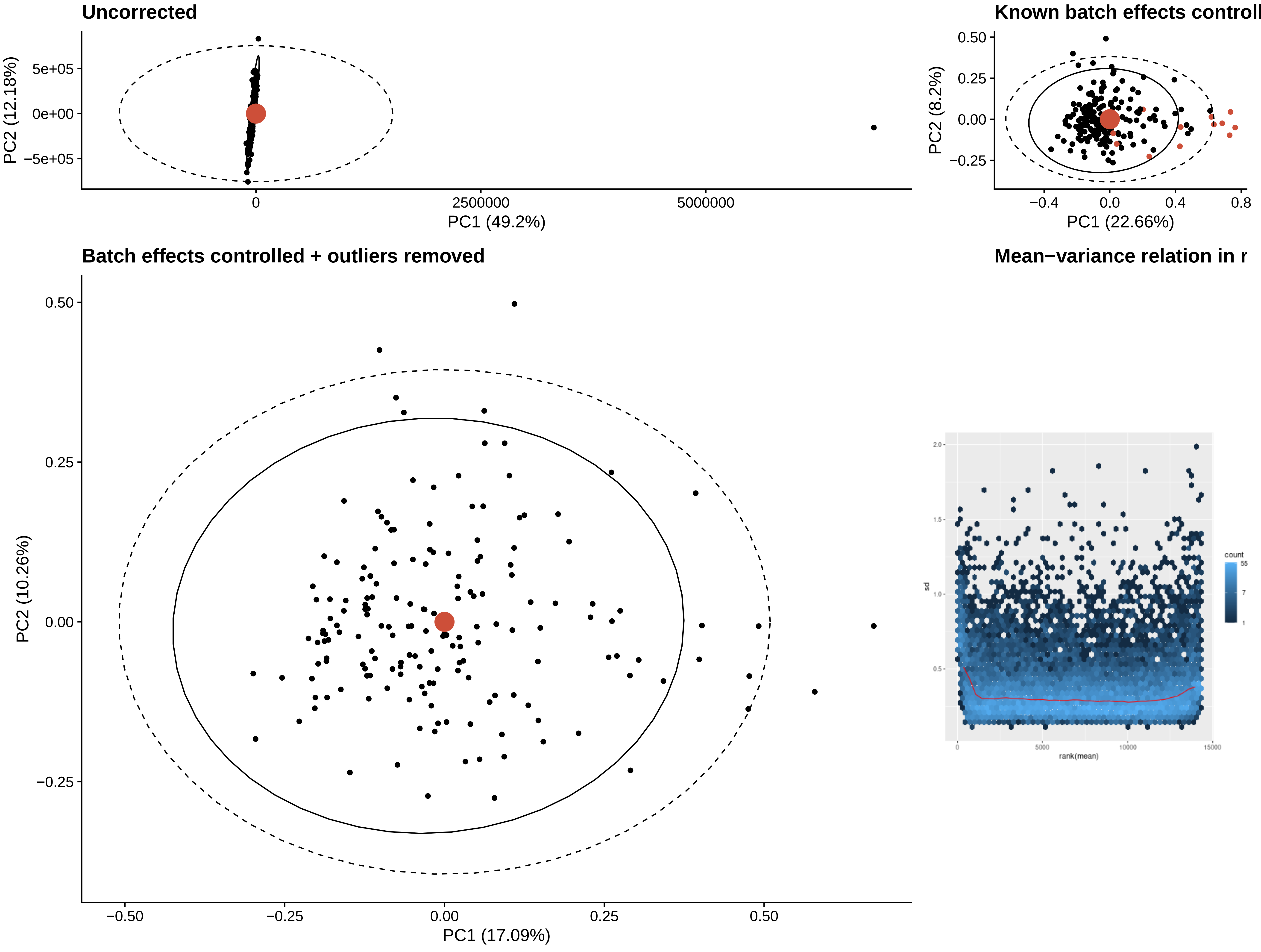
Uncorrected
PC2 (12.18%)
5e+05
0e+00
−5e+05
0
2500000
5000000
PC1 (49.2%)
Known batch effects controll
PC2 (8.2%)
0.50
0.25
0.00
−0.25
−0.4
0.0
0.4
0.8
PC1 (22.66%)
Batch effects controlled + outliers removed
PC2 (10.26%)
0.50
0.25
0.00
−0.25
−0.50
−0.25
0.00
0.25
0.50
PC1 (17.09%)
Mean−variance relation in r
sd
rank(mean)
count
55
7
1
0
5000
10000
15000
2.0
1.5
1.0
0.5

PANCREAS

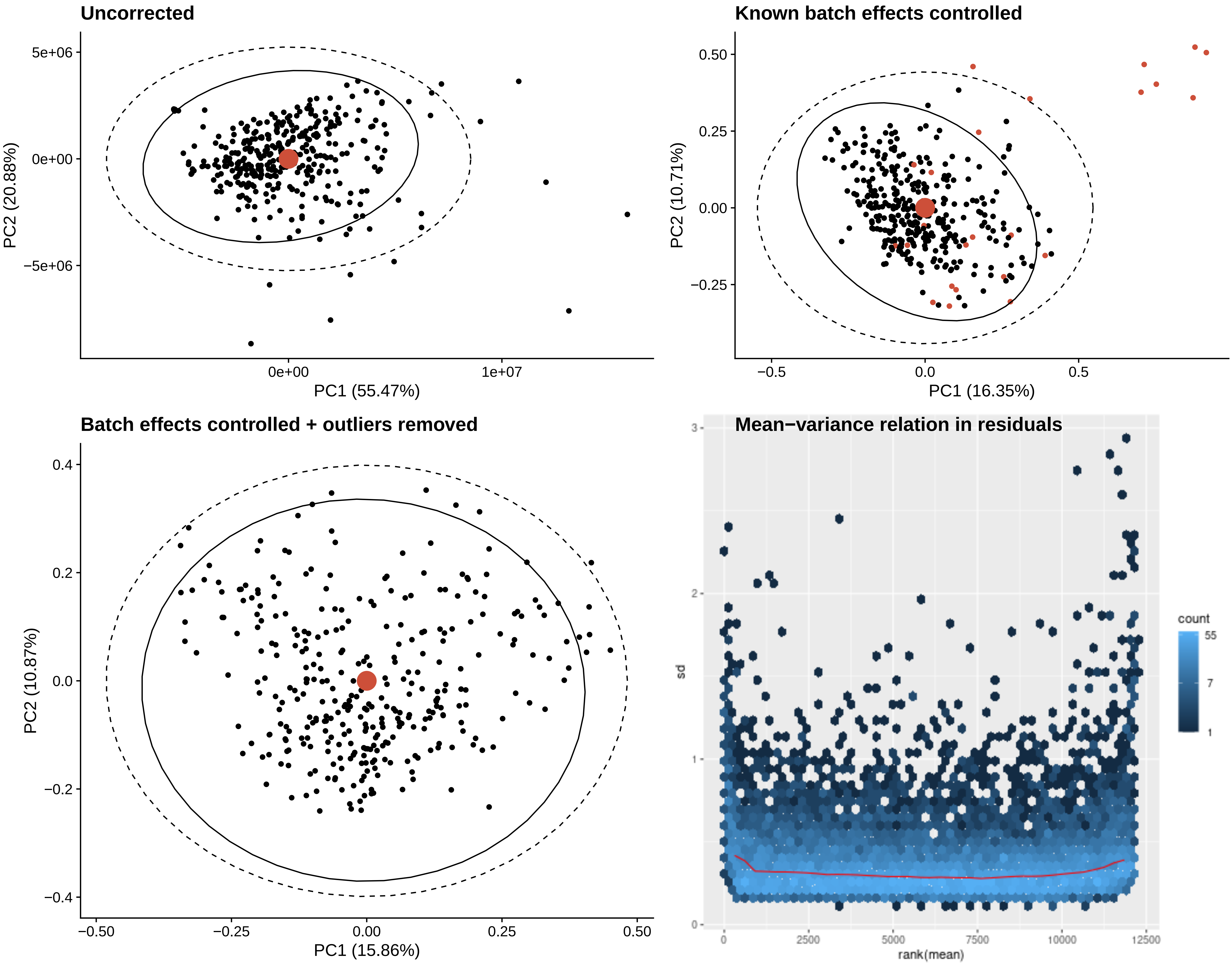
Uncorrected
5e+06
0e+00
−5e+06
0e+00
1e+07
PC1 (55.47%)
PC2 (20.88%)
Known batch effects controlled
0.50
0.25
0.00
−0.25
−0.5
0.0
0.5
PC1 (16.35%)
PC2 (10.71%)
Batch effects controlled + outliers removed
0.4
0.2
0.0
−0.2
−0.4
−0.50
−0.25
0.00
0.25
0.50
PC1 (15.86%)
PC2 (10.87%)
Mean−variance relation in residuals
sd
rank(mean)
0
2500
5000
7500
10000
12500
count
55
7
1

PITUITARY

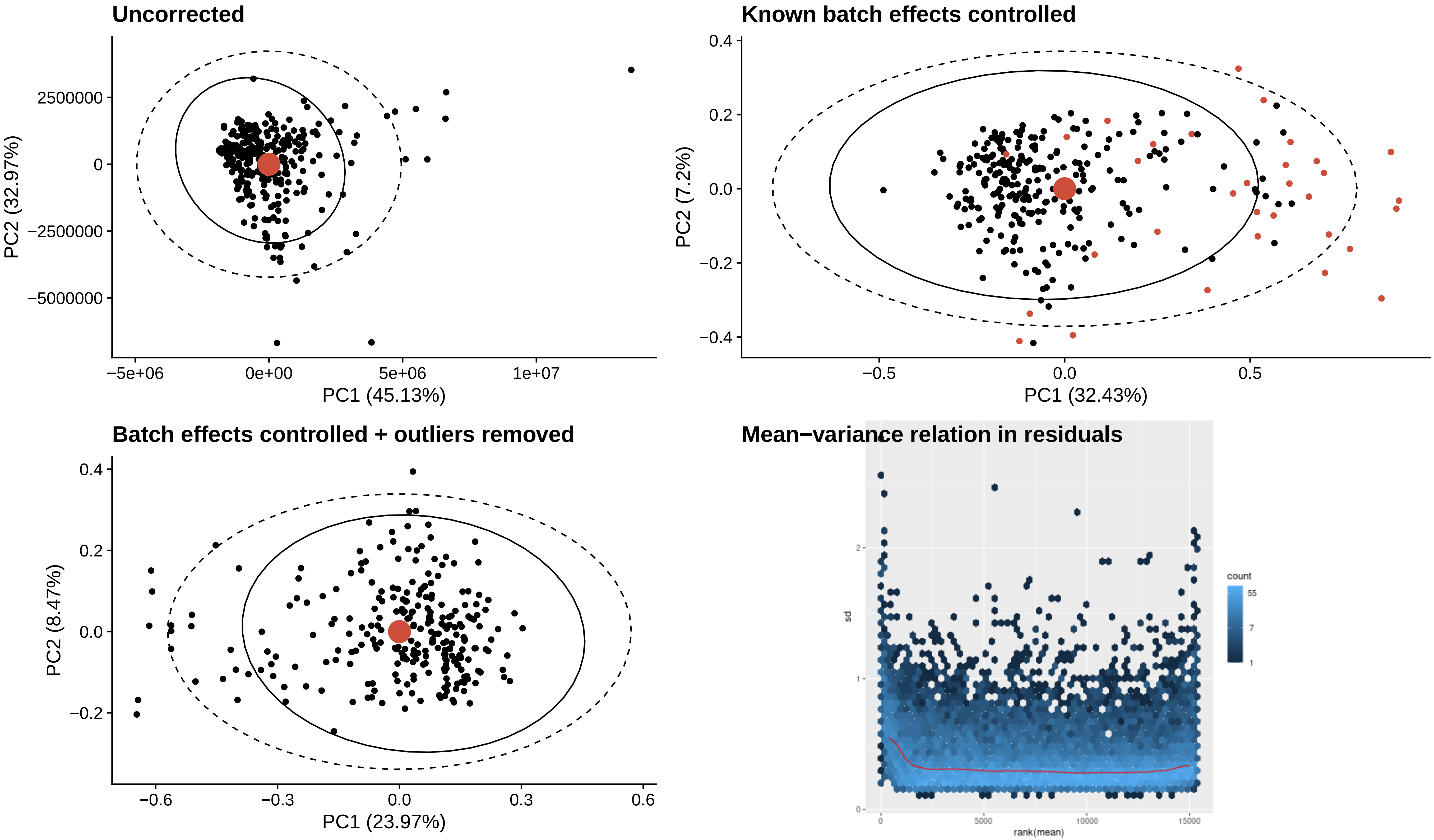
Uncorrected
PC2 (32.97%)
2500000
0
−2500000
−5000000
−5e+06
0e+00
5e+06
1e+07
PC1 (45.13%)
Known batch effects controlled
PC2 (7.2%)
0.4
0.2
0.0
−0.2
−0.4
−0.5
0.0
0.5
PC1 (32.43%)
Batch effects controlled + outliers removed
PC2 (8.47%)
0.4
0.2
0.0
−0.2
−0.6
−0.3
0.0
0.3
0.6
PC1 (23.97%)
Mean−variance relation in residuals
sd
2
1
0
0
5000
10000
15000
rank(mean)
count
55
7
1

PROSTATE

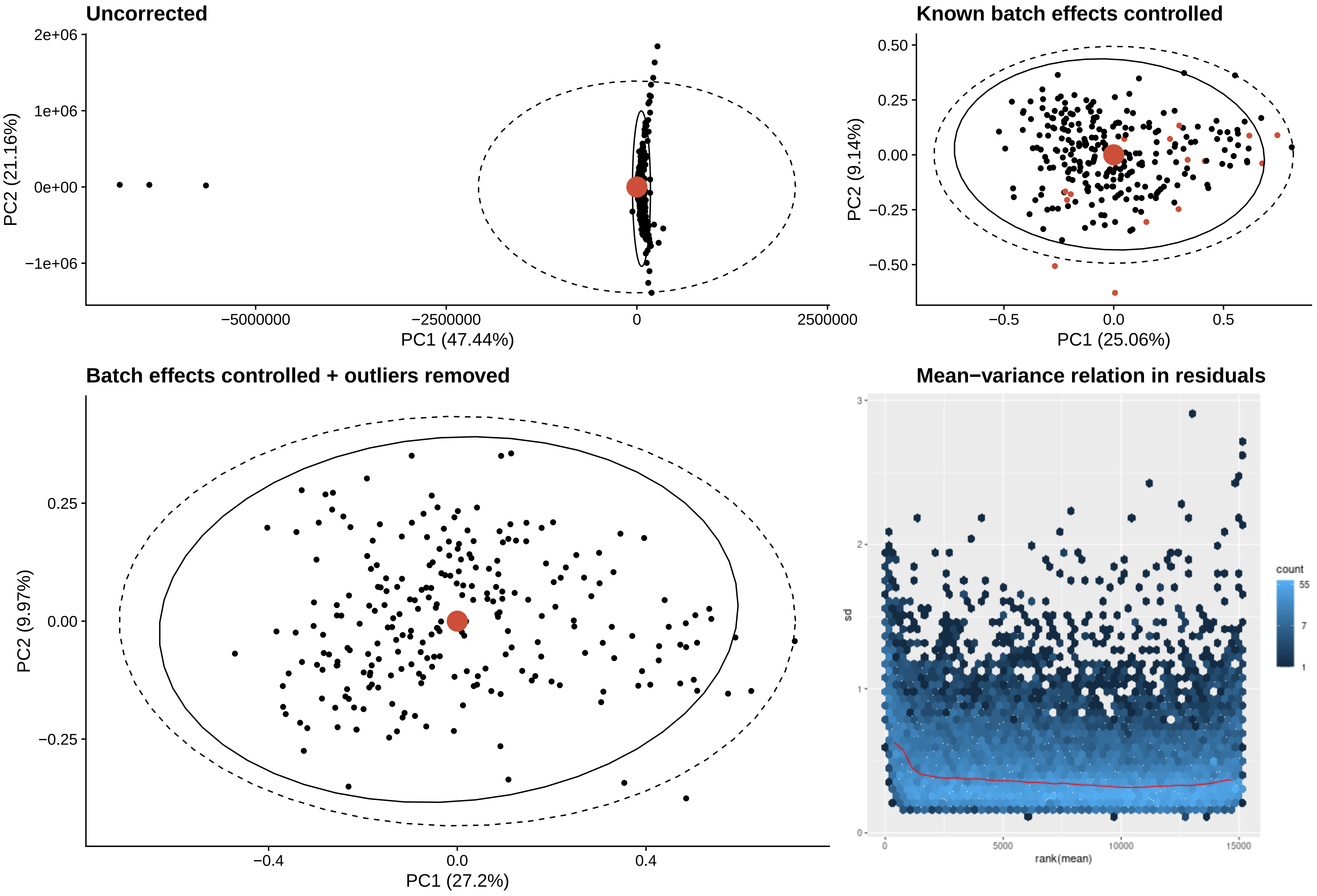
Uncorrected
2e+06
1e+06
0e+00
−1e+06
PC2 (21.16%)
−5000000
−2500000
0
2500000
PC1 (47.44%)
Known batch effects controlled
0.50
0.25
0.00
−0.25
−0.50
PC2 (9.14%)
−0.5
0.0
0.5
PC1 (25.06%)
Batch effects controlled + outliers removed
0.25
0.00
−0.25
PC2 (9.97%)
−0.4
0.0
0.4
PC1 (27.2%)
Mean−variance relation in residuals
sd
0
1
2
3
0
5000
10000
15000
rank(mean)
count
55
7
1

SALIVARY_GLAND

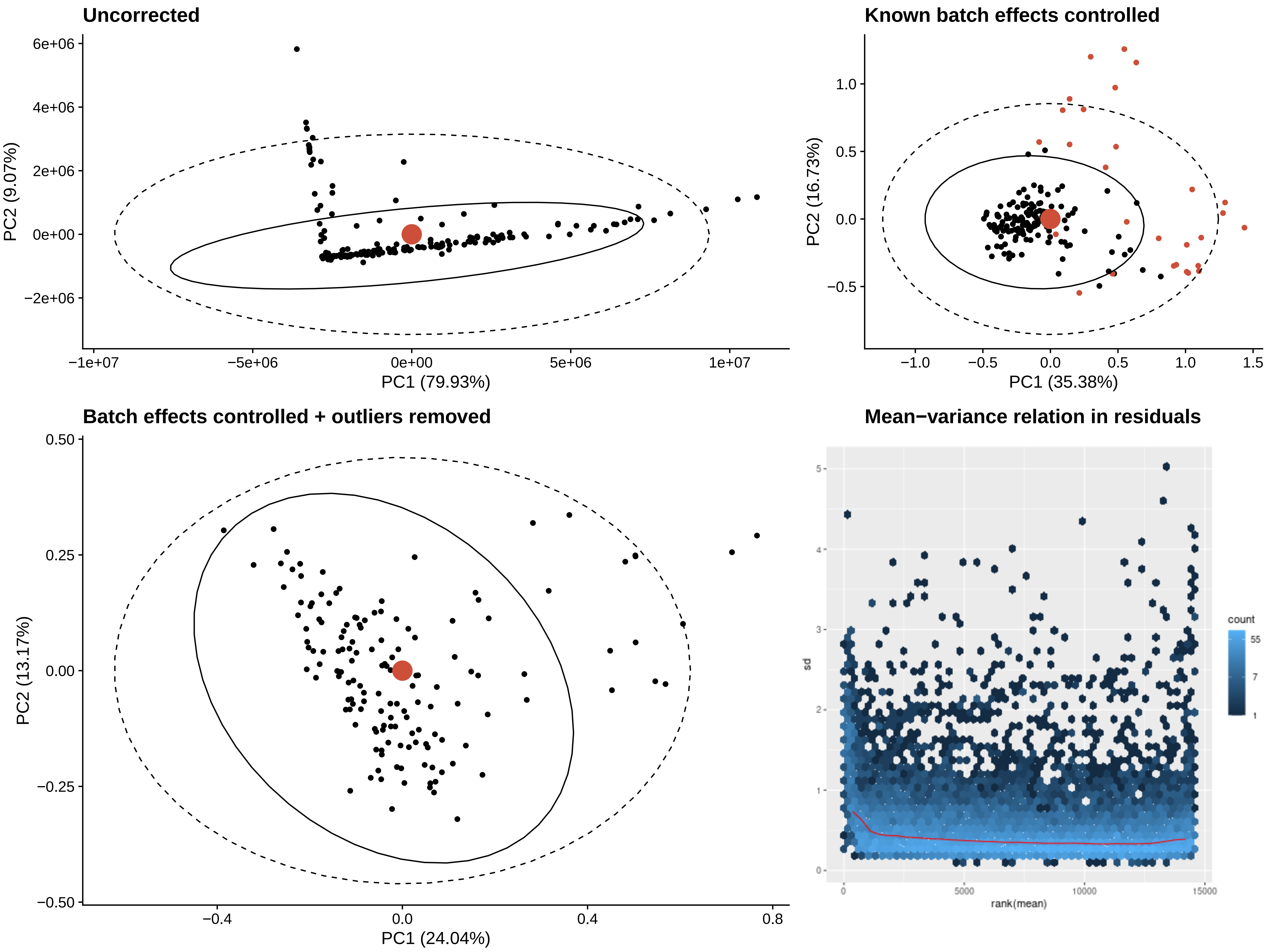

SKIN

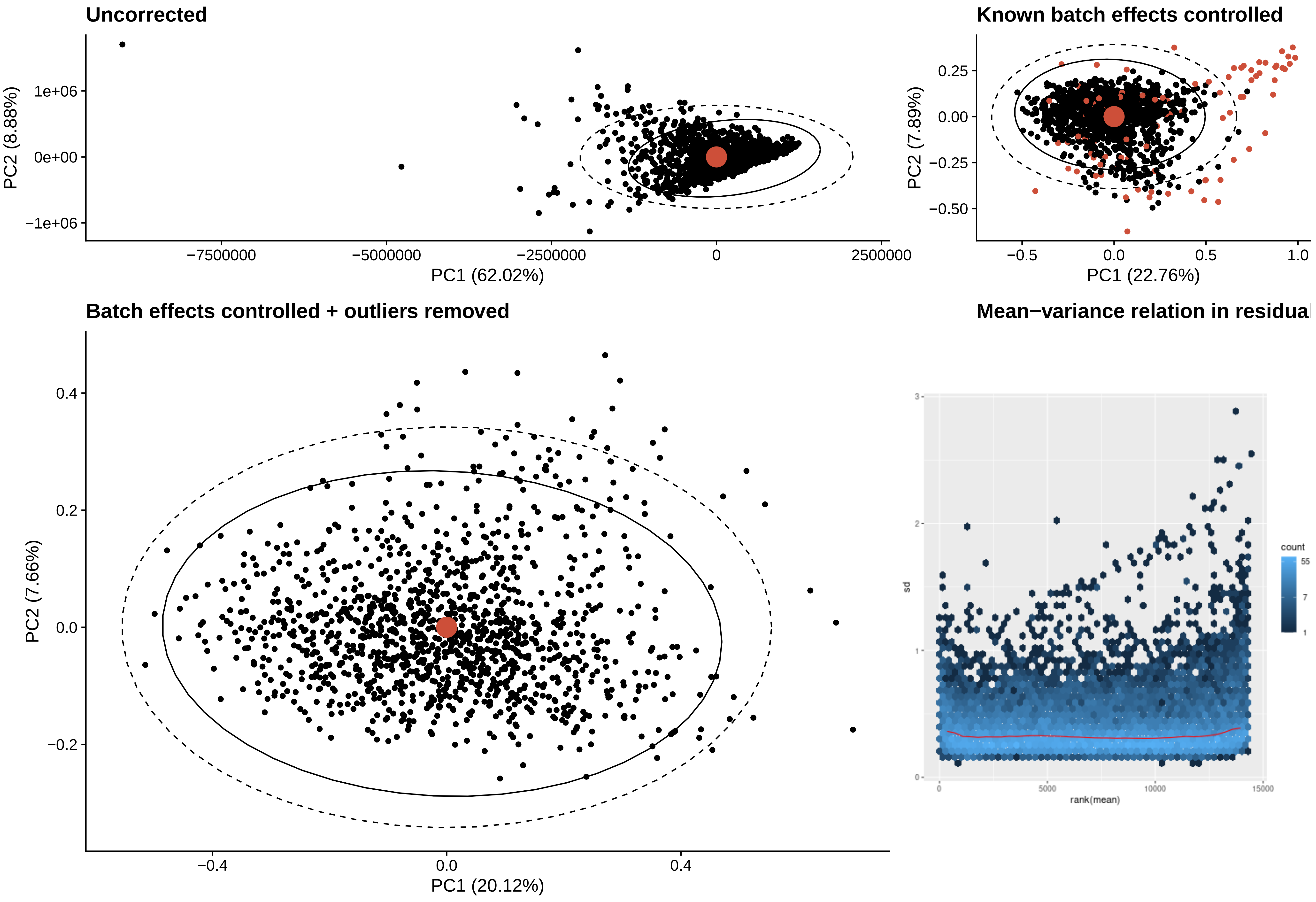

Uncorrected
PC2 (8.88%)
PC1 (62.02%)
Known batch effects controlled
PC2 (7.89%)
PC1 (22.76%)
Batch effects controlled + outliers removed
PC2 (7.66%)
PC1 (20.12%)
Mean−variance relation in residual
sd
rank(mean)
count

SPLEEN

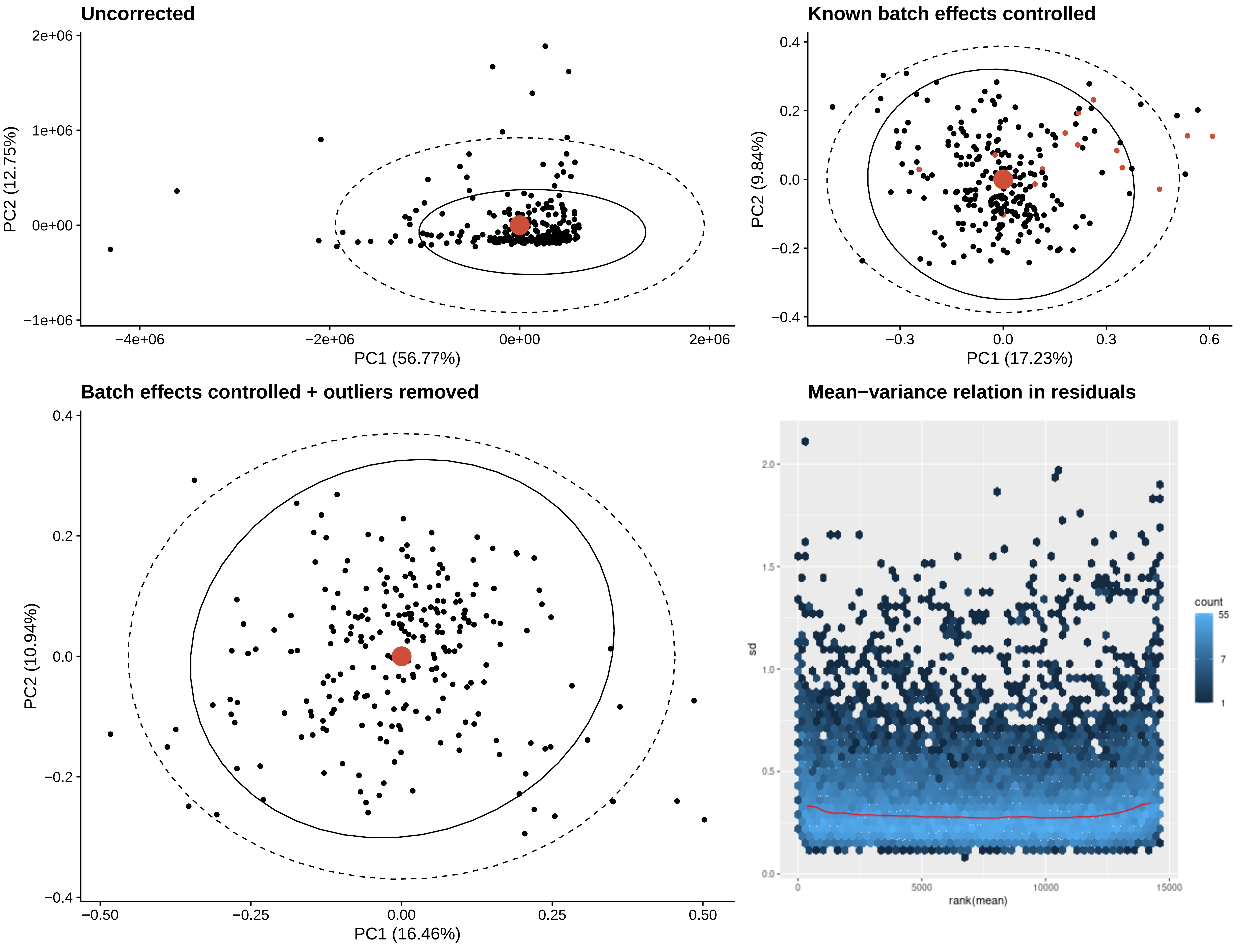

Uncorrected
2e+06
1e+06
0e+00
−1e+06
PC2 (12.75%)
−4e+06
−2e+06
0e+00
2e+06
PC1 (56.77%)
Known batch effects controlled
0.4
0.2
0.0
−0.2
−0.4
PC2 (9.84%)
−0.3
0.0
0.3
0.6
PC1 (17.23%)
Batch effects controlled + outliers removed
0.4
0.2
0.0
−0.2
−0.4
PC2 (10.94%)
−0.50
−0.25
0.00
0.25
0.50
PC1 (16.46%)
Mean−variance relation in residuals
2.0
1.5
1.0
0.5
0.0
sd
0
5000
10000
15000
rank(mean)
count
55
7
1

STOMACH

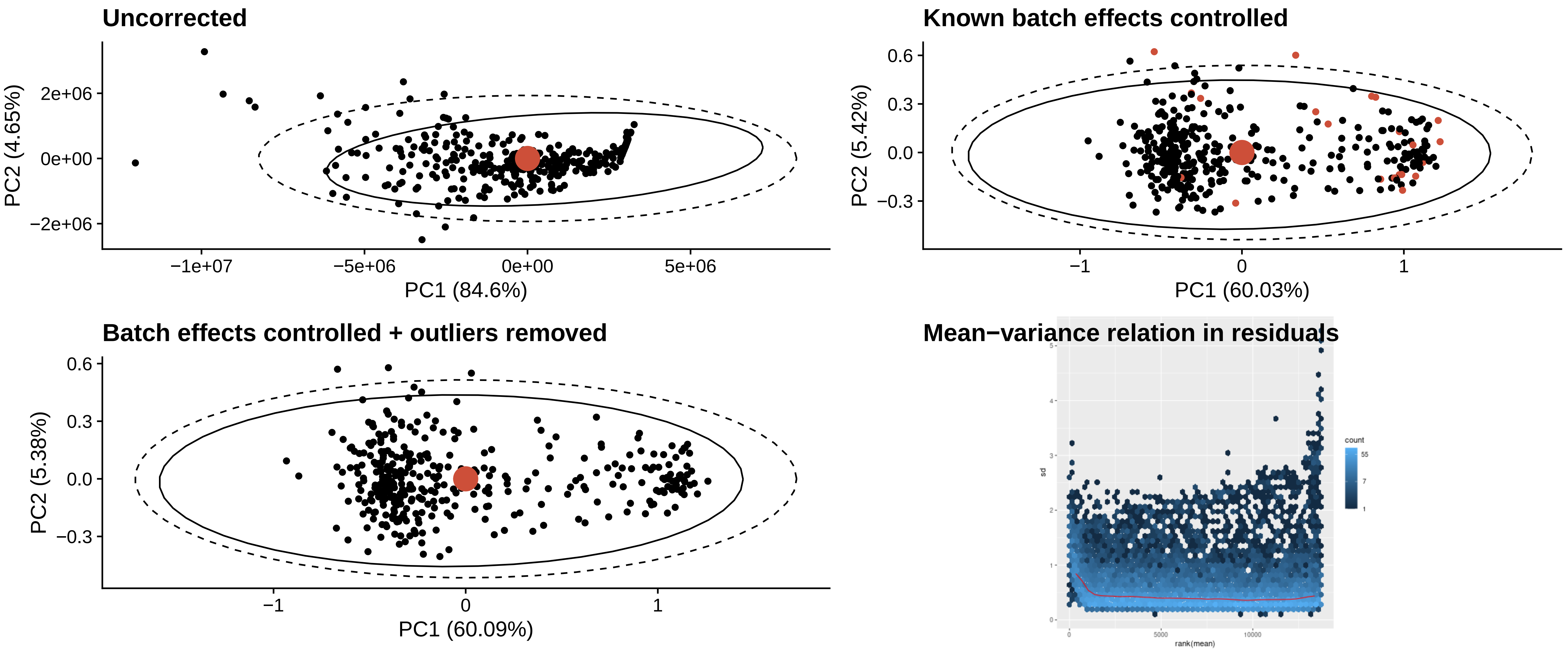
Uncorrected
PC2 (4.65%)
2e+06
0e+00
−2e+06
−1e+07
−5e+06
0e+00
5e+06
PC1 (84.6%)
Known batch effects controlled
PC2 (5.42%)
0.6
0.3
0.0
−0.3
−1
0
1
PC1 (60.03%)
Batch effects controlled + outliers removed
PC2 (5.38%)
0.6
0.3
0.0
−0.3
−1
0
1
PC1 (60.09%)
Mean−variance relation in residuals
count
rank(mean)

TESTIS

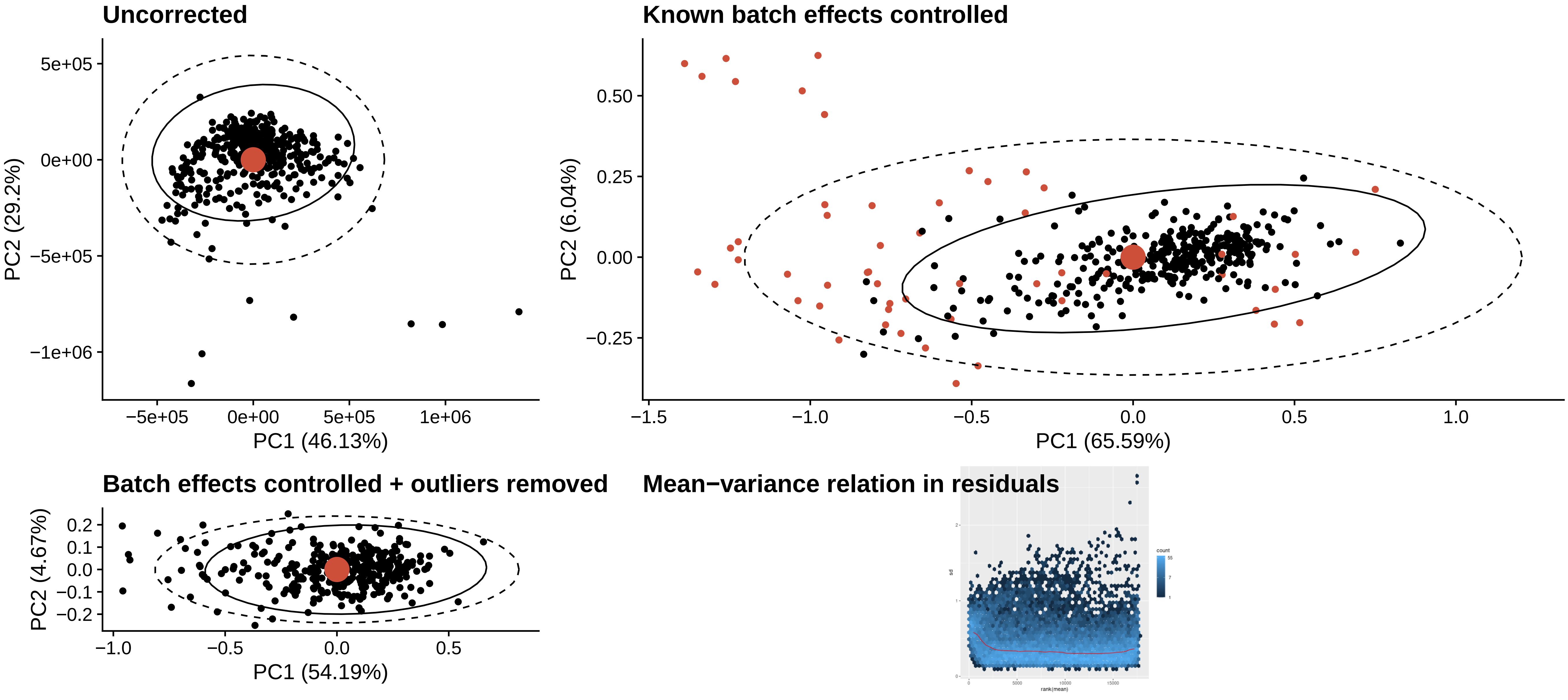

Uncorrected
5e+05
0e+00
−5e+05
−1e+06
PC2 (29.2%)
−5e+05
0e+00
5e+05
1e+06
PC1 (46.13%)
Known batch effects controlled
0.50
0.25
0.00
−0.25
PC2 (6.04%)
−1.5
−1.0
−0.5
0.0
0.5
1.0
PC1 (65.59%)
Batch effects controlled + outliers removed
0.2
0.1
0.0
−0.1
−0.2
PC2 (4.67%)
−1.0
−0.5
0.0
0.5
PC1 (54.19%)
Mean−variance relation in residuals

THYROID

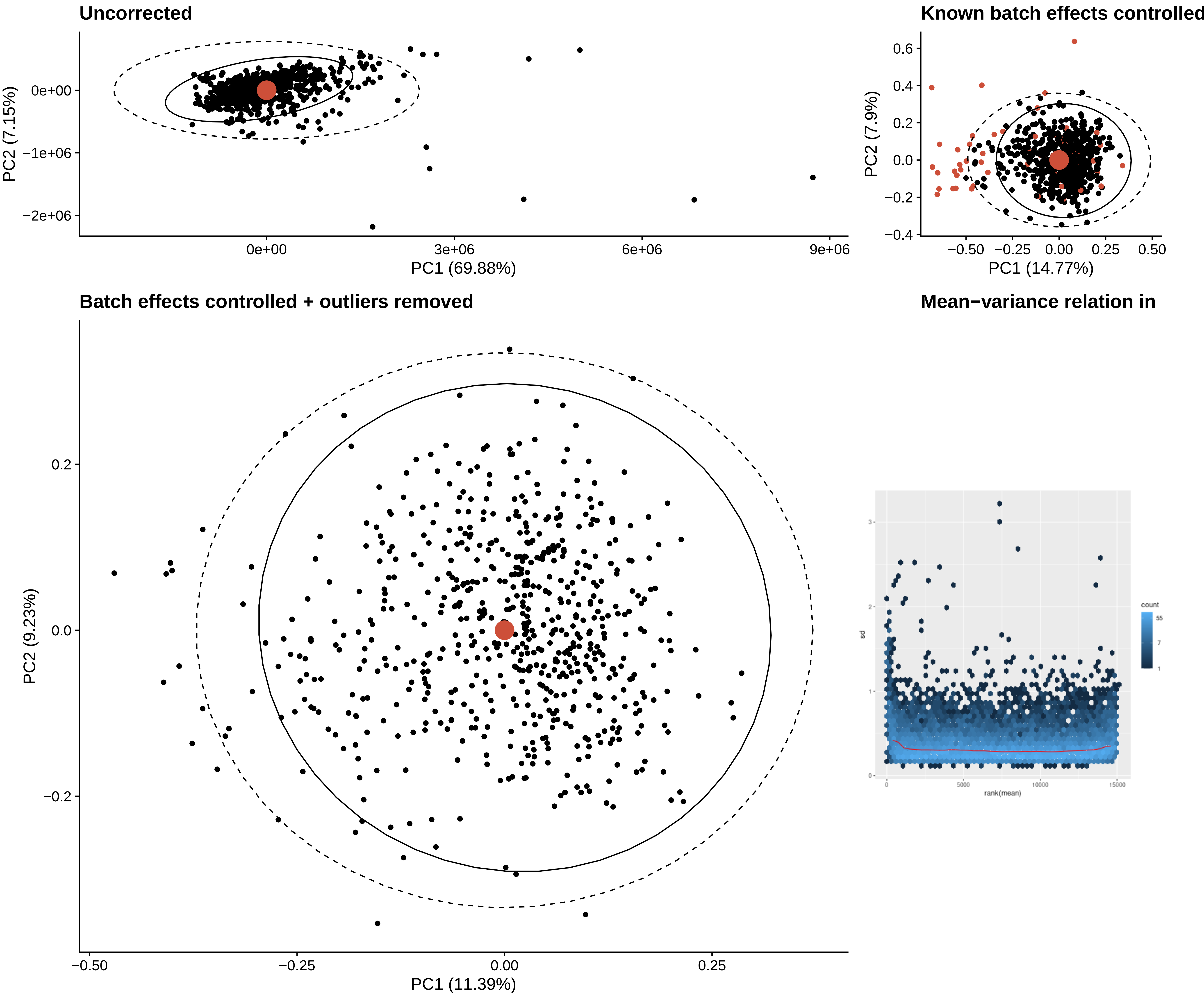

Uncorrected
PC2 (7.15%)
0e+00
−1e+06
−2e+06
0e+00
3e+06
6e+06
9e+06
PC1 (69.88%)
Known batch effects controlled
PC2 (7.9%)
0.6
0.4
0.2
0.0
−0.2
−0.4
−0.50
−0.25
0.00
0.25
0.50
PC1 (14.77%)
Batch effects controlled + outliers removed
PC2 (9.23%)
0.2
0.0
−0.2
−0.50
−0.25
0.00
0.25
PC1 (11.39%)
Mean−variance relation in
count
55
7
1
sd
rank(mean)
0
5000
10000
15000

BRCA

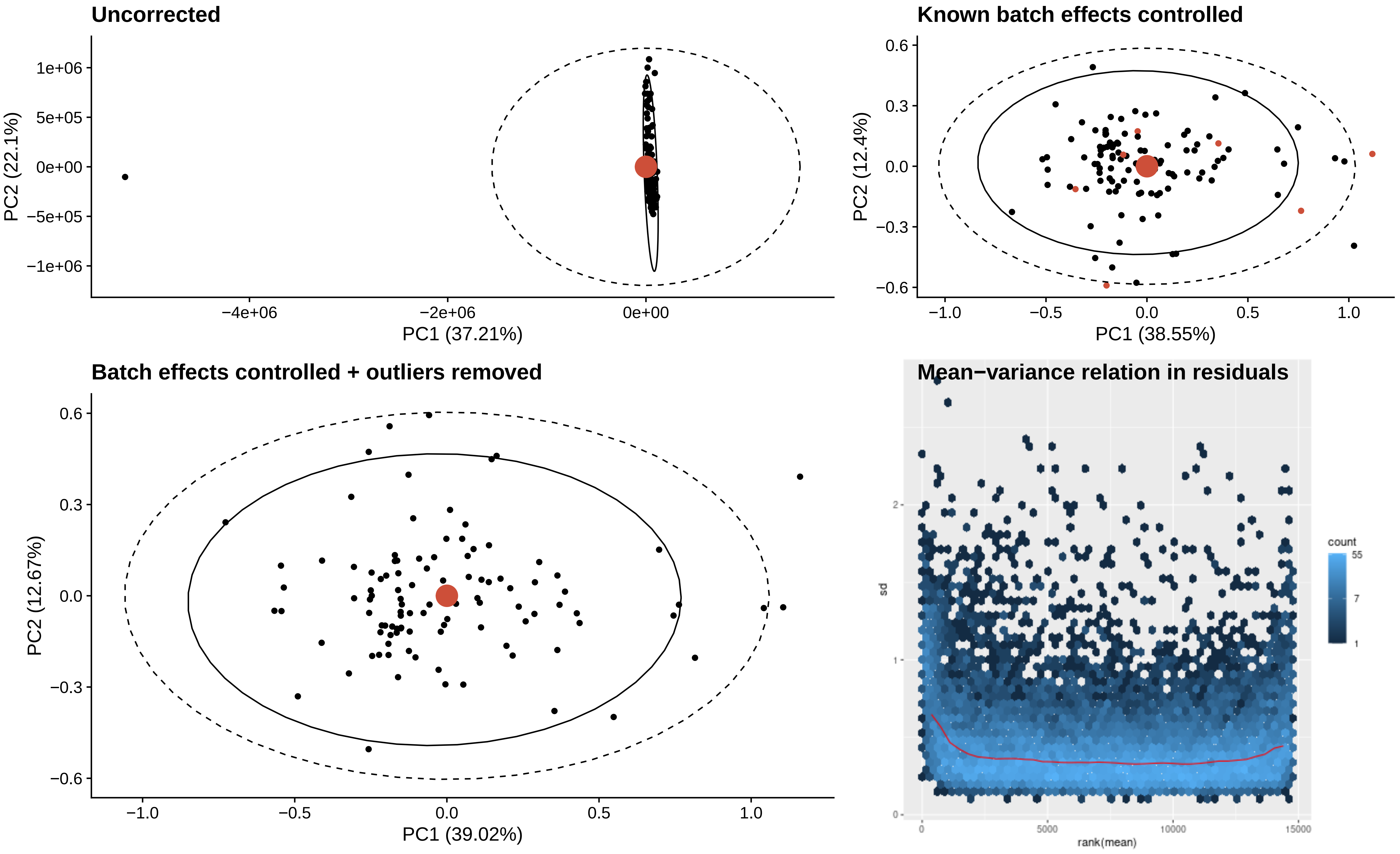
Uncorrected
PC2 (22.1%)
1e+06
5e+05
0e+00
−5e+05
−1e+06
−4e+06
−2e+06
0e+00
PC1 (37.21%)
Known batch effects controlled
PC2 (12.4%)
0.6
0.3
0.0
−0.3
−0.6
−1.0
−0.5
0.0
0.5
1.0
PC1 (38.55%)
Batch effects controlled + outliers removed
PC2 (12.67%)
0.6
0.3
0.0
−0.3
−0.6
−1.0
−0.5
0.0
0.5
1.0
PC1 (39.02%)
Mean−variance relation in residuals
sd
2
1
0
0
5000
10000
15000
rank(mean)
count
55
7
1

COAD

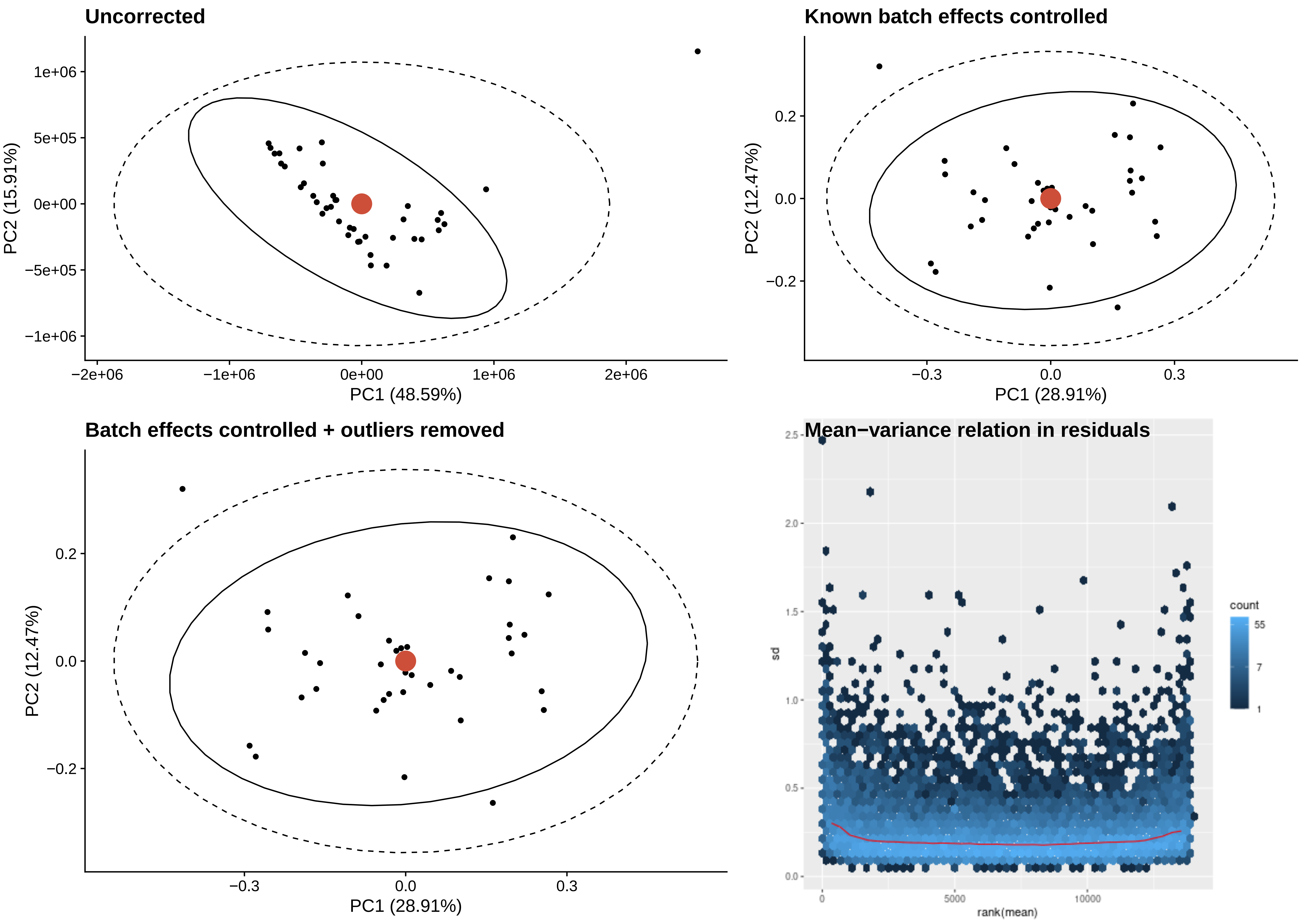

Uncorrected
1e+06
5e+05
0e+00
−5e+05
−1e+06
PC2 (15.91%)
−2e+06
−1e+06
0e+00
1e+06
2e+06
PC1 (48.59%)
Known batch effects controlled
0.2
0.0
−0.2
PC2 (12.47%)
−0.3
0.0
0.3
PC1 (28.91%)
Batch effects controlled + outliers removed
0.2
0.0
−0.2
PC2 (12.47%)
−0.3
0.0
0.3
PC1 (28.91%)
Mean−variance relation in residuals
2.5
2.0
1.5
1.0
0.5
0.0
sd
0
5000
10000
rank(mean)
count
55
7
1

KIRC

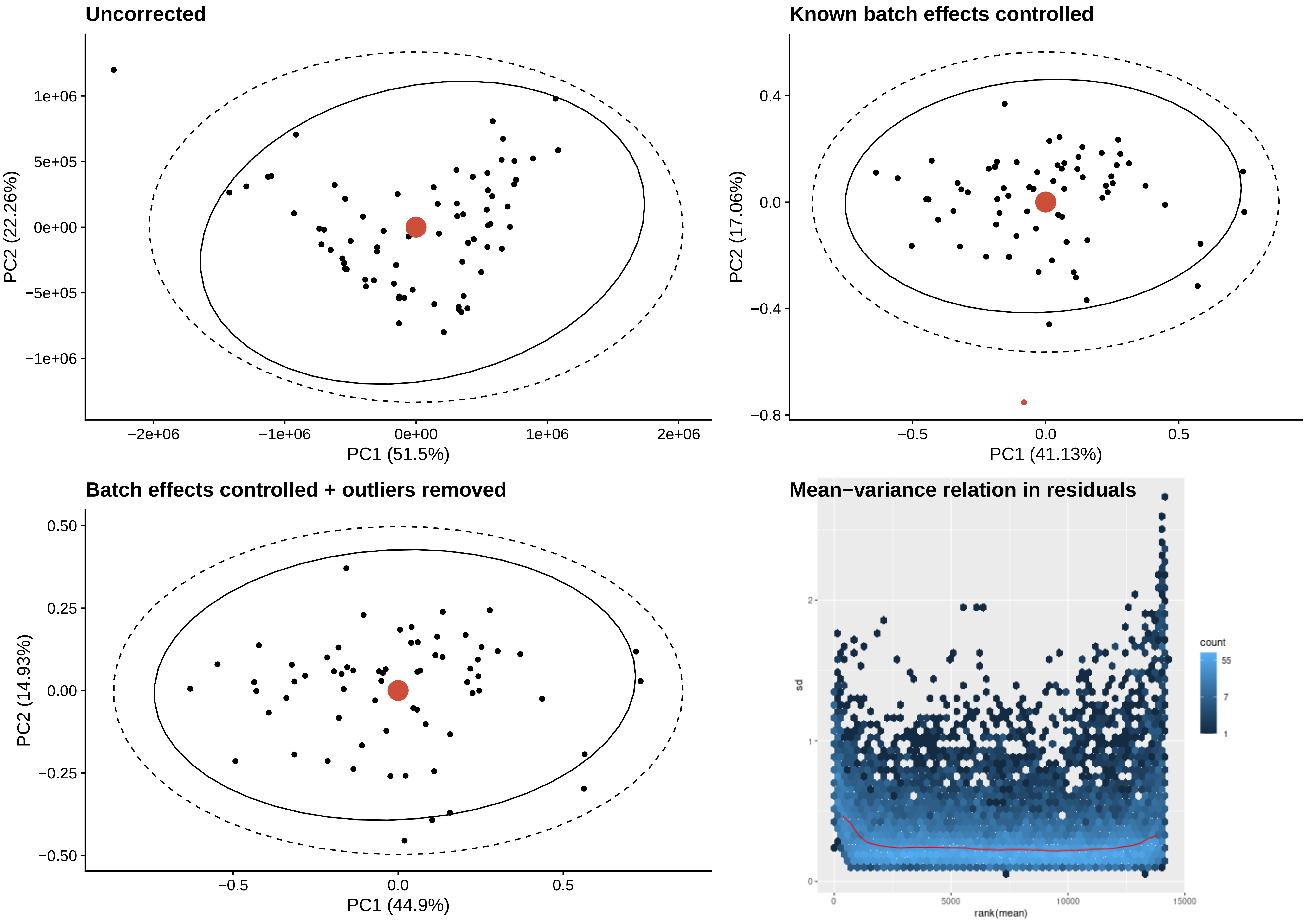

Uncorrected
1e+06
5e+05
0e+00
−5e+05
−1e+06
PC2 (22.26%)
−2e+06
−1e+06
0e+00
1e+06
2e+06
PC1 (51.5%)
Known batch effects controlled
0.4
0.0
−0.4
−0.8
PC2 (17.06%)
−0.5
0.0
0.5
PC1 (41.13%)
Batch effects controlled + outliers removed
0.50
0.25
0.00
−0.25
−0.50
PC2 (14.93%)
−0.5
0.0
0.5
PC1 (44.9%)
Mean−variance relation in residuals
sd
2
1
0
0
5000
10000
15000
rank(mean)
count
55
7
1

KIRP

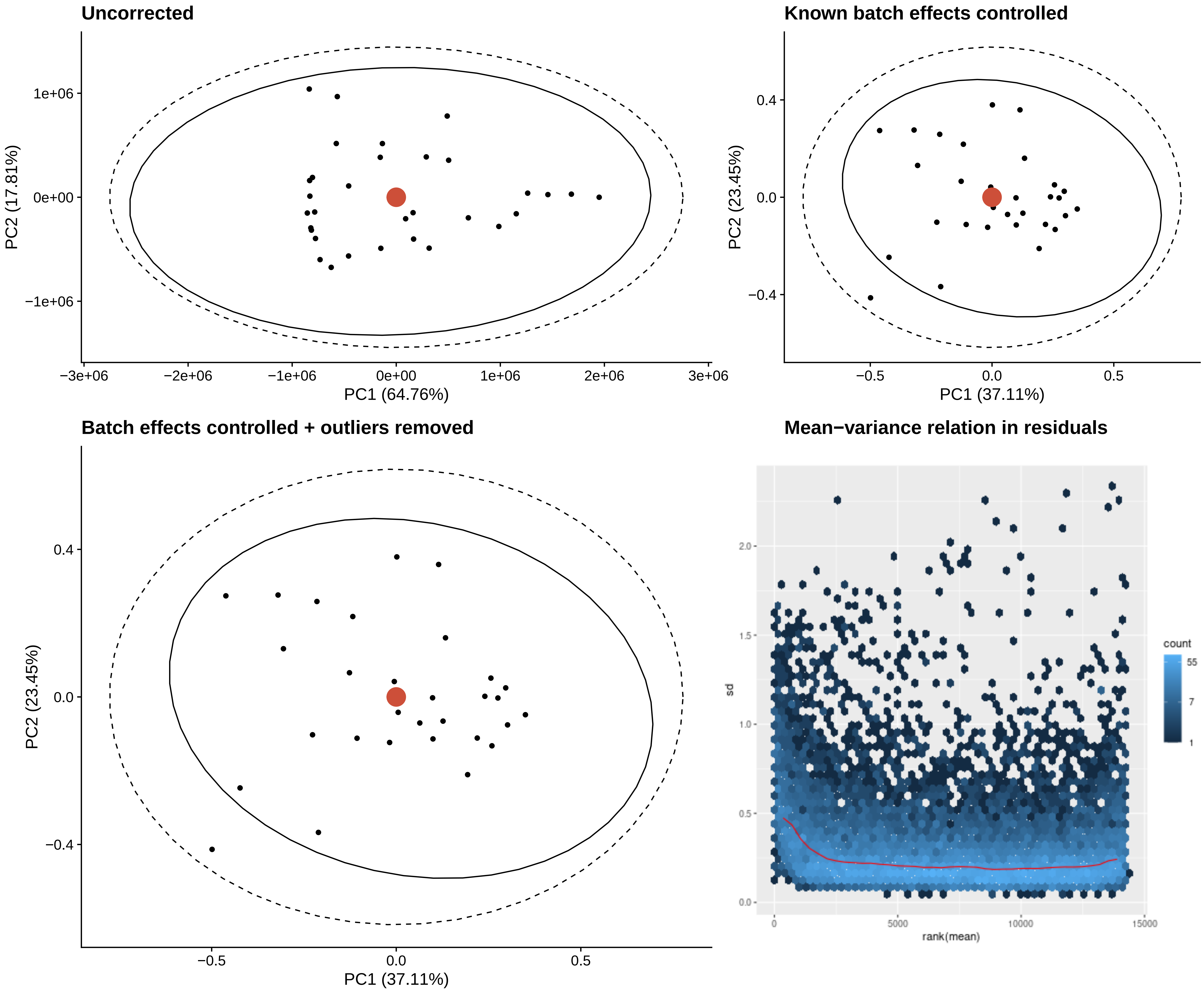

Uncorrected
1e+06
0e+00
−1e+06
PC2 (17.81%)
−3e+06
−2e+06
−1e+06
0e+00
1e+06
2e+06
3e+06
PC1 (64.76%)
Known batch effects controlled
0.4
0.0
−0.4
PC2 (23.45%)
−0.5
0.0
0.5
PC1 (37.11%)
Batch effects controlled + outliers removed
0.4
0.0
−0.4
PC2 (23.45%)
−0.5
0.0
0.5
PC1 (37.11%)
Mean−variance relation in residuals
sd
rank(mean)
0
5000
10000
15000
0.0
0.5
1.0
1.5
2.0
count
55
7
1

LIHC

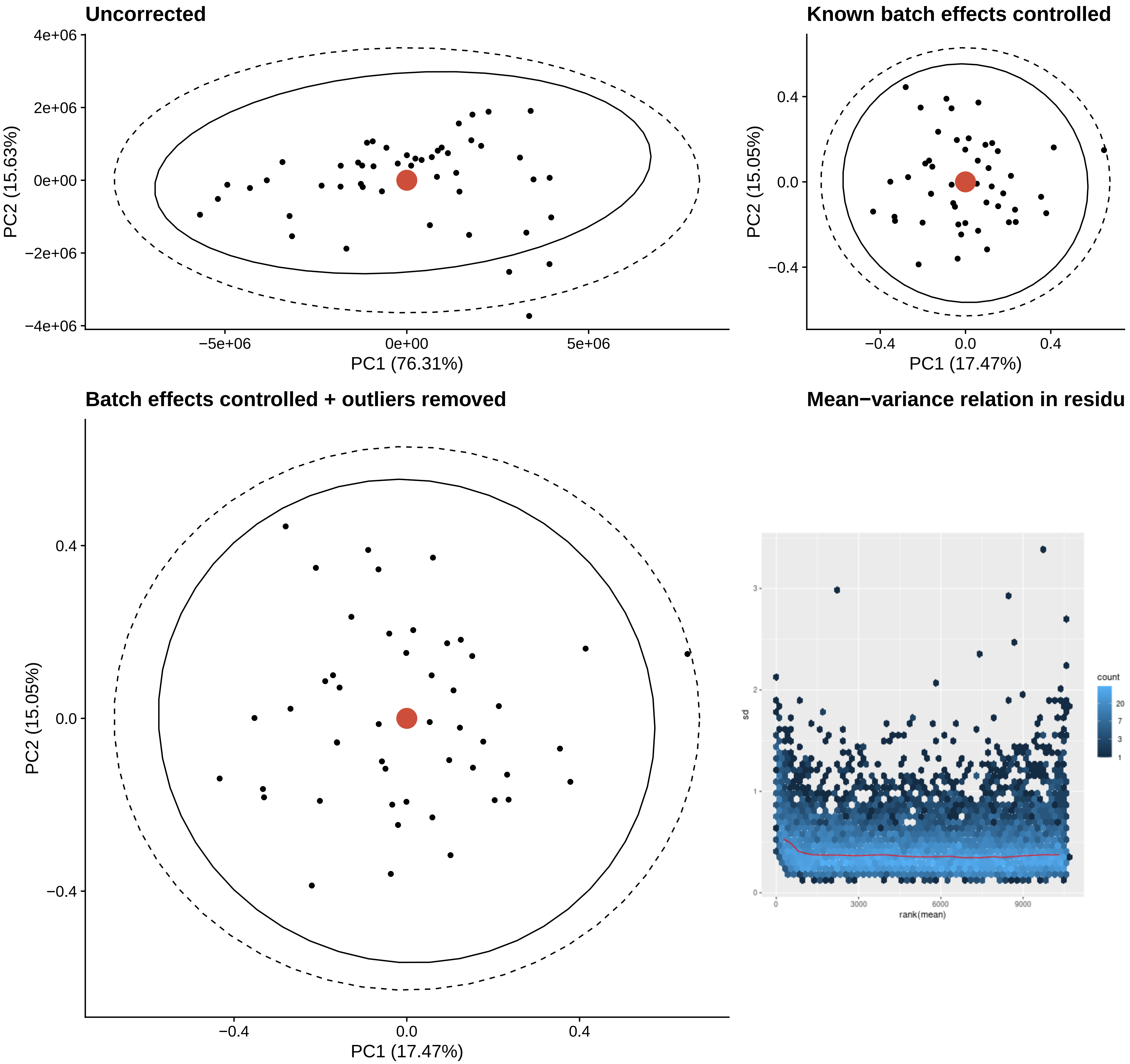
Uncorrected
4e+06
2e+06
0e+00
−2e+06
−4e+06
PC2 (15.63%)
−5e+06
0e+00
5e+06
PC1 (76.31%)
Known batch effects controlled
0.4
0.0
−0.4
PC2 (15.05%)
−0.4
0.0
0.4
PC1 (17.47%)
Batch effects controlled + outliers removed
0.4
0.0
−0.4
PC2 (15.05%)
−0.4
0.0
0.4
PC1 (17.47%)
Mean−variance relation in residu
sd
3
2
1
0
0
3000
6000
9000
rank(mean)
count
20
7
3
1

LUAD

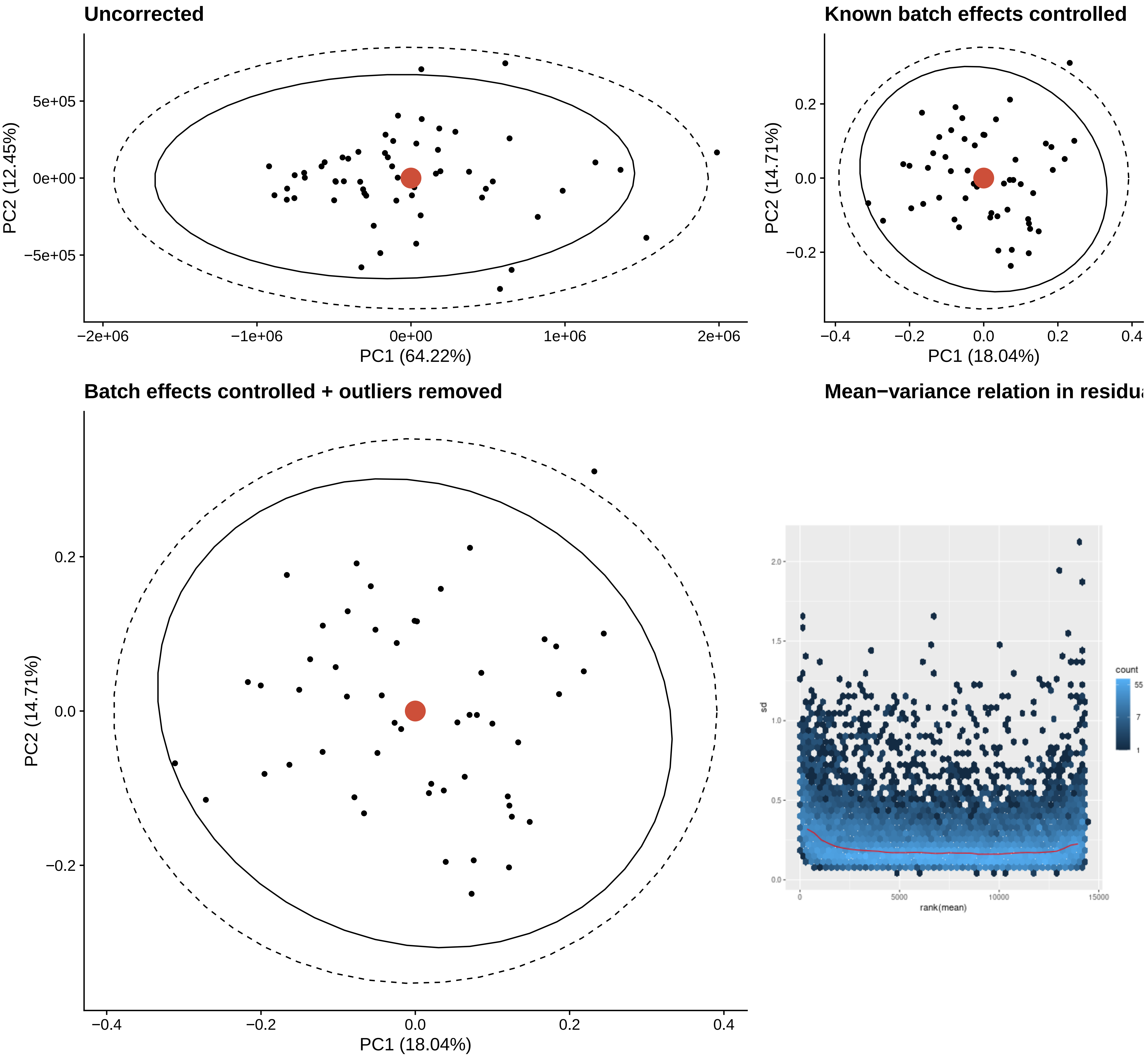

Uncorrected
PC2 (12.45%)
PC1 (64.22%)
5e+05
0e+00
−5e+05
−2e+06
−1e+06
0e+00
1e+06
2e+06
Known batch effects controlled
PC2 (14.71%)
PC1 (18.04%)
Batch effects controlled + outliers removed
PC2 (14.71%)
PC1 (18.04%)
0.2
0.0
−0.2
−0.4
0.4
Mean−variance relation in residu
count
sd
rank(mean)

LUSC

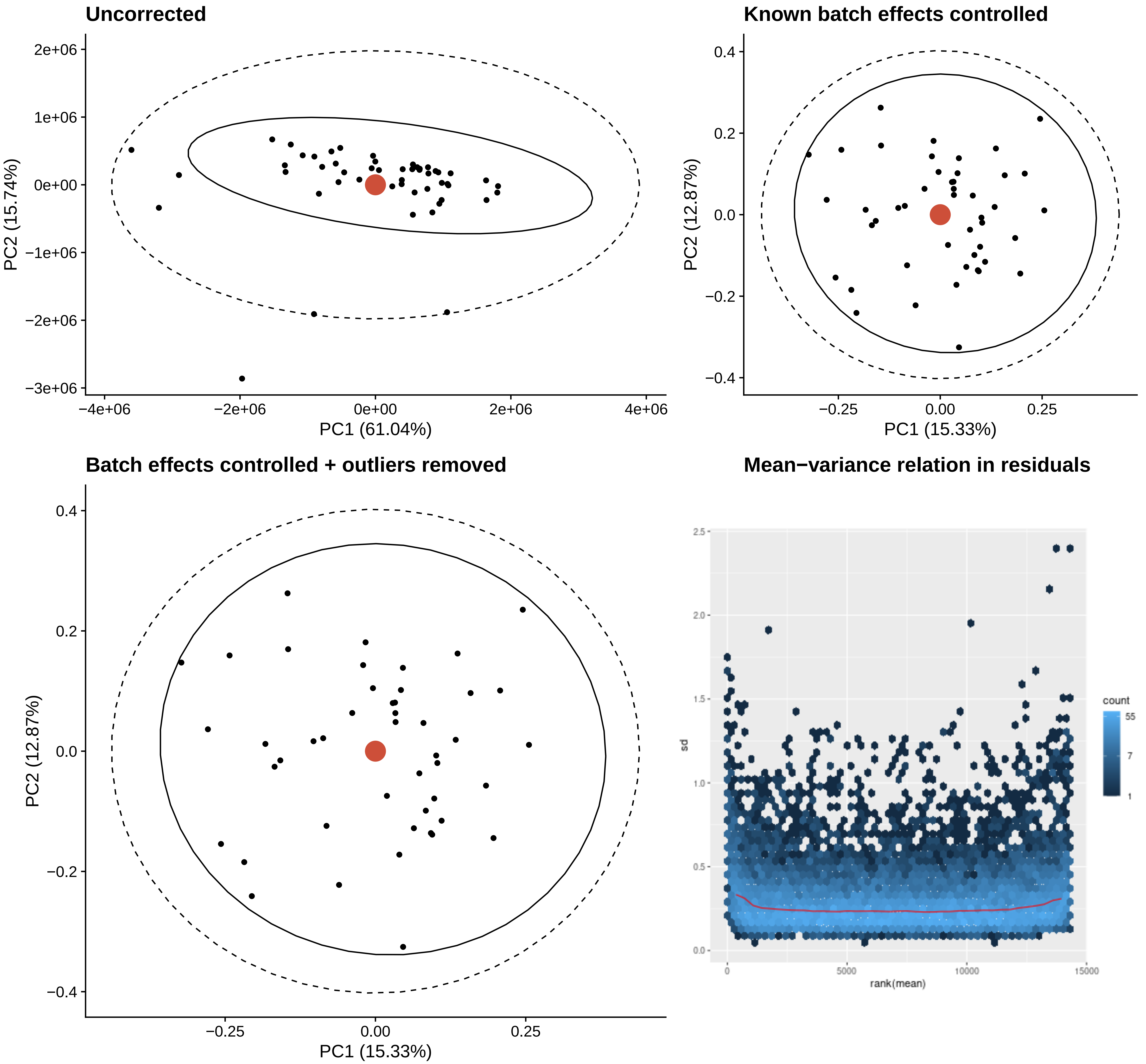

Uncorrected
2e+06
1e+06
0e+00
−1e+06
−2e+06
−3e+06
PC2 (15.74%)
−4e+06
−2e+06
0e+00
2e+06
4e+06
PC1 (61.04%)
Known batch effects controlled
0.4
0.2
0.0
−0.2
−0.4
PC2 (12.87%)
−0.25
0.00
0.25
PC1 (15.33%)
Batch effects controlled + outliers removed
0.4
0.2
0.0
−0.2
−0.4
PC2 (12.87%)
−0.25
0.00
0.25
PC1 (15.33%)
Mean−variance relation in residuals
2.5
2.0
1.5
1.0
0.5
0.0
sd
0
5000
10000
15000
rank(mean)
count
55
7
1

PRAD

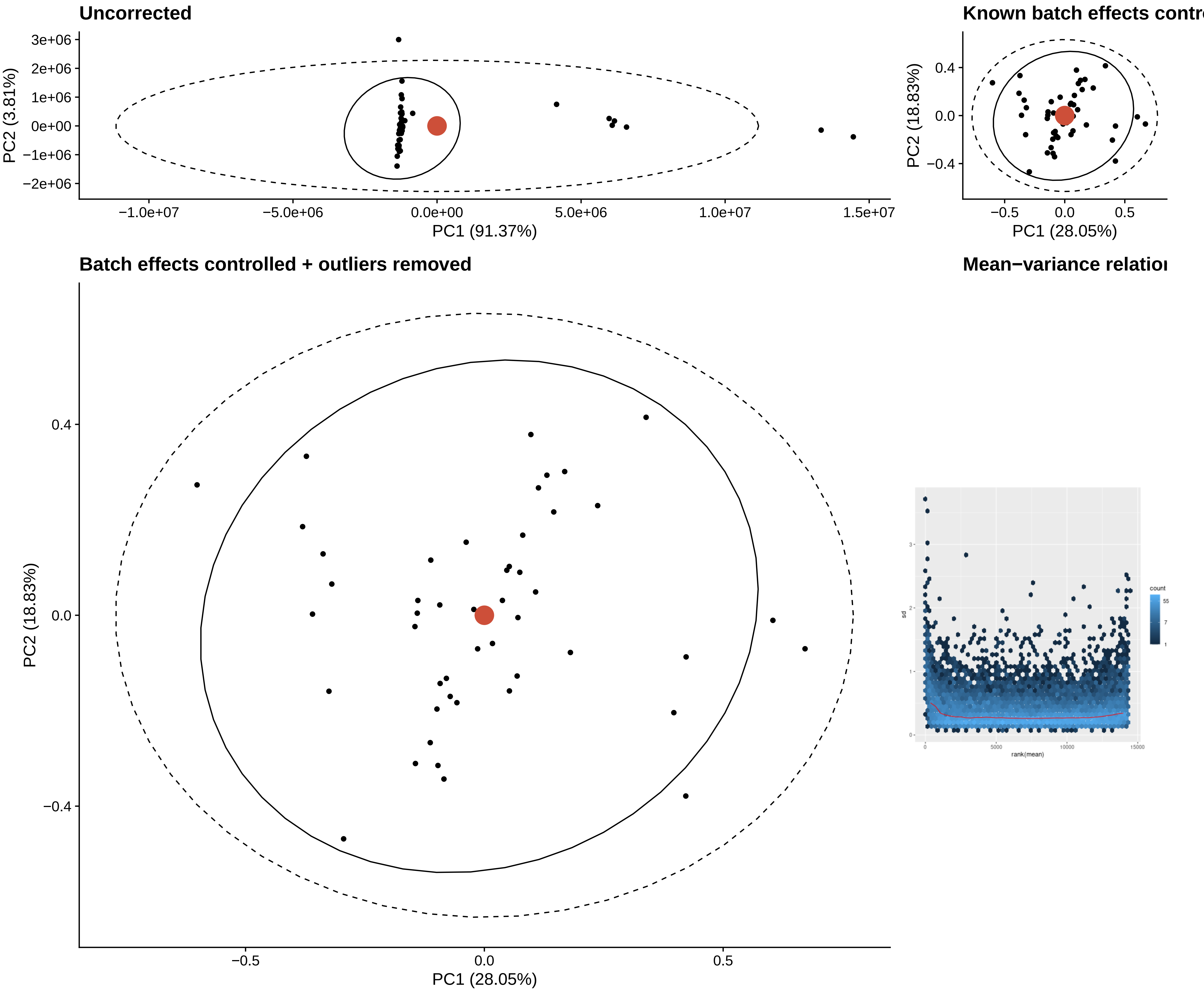
Uncorrected
PC2 (3.81%)
3e+06
2e+06
1e+06
0e+00
−1e+06
−2e+06
−1.0e+07
−5.0e+06
0.0e+00
5.0e+06
1.0e+07
1.5e+07
PC1 (91.37%)
Known batch effects contr
PC2 (18.83%)
0.4
0.0
−0.4
−0.5
0.0
0.5
PC1 (28.05%)
Batch effects controlled + outliers removed
PC2 (18.83%)
0.4
0.0
−0.4
−0.5
0.0
0.5
PC1 (28.05%)
Mean−variance relatio
sd
rank(mean)
count

STAD

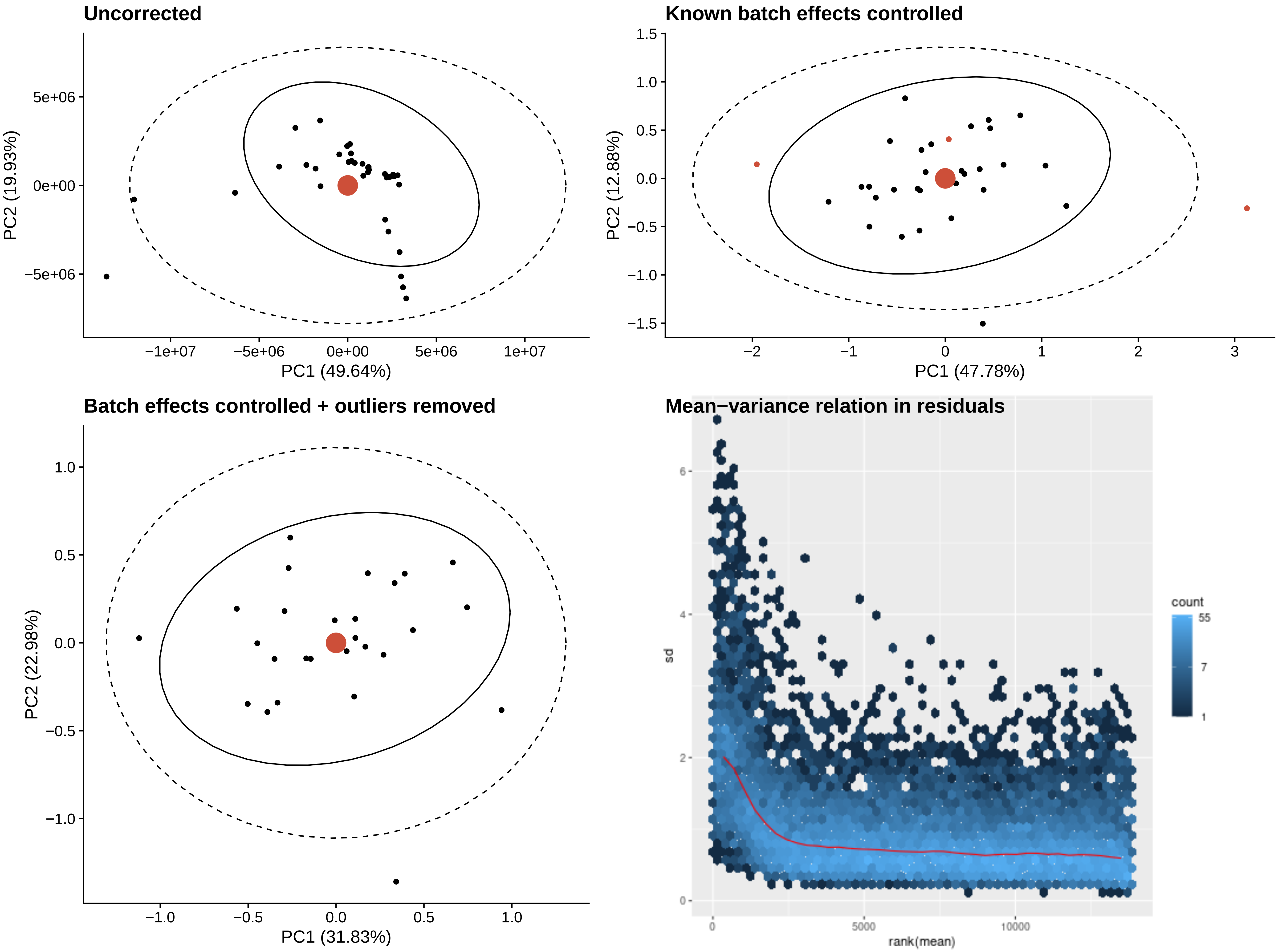
Uncorrected
5e+06
0e+00
−5e+06
−1e+07
−5e+06
0e+00
5e+06
1e+07
PC1 (49.64%)
PC2 (19.93%)
Known batch effects controlled
1.5
1.0
0.5
0.0
−0.5
−1.0
−1.5
−2
−1
0
1
2
3
PC1 (47.78%)
PC2 (12.88%)
Batch effects controlled + outliers removed
1.0
0.5
0.0
−0.5
−1.0
−1.0
−0.5
0.0
0.5
1.0
PC1 (31.83%)
PC2 (22.98%)
Mean−variance relation in residuals
6
4
2
0
0
5000
10000
rank(mean)
sd
count
55
7
1

THCA

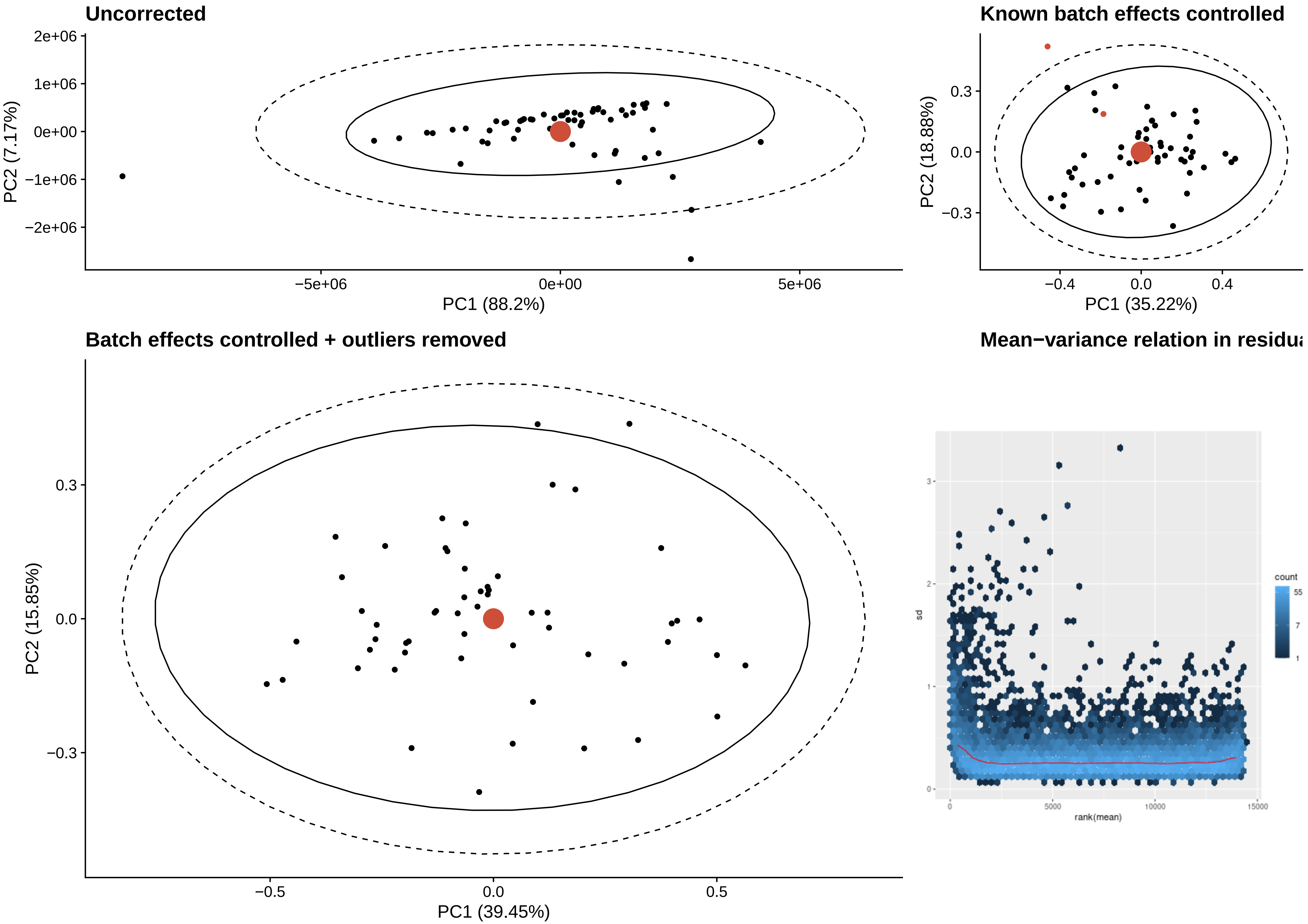

Uncorrected
2e+06
1e+06
0e+00
−1e+06
−2e+06
PC2 (7.17%)
−5e+06
0e+00
5e+06
PC1 (88.2%)
Known batch effects controlled
0.3
0.0
−0.3
PC2 (18.88%)
−0.4
0.0
0.4
PC1 (35.22%)
Batch effects controlled + outliers removed
0.3
0.0
−0.3
PC2 (15.85%)
−0.5
0.0
0.5
PC1 (39.45%)
Mean−variance relation in residua
sd
rank(mean)
count

UCEC

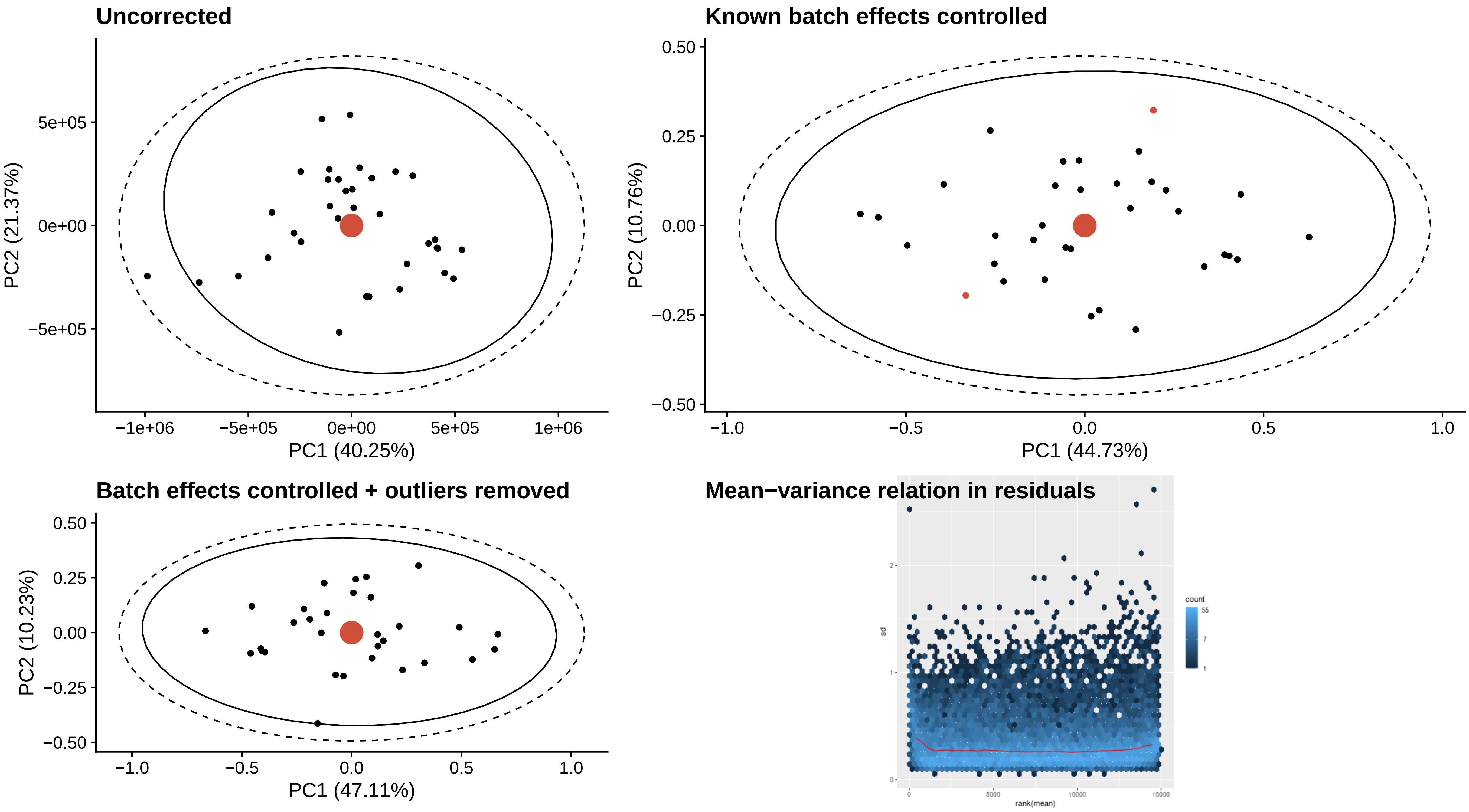

Uncorrected
PC2 (21.37%)
PC1 (40.25%)
5e+05
0e+00
−5e+05
−1e+06
−5e+05
0e+00
5e+05
1e+06
Known batch effects controlled
PC2 (10.76%)
PC1 (44.73%)
0.50
0.25
0.00
−0.25
−0.50
−1.0
−0.5
0.0
0.5
1.0
Batch effects controlled + outliers removed
PC2 (10.23%)
PC1 (47.11%)
0.50
0.25
0.00
−0.25
−0.50
−1.0
−0.5
0.0
0.5
1.0
Mean−variance relation in residuals
sd
rank(mean)
count
55
7
1
0
5000
10000
15000
2
1
0

E−ENAD−33

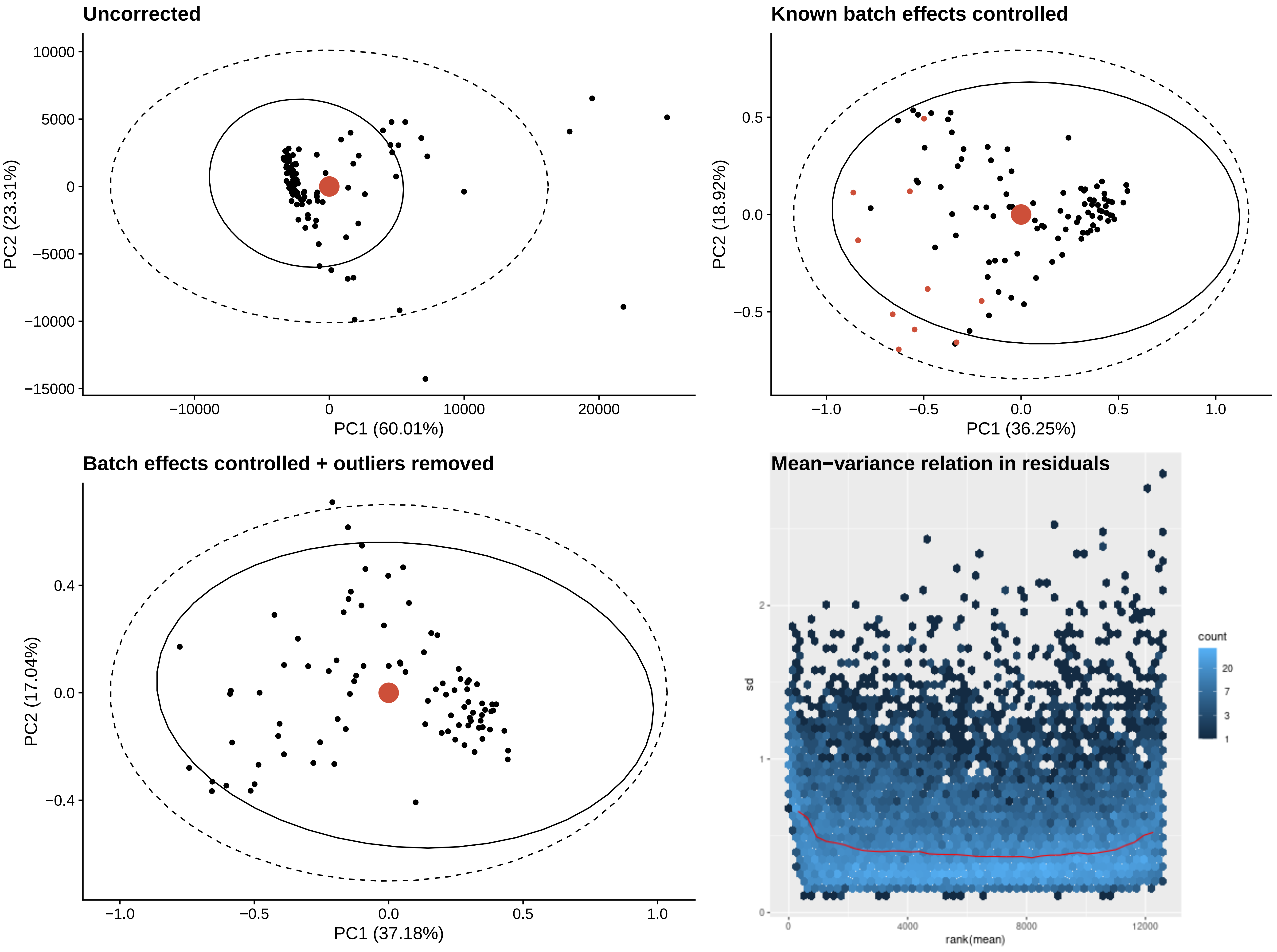
Uncorrected
10000
5000
0
−5000
−10000
−15000
PC2 (23.31%)
−10000
0
10000
20000
PC1 (60.01%)
Known batch effects controlled
0.5
0.0
−0.5
PC2 (18.92%)
−1.0
−0.5
0.0
0.5
1.0
PC1 (36.25%)
Batch effects controlled + outliers removed
0.4
0.0
−0.4
PC2 (17.04%)
−1.0
−0.5
0.0
0.5
1.0
PC1 (37.18%)
Mean−variance relation in residuals
sd
2
1
0
0
4000
8000
12000
rank(mean)
count
20
7
3
1

E−ENAD−34

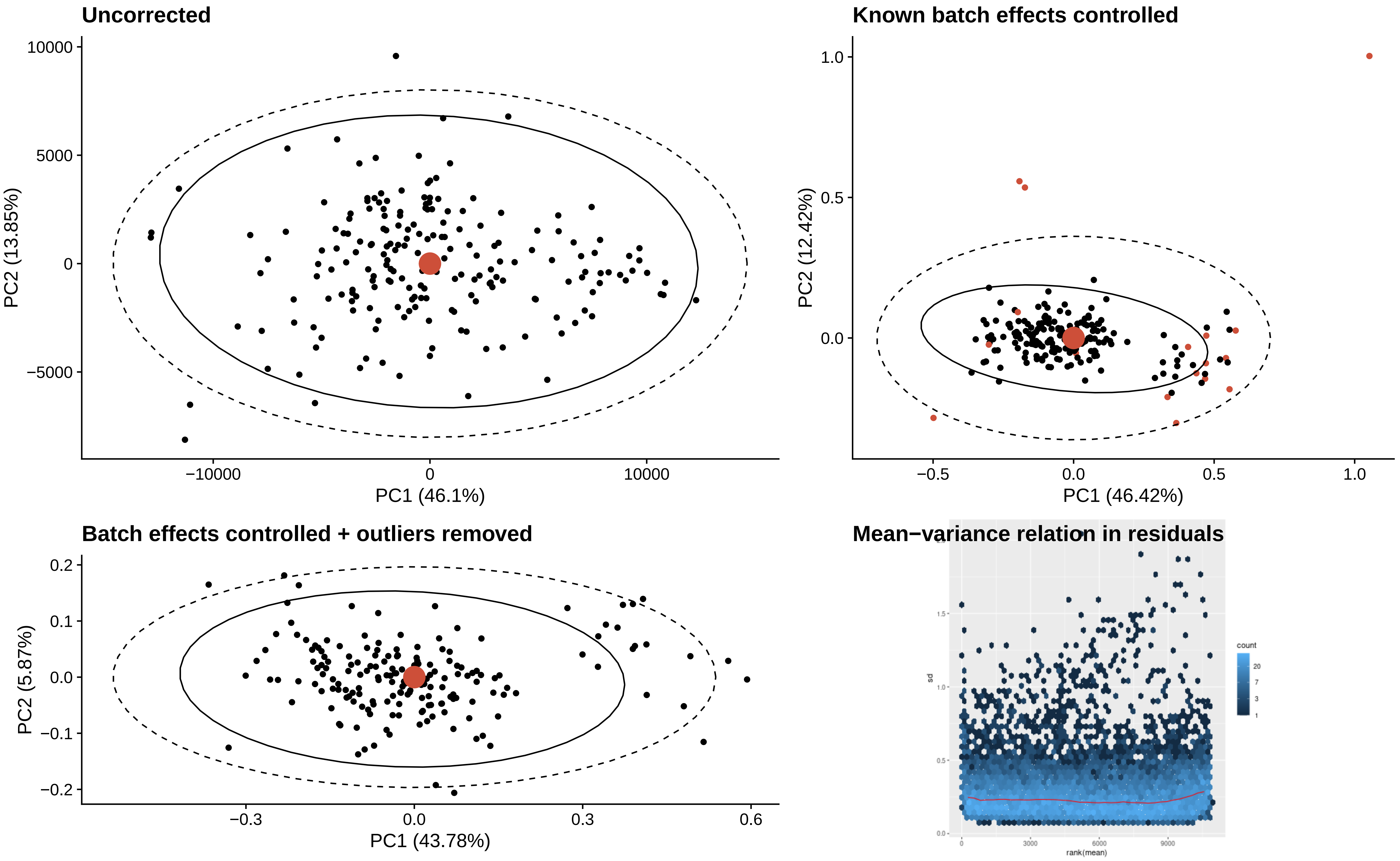
Uncorrected
10000
5000
0
−5000
PC2 (13.85%)
−10000
0
10000
PC1 (46.1%)
Known batch effects controlled
1.0
0.5
0.0
PC2 (12.42%)
−0.5
0.0
0.5
1.0
PC1 (46.42%)
Batch effects controlled + outliers removed
0.2
0.1
0.0
−0.1
−0.2
PC2 (5.87%)
−0.3
0.0
0.3
0.6
PC1 (43.78%)
Mean−variance relation in residuals
sd
rank(mean)
count

E−GEOD−57945

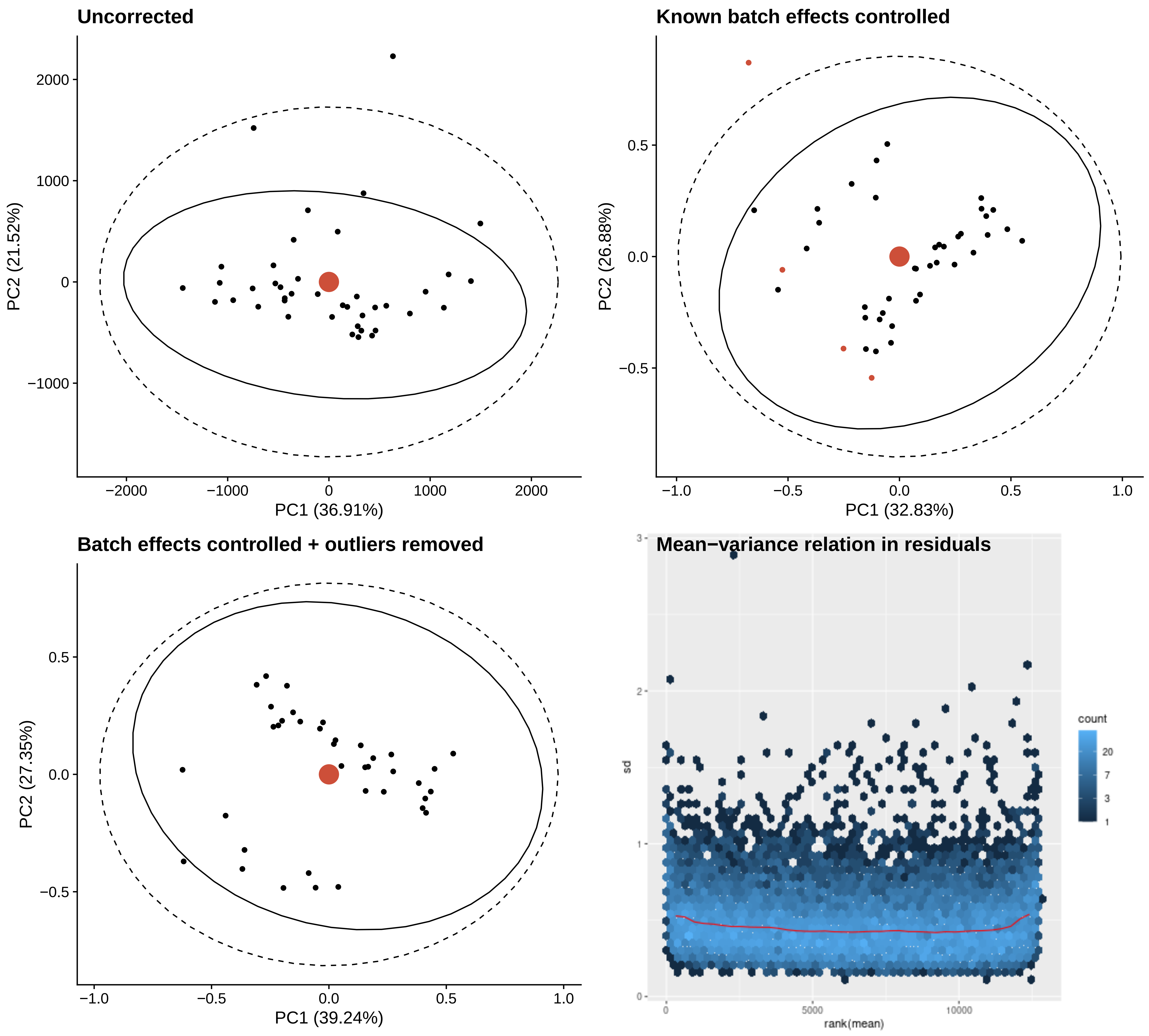

Uncorrected
2000
1000
0
−1000
PC2 (21.52%)
−2000
−1000
0
1000
2000
PC1 (36.91%)
Known batch effects controlled
0.5
0.0
−0.5
PC2 (26.88%)
−1.0
−0.5
0.0
0.5
1.0
PC1 (32.83%)
Batch effects controlled + outliers removed
0.5
0.0
−0.5
PC2 (27.35%)
−1.0
−0.5
0.0
0.5
1.0
PC1 (39.24%)
Mean−variance relation in residuals
3
2
1
0
sd
0
5000
10000
rank(mean)
count
20
7
3
1

E−GEUV−1

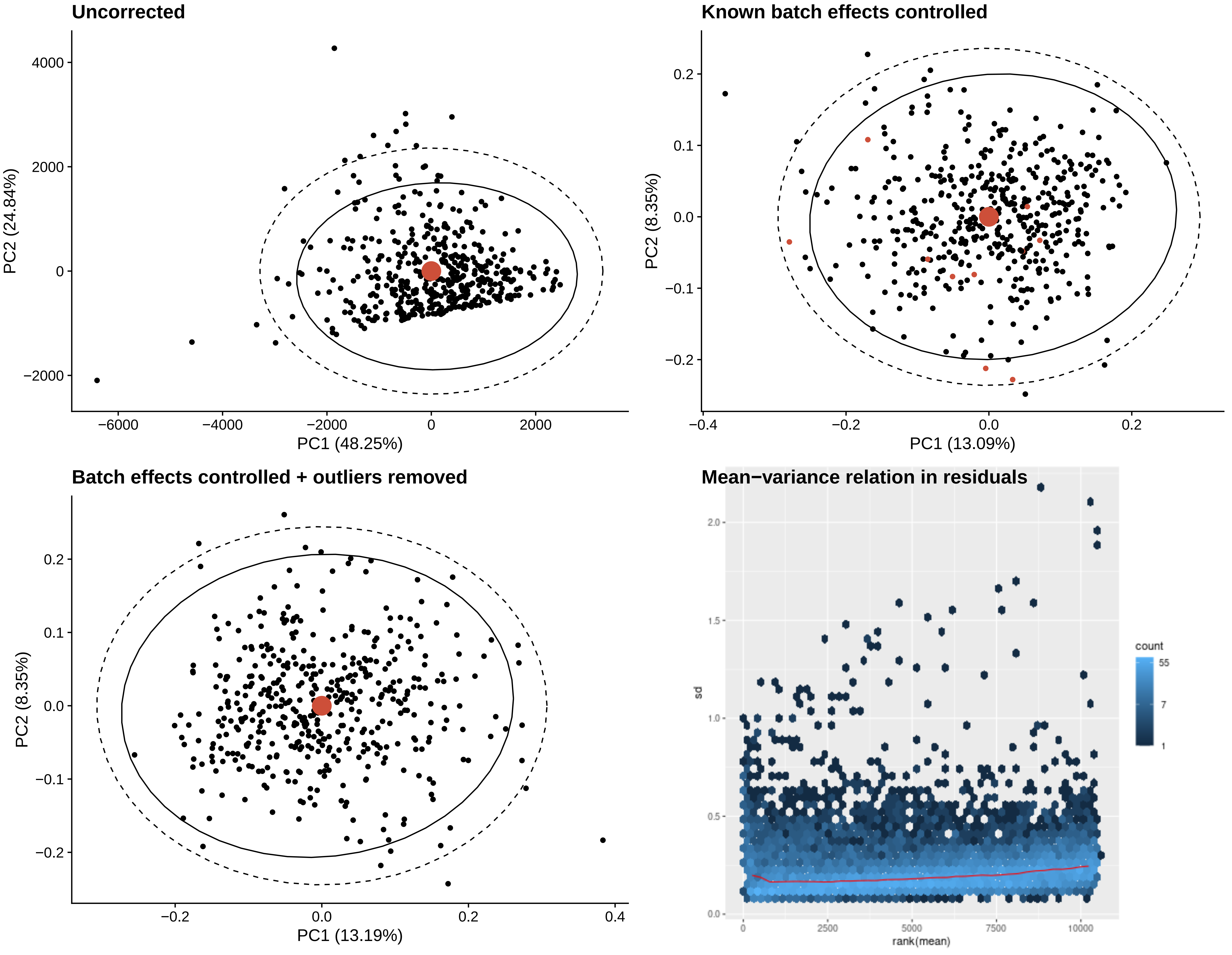

Uncorrected
4000
2000
0
−2000
PC2 (24.84%)
−6000
−4000
−2000
0
2000
PC1 (48.25%)
Known batch effects controlled
0.2
0.1
0.0
−0.1
−0.2
PC2 (8.35%)
−0.4
−0.2
0.0
0.2
PC1 (13.09%)
Batch effects controlled + outliers removed
0.2
0.1
0.0
−0.1
−0.2
PC2 (8.35%)
−0.2
0.0
0.2
0.4
PC1 (13.19%)
Mean−variance relation in residuals
sd
rank(mean)
count
55
7
1

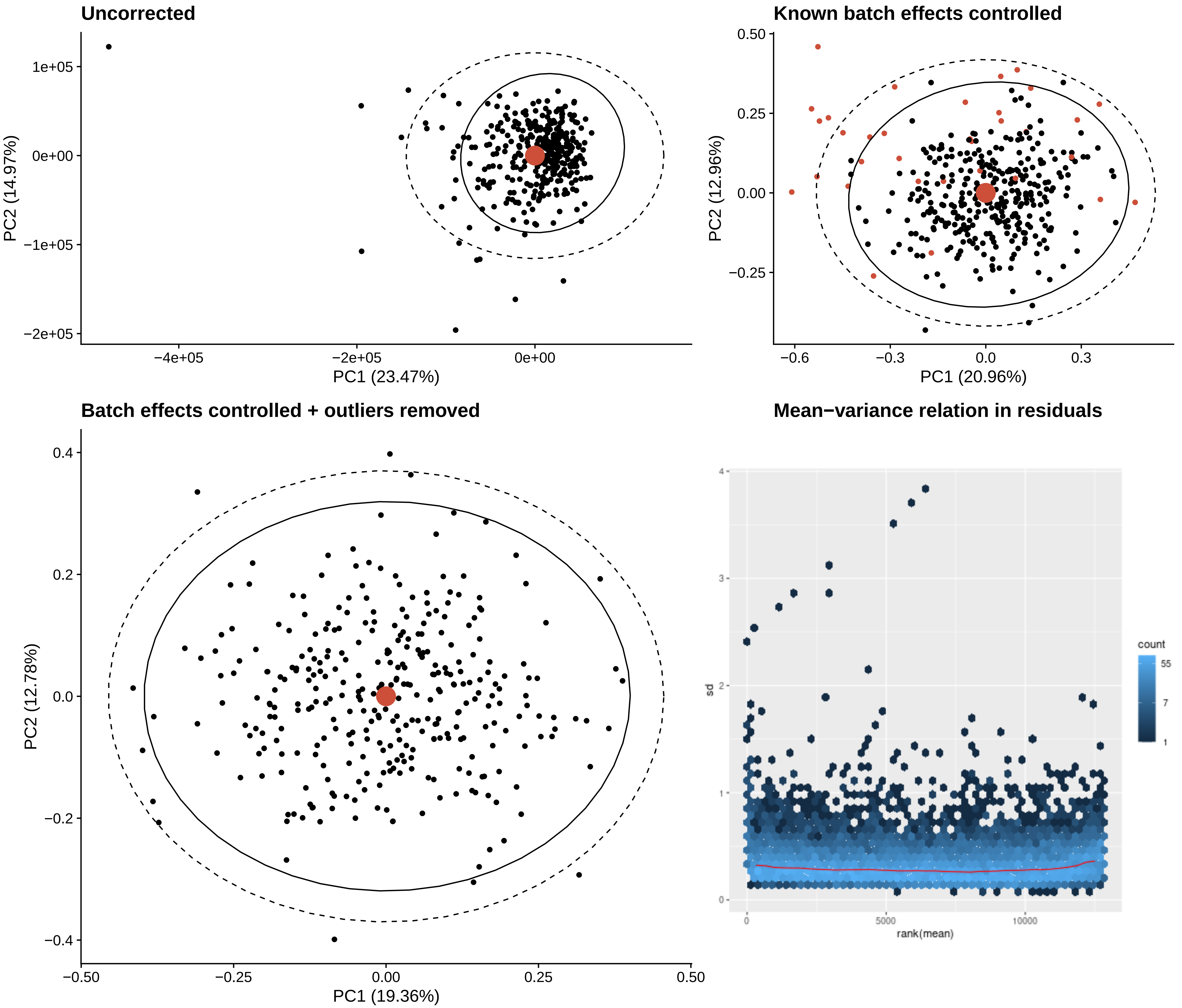
Uncorrected
1e+05
0e+00
−1e+05
−2e+05
PC2 (14.97%)
−4e+05
−2e+05
0e+00
PC1 (23.47%)
Known batch effects controlled
0.50
0.25
0.00
−0.25
PC2 (12.96%)
−0.6
−0.3
0.0
0.3
PC1 (20.96%)
Batch effects controlled + outliers removed
0.4
0.2
0.0
−0.2
−0.4
PC2 (12.78%)
−0.50
−0.25
0.00
0.25
0.50
PC1 (19.36%)
Mean−variance relation in residuals
4
3
2
1
0
sd
0
5000
10000
rank(mean)
count
55
7
1

SRP032775

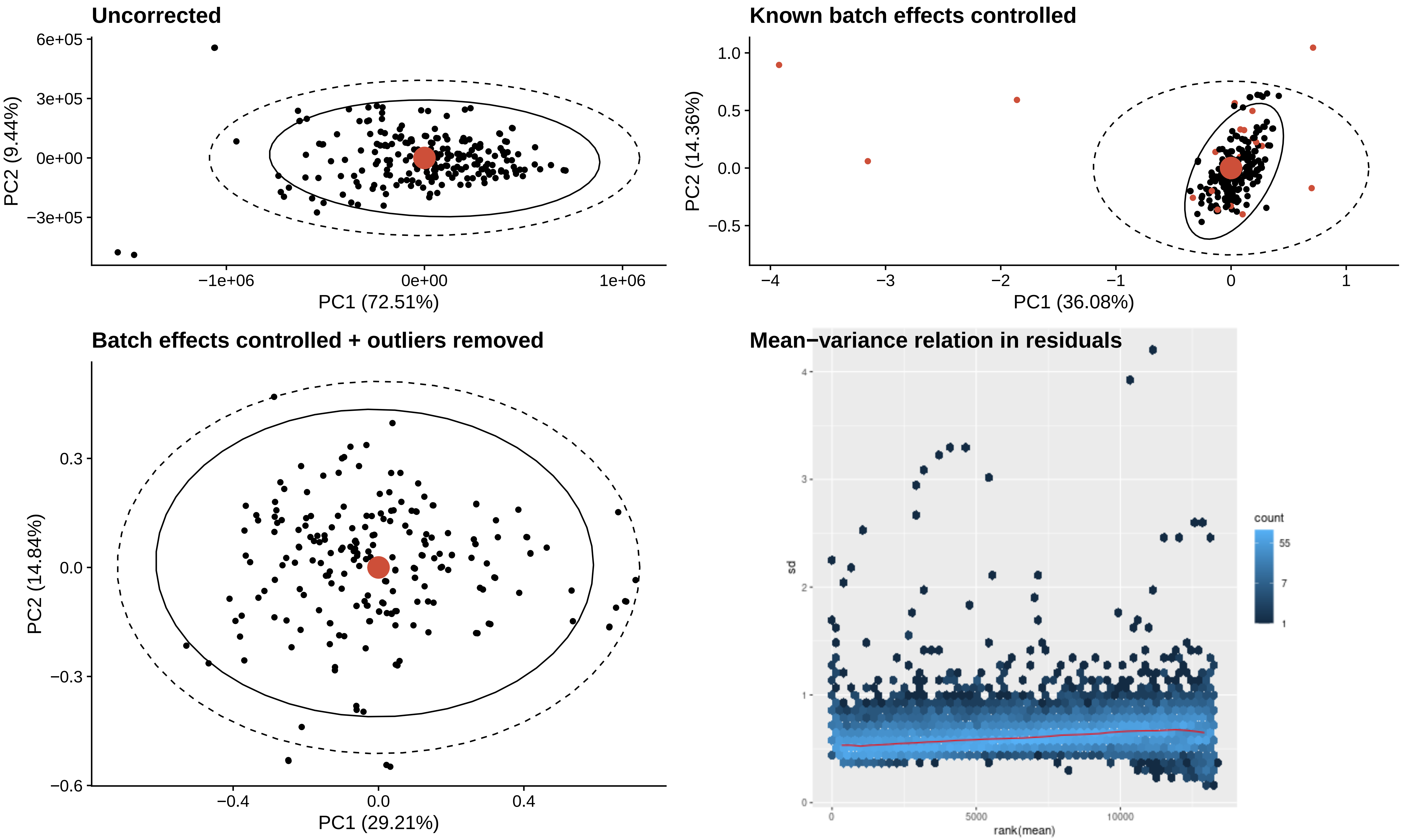
Uncorrected
6e+05
3e+05
0e+00
−3e+05
PC2 (9.44%)
−1e+06
0e+00
1e+06
PC1 (72.51%)
Known batch effects controlled
1.0
0.5
0.0
−0.5
PC2 (14.36%)
−4
−3
−2
−1
0
1
PC1 (36.08%)
Batch effects controlled + outliers removed
0.3
0.0
−0.3
−0.6
PC2 (14.84%)
−0.4
0.0
0.4
PC1 (29.21%)
Mean−variance relation in residuals
sd
rank(mean)
count
55
7
1
0
5000
10000

SRP035988

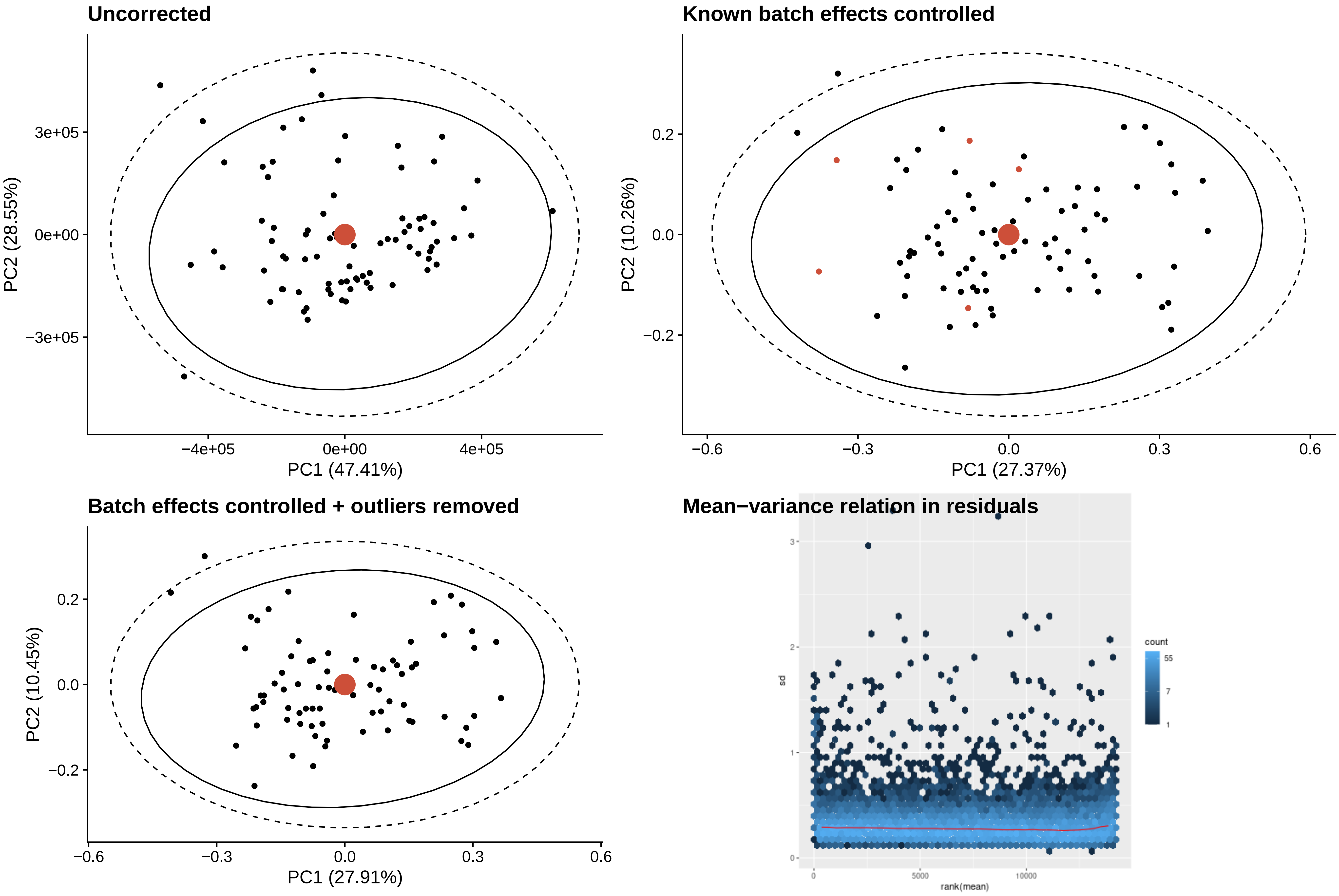

Uncorrected
3e+05
0e+00
−3e+05
PC2 (28.55%)
−4e+05
0e+00
4e+05
PC1 (47.41%)
Known batch effects controlled
0.2
0.0
−0.2
PC2 (10.26%)
−0.6
−0.3
0.0
0.3
0.6
PC1 (27.37%)
Batch effects controlled + outliers removed
0.2
0.0
−0.2
PC2 (10.45%)
−0.6
−0.3
0.0
0.3
0.6
PC1 (27.91%)
Mean−variance relation in residuals
sd
rank(mean)
count
55
7
1

SRP051848

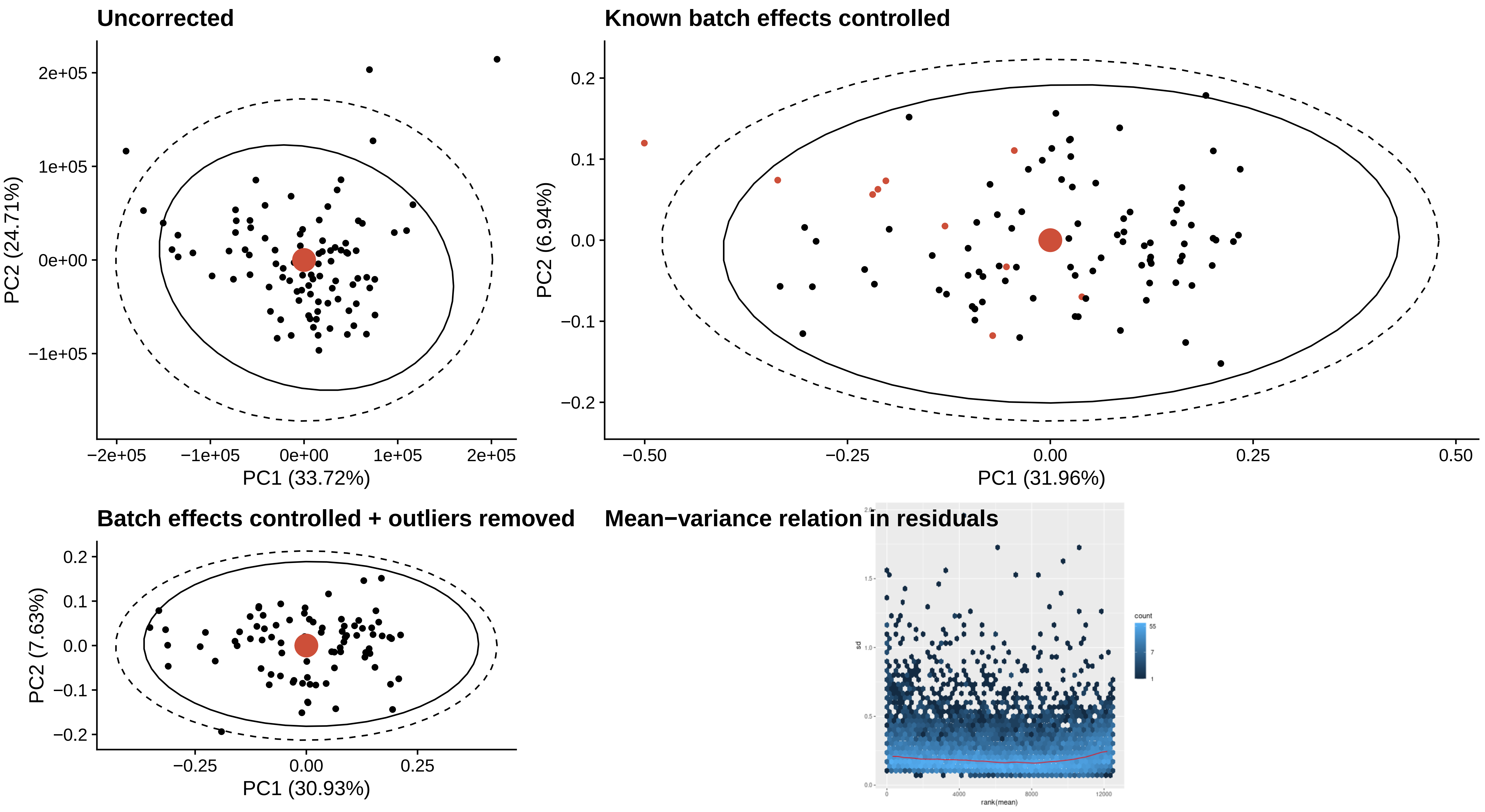
Uncorrected
2e+05
1e+05
0e+00
−1e+05
PC2 (24.71%)
−2e+05
−1e+05
0e+00
1e+05
2e+05
PC1 (33.72%)
Known batch effects controlled
0.2
0.1
0.0
−0.1
−0.2
PC2 (6.94%)
−0.50
−0.25
0.00
0.25
0.50
PC1 (31.96%)
Batch effects controlled + outliers removed
0.2
0.1
0.0
−0.1
−0.2
PC2 (7.63%)
−0.25
0.00
0.25
PC1 (30.93%)
Mean−variance relation in residuals

SRP057500

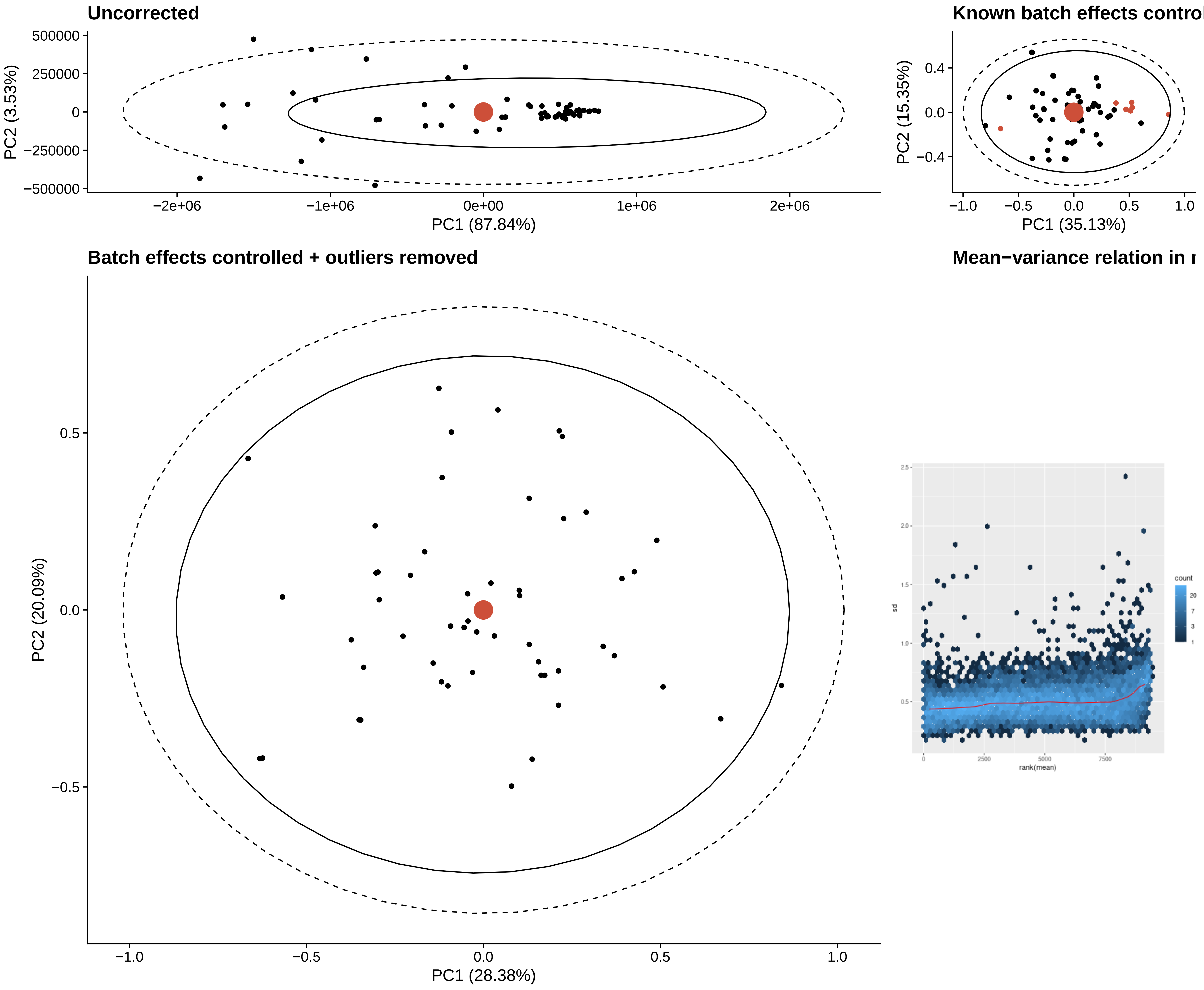

Uncorrected
500000
250000
0
−250000
−500000
PC2 (3.53%)
−2e+06
−1e+06
0e+00
1e+06
2e+06
PC1 (87.84%)
Known batch effects controll
0.4
0.0
−0.4
PC2 (15.35%)
−1.0
−0.5
0.0
0.5
1.0
PC1 (35.13%)
Batch effects controlled + outliers removed
0.5
0.0
−0.5
PC2 (20.09%)
−1.0
−0.5
0.0
0.5
1.0
PC1 (28.38%)
Mean−variance relation in r
sd
rank(mean)
count

SRP059039

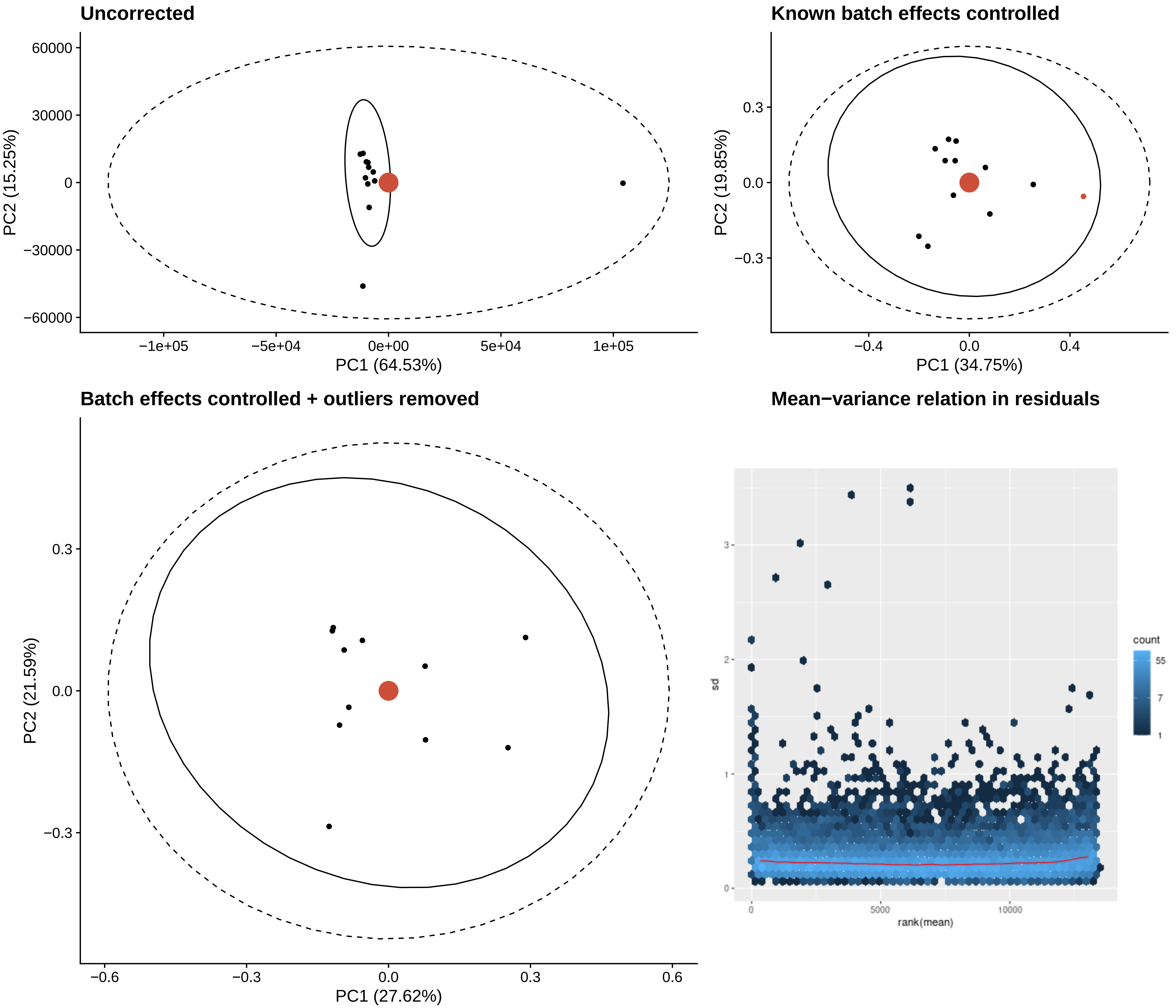
Uncorrected
60000
30000
0
−30000
−60000
PC2 (15.25%)
−1e+05
−5e+04
0e+00
5e+04
1e+05
PC1 (64.53%)
Known batch effects controlled
0.3
0.0
−0.3
PC2 (19.85%)
−0.4
0.0
0.4
PC1 (34.75%)
Batch effects controlled + outliers removed
0.3
0.0
−0.3
PC2 (21.59%)
−0.6
−0.3
0.0
0.3
0.6
PC1 (27.62%)
Mean−variance relation in residuals
sd
3
2
1
0
0
5000
10000
rank(mean)
count
55
7
1

SRP069212

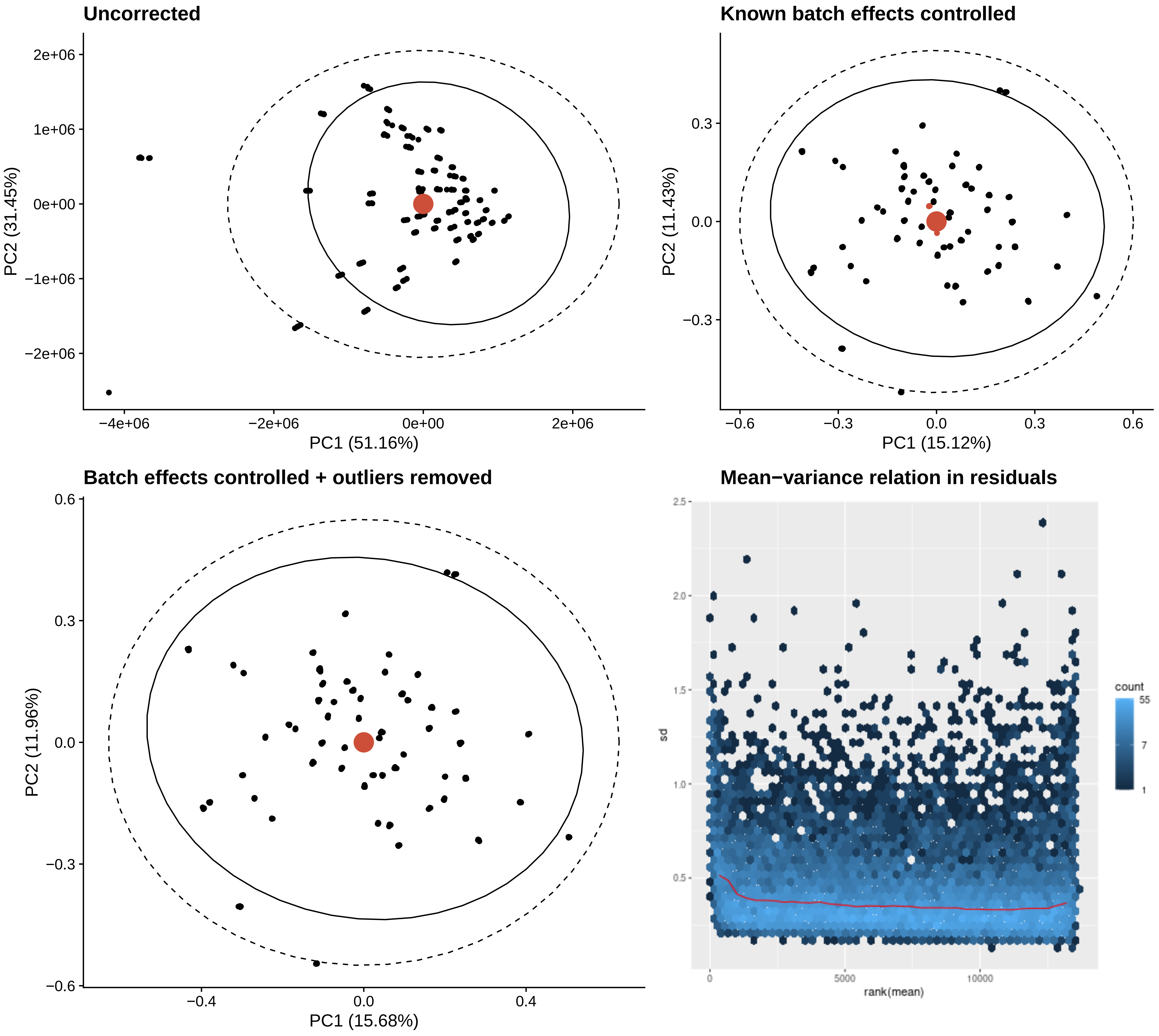

Uncorrected
2e+06
1e+06
0e+00
−1e+06
−2e+06
PC2 (31.45%)
−4e+06
−2e+06
0e+00
2e+06
PC1 (51.16%)
Known batch effects controlled
0.3
0.0
−0.3
PC2 (11.43%)
−0.6
−0.3
0.0
0.3
0.6
PC1 (15.12%)
Batch effects controlled + outliers removed
0.6
0.3
0.0
−0.3
−0.6
PC2 (11.96%)
−0.4
0.0
0.4
PC1 (15.68%)
Mean−variance relation in residuals
2.5
2.0
1.5
1.0
0.5
sd
0
5000
10000
rank(mean)
count
55
7
1

SRP095272

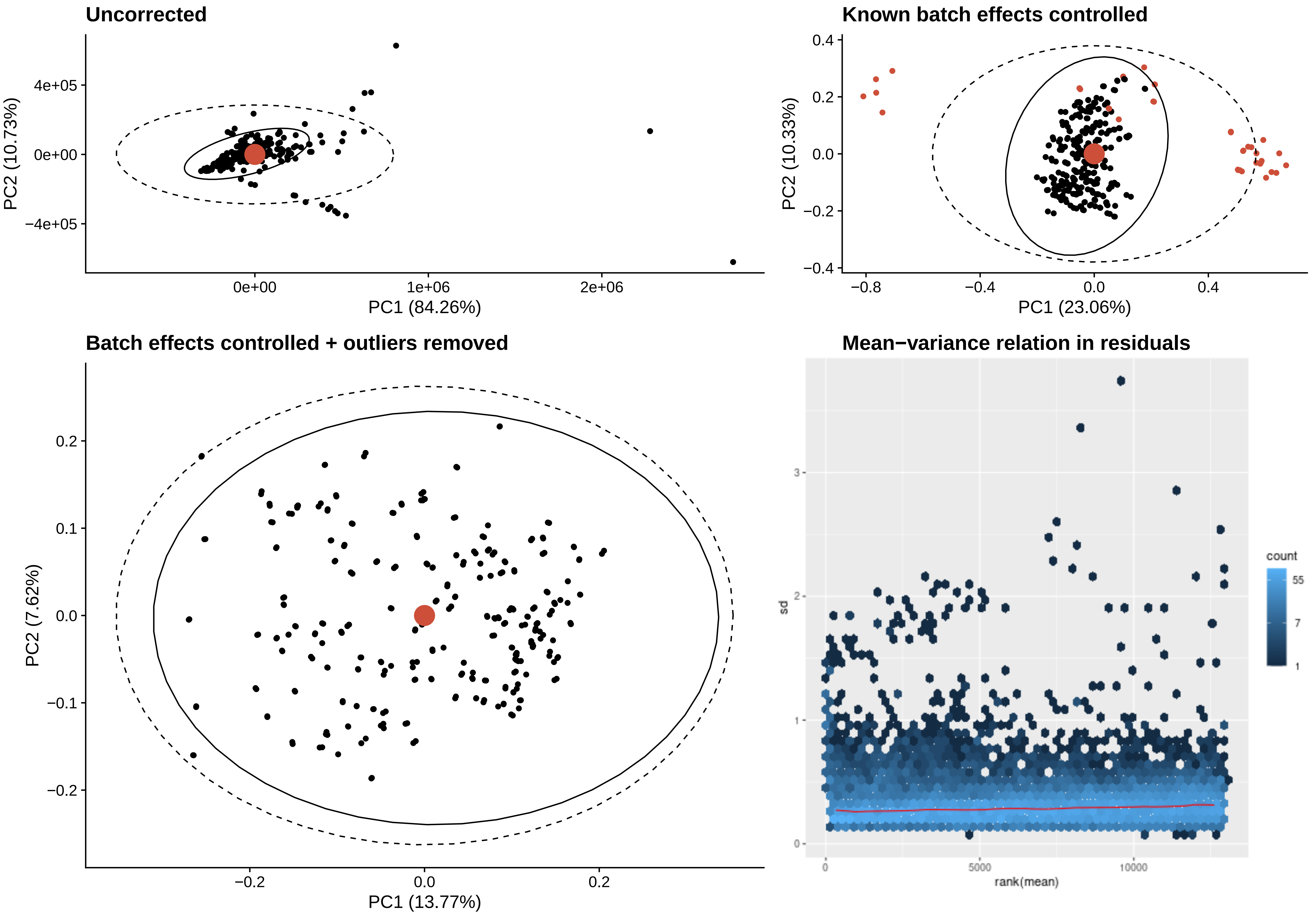

Uncorrected
4e+05
0e+00
−4e+05
PC2 (10.73%)
0e+00
1e+06
2e+06
PC1 (84.26%)
Known batch effects controlled
0.4
0.2
0.0
−0.2
−0.4
PC2 (10.33%)
−0.8
−0.4
0.0
0.4
PC1 (23.06%)
Batch effects controlled + outliers removed
0.2
0.1
0.0
−0.1
−0.2
PC2 (7.62%)
−0.2
0.0
0.2
PC1 (13.77%)
Mean−variance relation in residuals
sd
3
2
1
0
0
5000
10000
rank(mean)
count
55
7
1

SRP098758

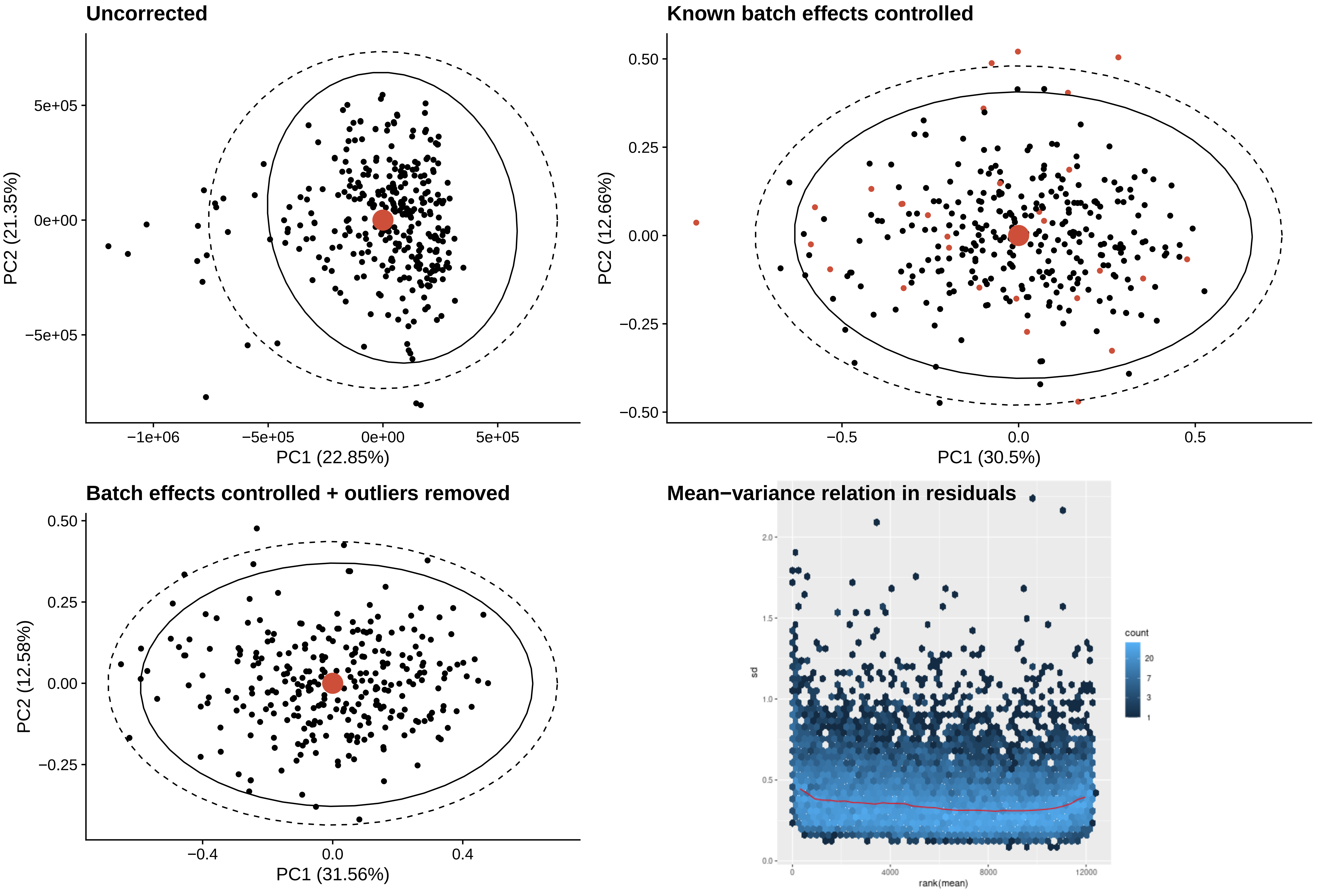
Uncorrected
PC2 (21.35%)
PC1 (22.85%)
5e+05
0e+00
−5e+05
−1e+06
−5e+05
0e+00
5e+05
Known batch effects controlled
PC2 (12.66%)
PC1 (30.5%)
0.50
0.25
0.00
−0.25
−0.50
−0.5
0.0
0.5
Batch effects controlled + outliers removed
PC2 (12.58%)
PC1 (31.56%)
0.50
0.25
0.00
−0.25
−0.4
0.0
0.4
Mean−variance relation in residuals
sd
rank(mean)
count
20
7
3
1
2.0
1.5
1.0
0.5
0.0
0
4000
8000
12000

SRP101294

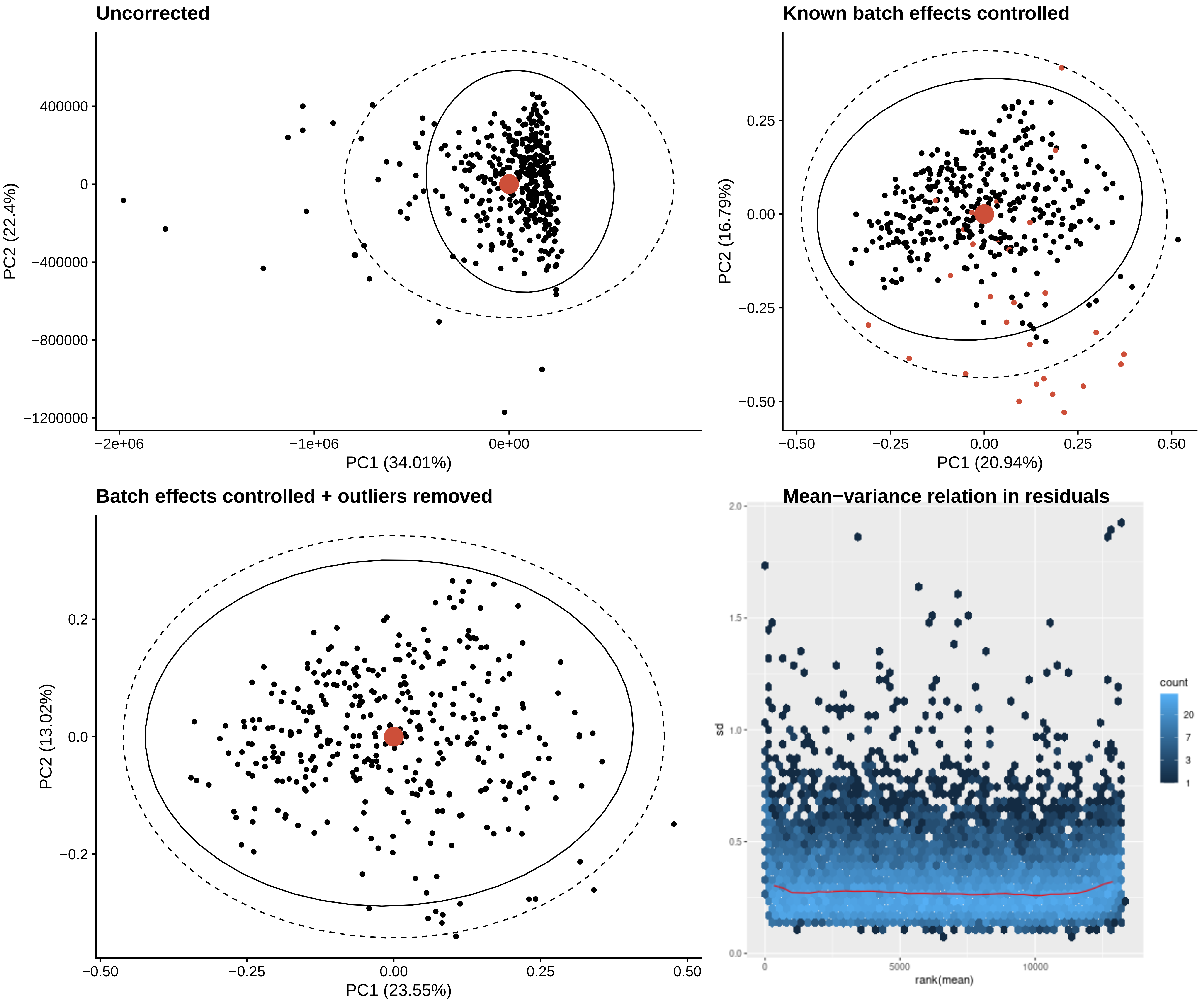

Uncorrected
400000
0
−400000
−800000
−1200000
PC2 (22.4%)
−2e+06
−1e+06
0e+00
PC1 (34.01%)
Known batch effects controlled
0.25
0.00
−0.25
−0.50
PC2 (16.79%)
−0.50
−0.25
0.00
0.25
0.50
PC1 (20.94%)
Batch effects controlled + outliers removed
0.2
0.0
−0.2
PC2 (13.02%)
−0.50
−0.25
0.00
0.25
0.50
PC1 (23.55%)
Mean−variance relation in residuals
2.0
1.5
1.0
0.5
0.0
sd
0
5000
10000
rank(mean)
count
20
7
3
1

SRP102999

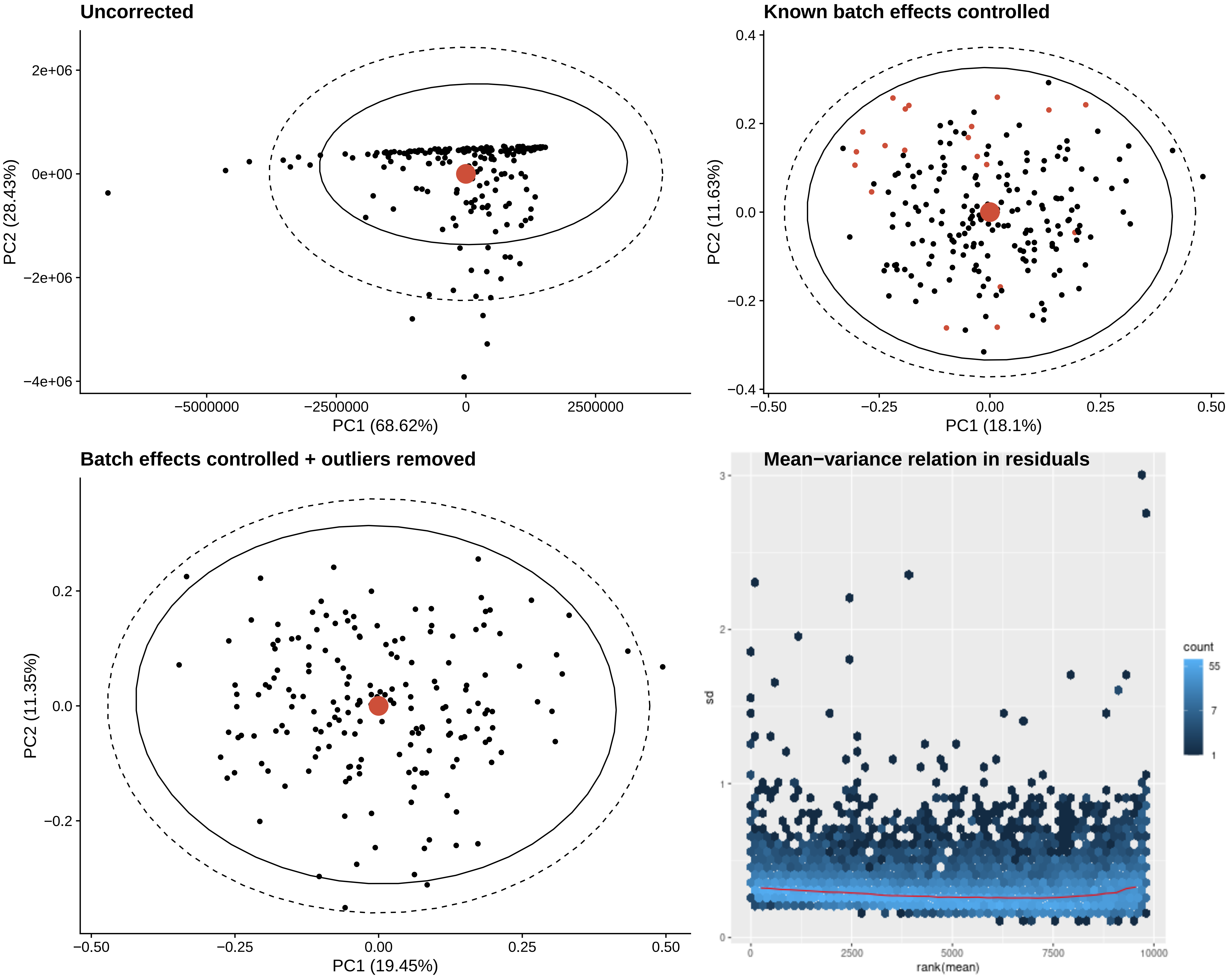

Uncorrected
2e+06
0e+00
−2e+06
−4e+06
PC2 (28.43%)
−5000000
−2500000
0
2500000
PC1 (68.62%)
Known batch effects controlled
0.4
0.2
0.0
−0.2
−0.4
PC2 (11.63%)
−0.50
−0.25
0.00
0.25
0.50
PC1 (18.1%)
Batch effects controlled + outliers removed
0.2
0.0
−0.2
PC2 (11.35%)
−0.50
−0.25
0.00
0.25
0.50
PC1 (19.45%)
Mean−variance relation in residuals
3
2
1
0
sd
0
2500
5000
7500
10000
rank(mean)
count
55
7
1

SRP145493

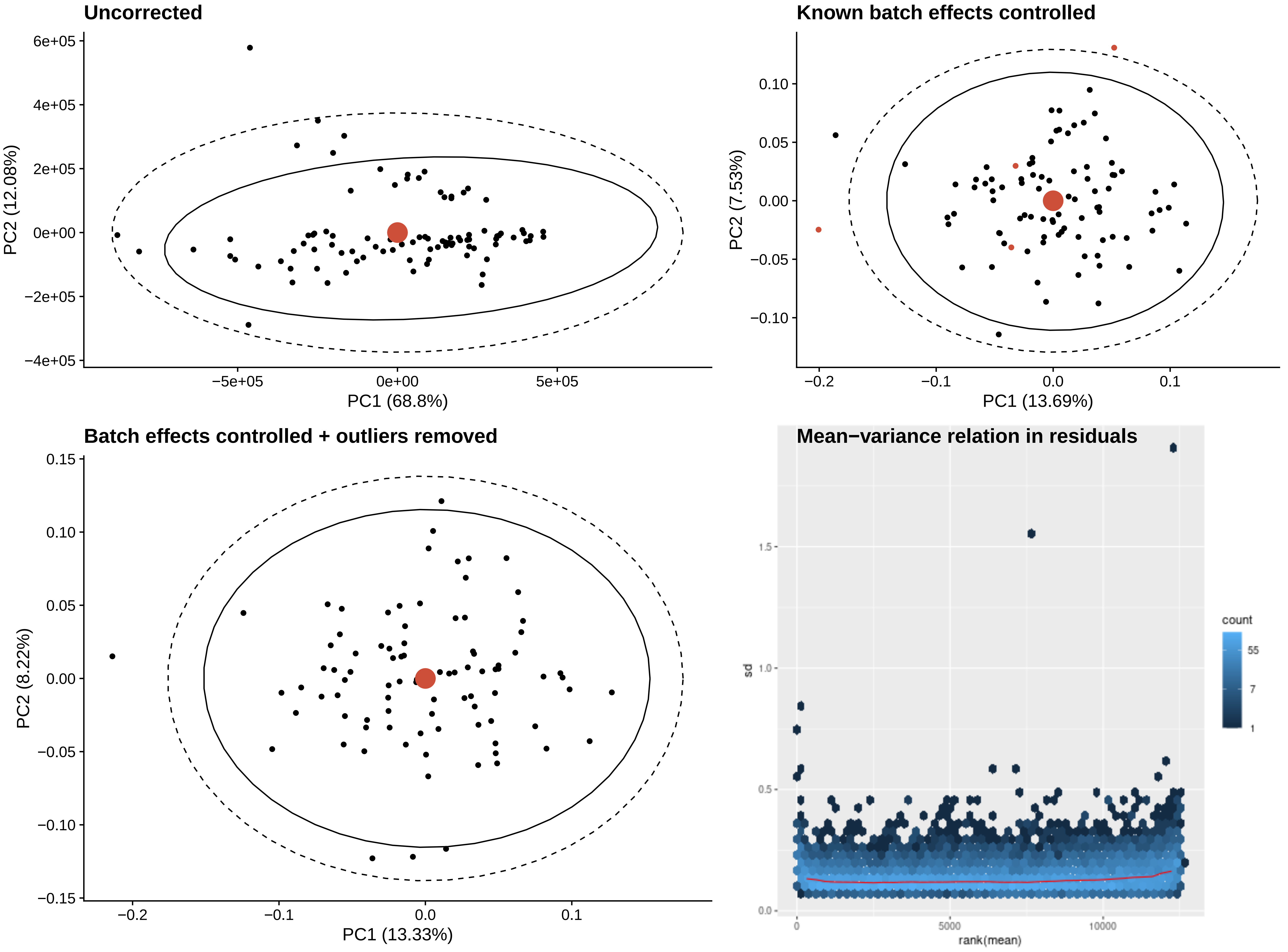

Uncorrected
PC1 (68.8%)
PC2 (12.08%)
Known batch effects controlled
PC1 (13.69%)
PC2 (7.53%)
Batch effects controlled + outliers removed
PC1 (13.33%)
PC2 (8.22%)
Mean−variance relation in residuals
rank(mean)
sd
count

SRP150552

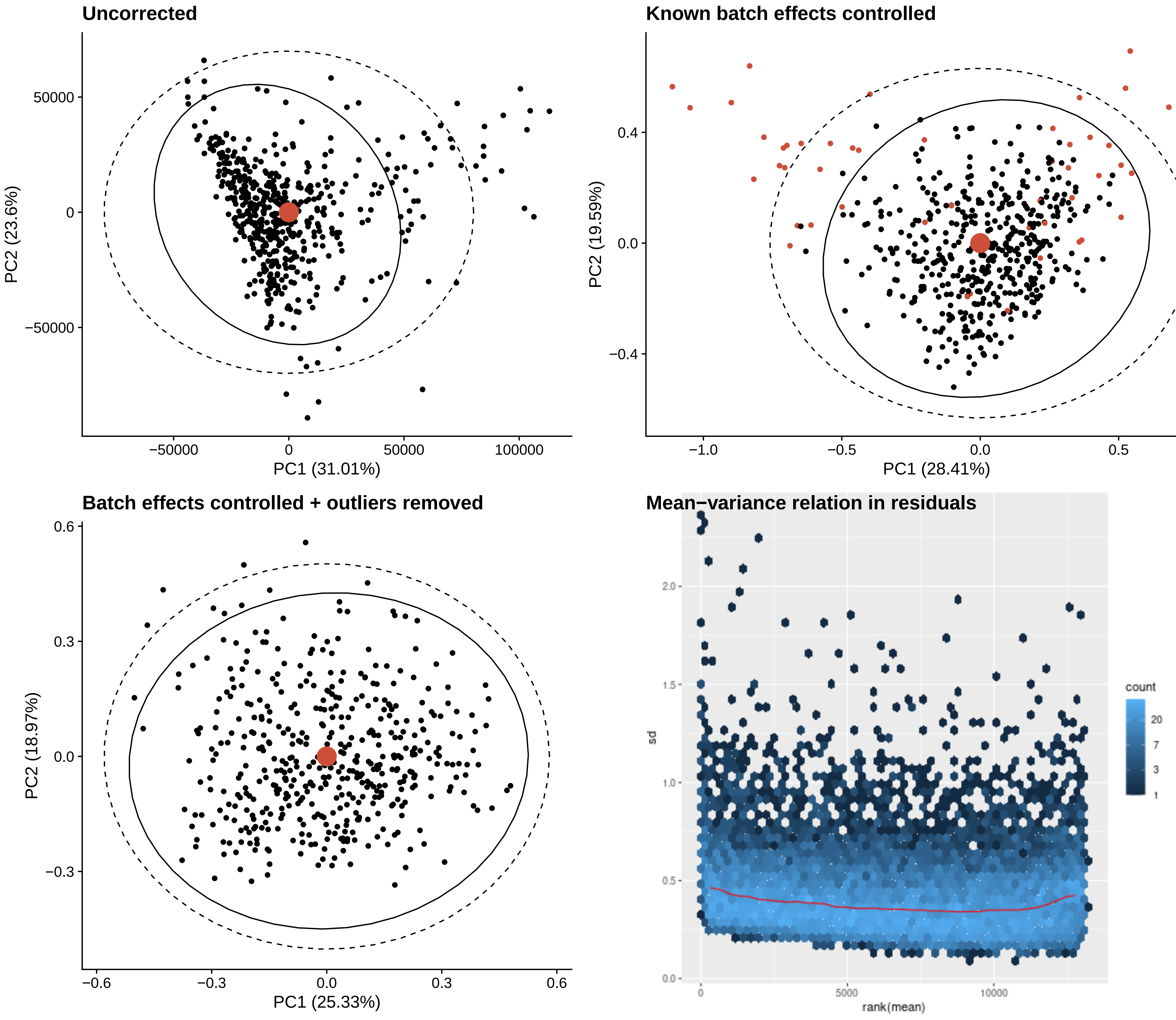
Uncorrected
PC2 (23.6%)
PC1 (31.01%)
50000
0
−50000
−50000
0
50000
100000
Known batch effects controlled
PC2 (19.59%)
PC1 (28.41%)
0.4
0.0
−0.4
−1.0
−0.5
0.0
0.5
Batch effects controlled + outliers removed
PC2 (18.97%)
PC1 (25.33%)
0.6
0.3
0.0
−0.3
−0.6
−0.3
0.0
0.3
0.6
Mean−variance relation in residuals
sd
rank(mean)
2.0
1.5
1.0
0.5
0.0
0
5000
10000
count
20
7
3
1

SRP162654

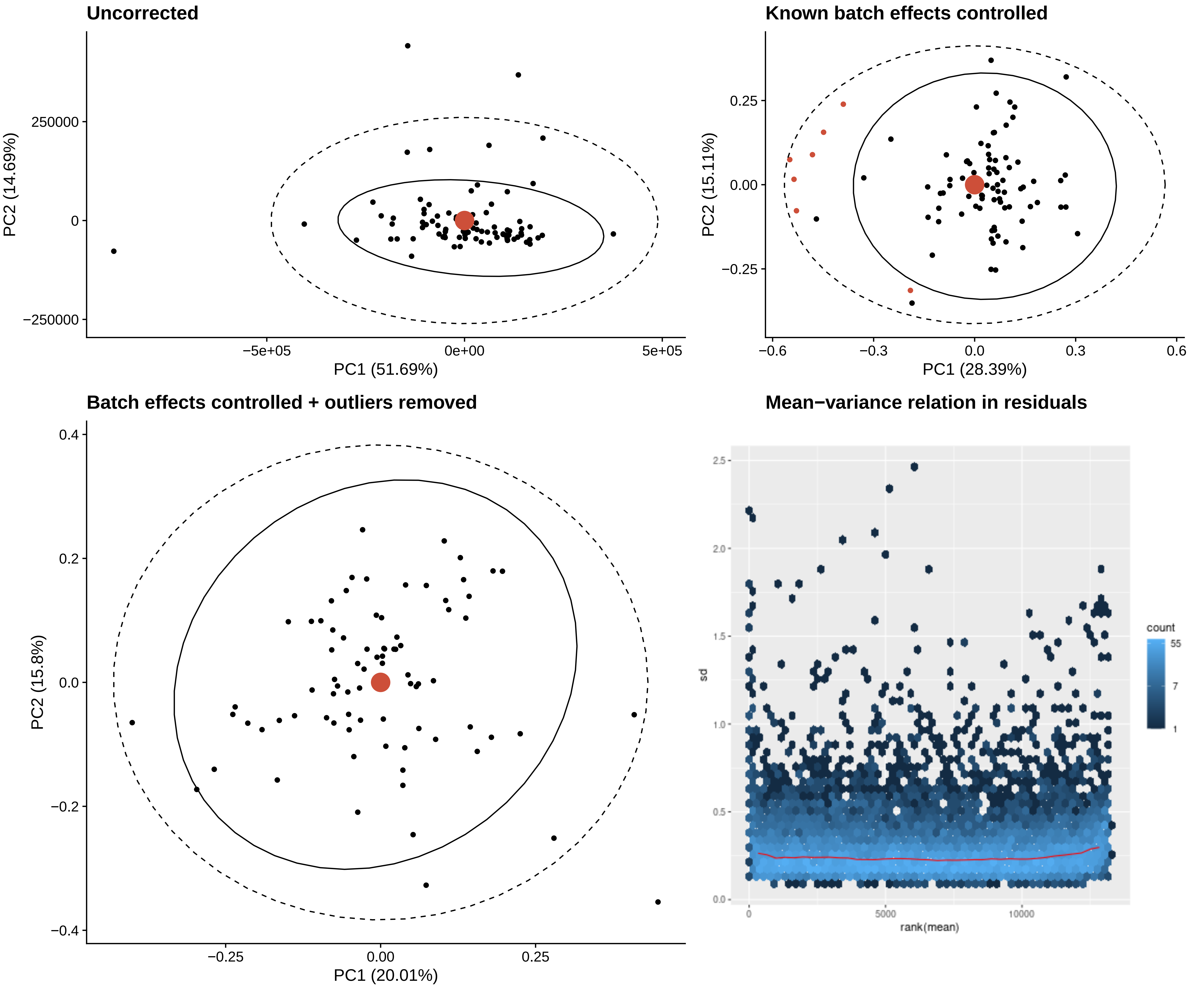

Uncorrected
250000
0
−250000
−5e+05
0e+00
5e+05
PC1 (51.69%)
PC2 (14.69%)
Known batch effects controlled
0.25
0.00
−0.25
−0.6
−0.3
0.0
0.3
0.6
PC1 (28.39%)
PC2 (15.11%)
Batch effects controlled + outliers removed
0.4
0.2
0.0
−0.2
−0.4
−0.25
0.00
0.25
PC1 (20.01%)
PC2 (15.8%)
Mean−variance relation in residuals
2.5
2.0
1.5
1.0
0.5
0.0
0
5000
10000
rank(mean)
sd
count
55
7
1

SRP174638

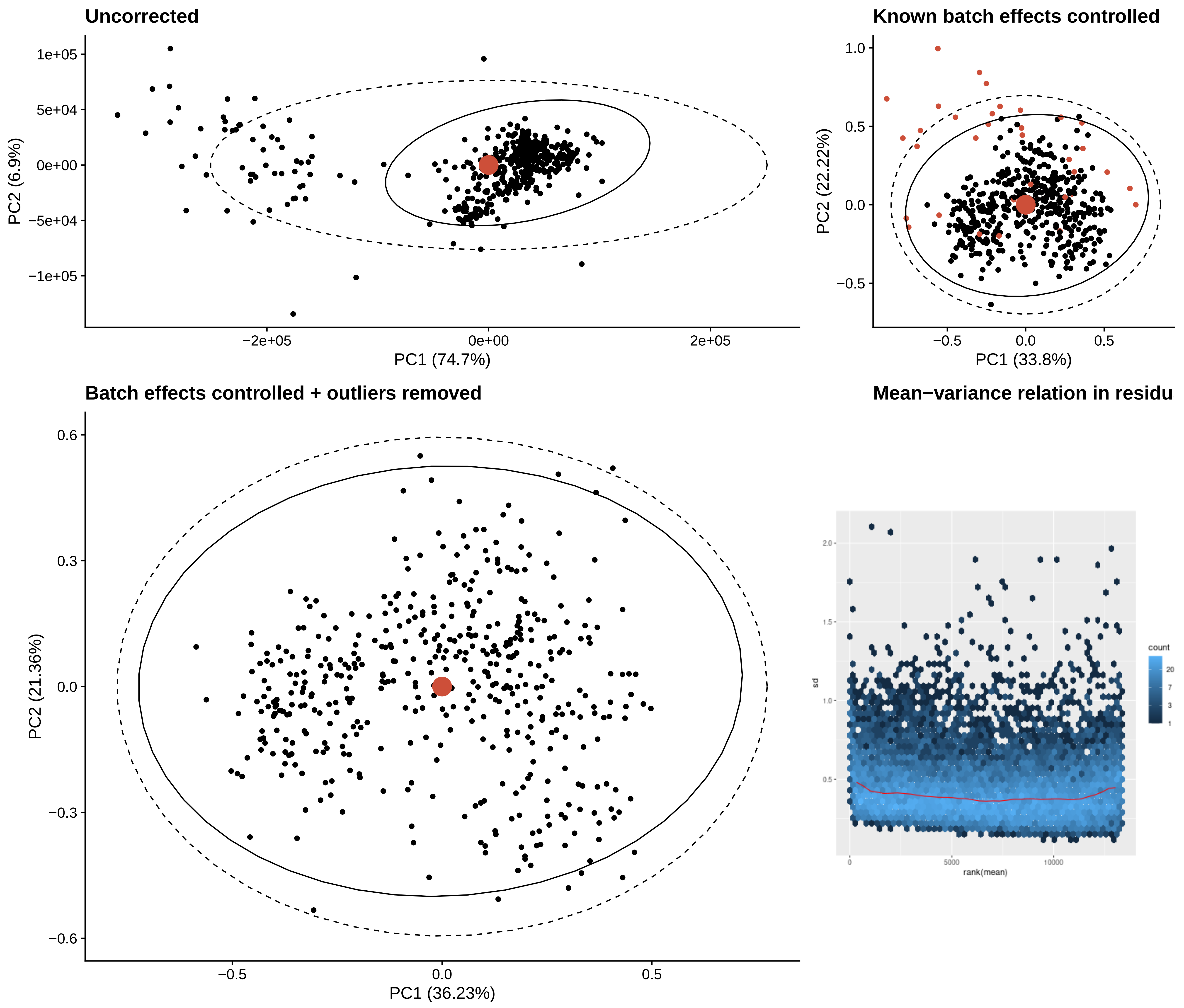
Uncorrected
PC2 (6.9%)
PC1 (74.7%)
1e+05
5e+04
0e+00
−5e+04
−1e+05
−2e+05
0e+00
2e+05
Known batch effects controlled
PC2 (22.22%)
PC1 (33.8%)
1.0
0.5
0.0
−0.5
Batch effects controlled + outliers removed
PC2 (21.36%)
PC1 (36.23%)
0.6
0.3
0.0
−0.3
−0.6
−0.5
0.0
0.5
Mean−variance relation in residu
sd
rank(mean)
count
20
7
3
1
2.0
1.5
1.0
0.5
0
5000
10000

SRP181886

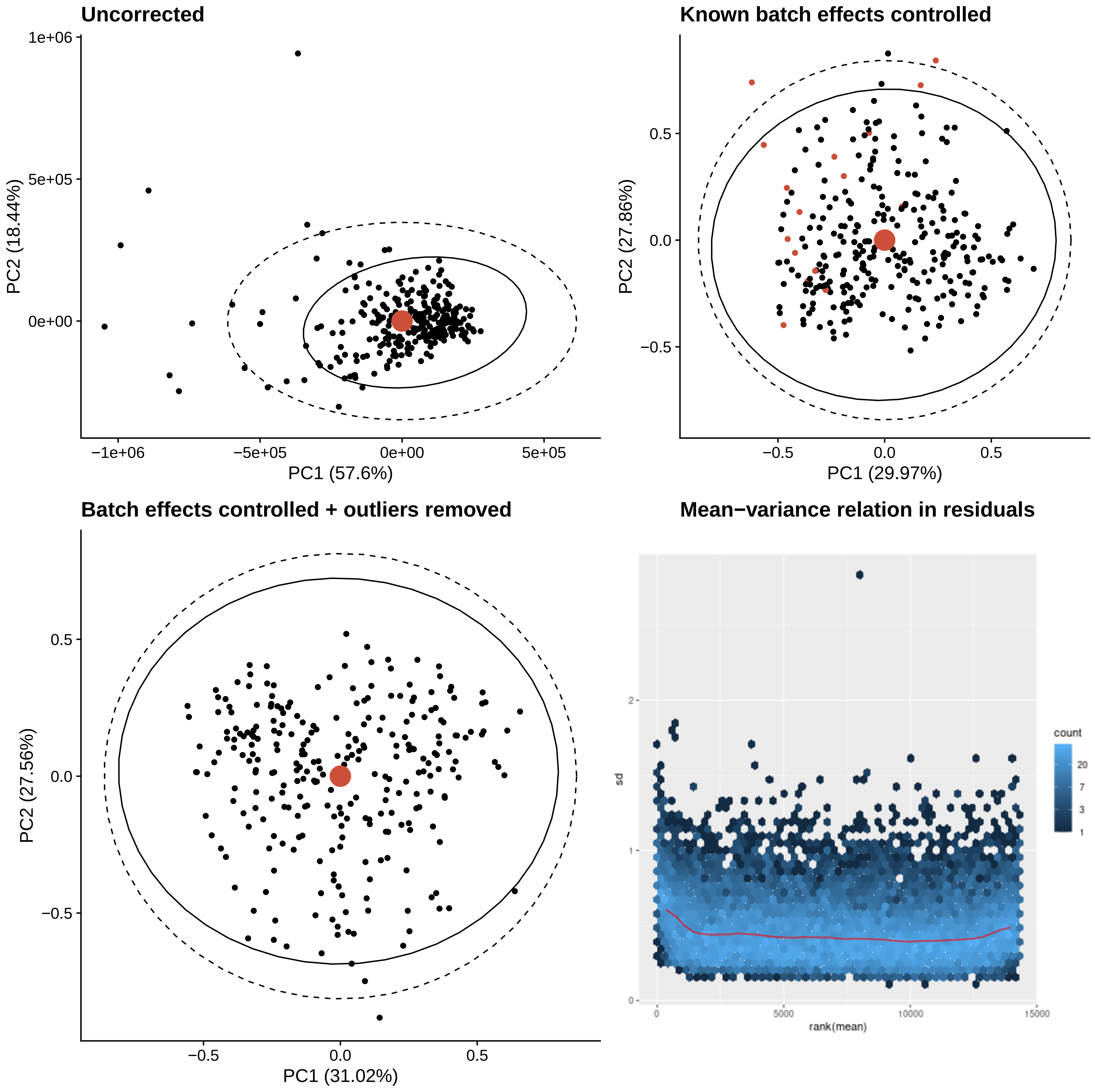
Uncorrected
1e+06
5e+05
0e+00
PC2 (18.44%)
−1e+06
−5e+05
0e+00
5e+05
PC1 (57.6%)
Known batch effects controlled
0.5
0.0
−0.5
PC2 (27.86%)
−0.5
0.0
0.5
PC1 (29.97%)
Batch effects controlled + outliers removed
0.5
0.0
−0.5
PC2 (27.56%)
−0.5
0.0
0.5
PC1 (31.02%)
Mean−variance relation in residuals
sd
2
1
0
0
5000
10000
15000
rank(mean)
count
20
7
3
1

SRP187978

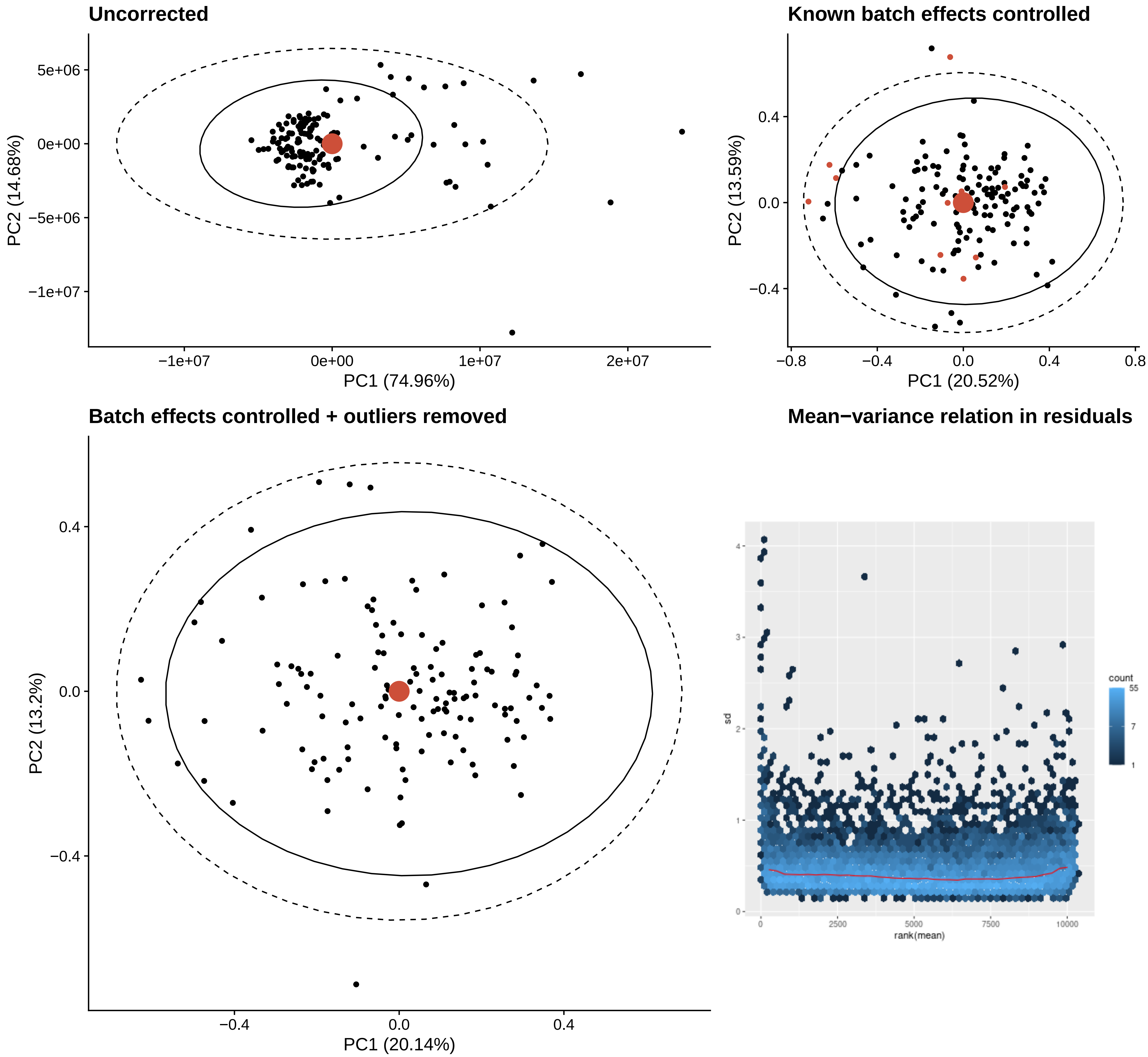
Uncorrected
PC2 (14.68%)
5e+06
0e+00
−5e+06
−1e+07
−1e+07
0e+00
1e+07
2e+07
PC1 (74.96%)
Known batch effects controlled
PC2 (13.59%)
0.4
0.0
−0.4
−0.8
−0.4
0.0
0.4
0.8
PC1 (20.52%)
Batch effects controlled + outliers removed
PC2 (13.2%)
0.4
0.0
−0.4
−0.4
0.0
0.4
PC1 (20.14%)
Mean−variance relation in residuals
sd
4
3
2
1
0
0
2500
5000
7500
10000
rank(mean)
count
55
7
1

SRP192714

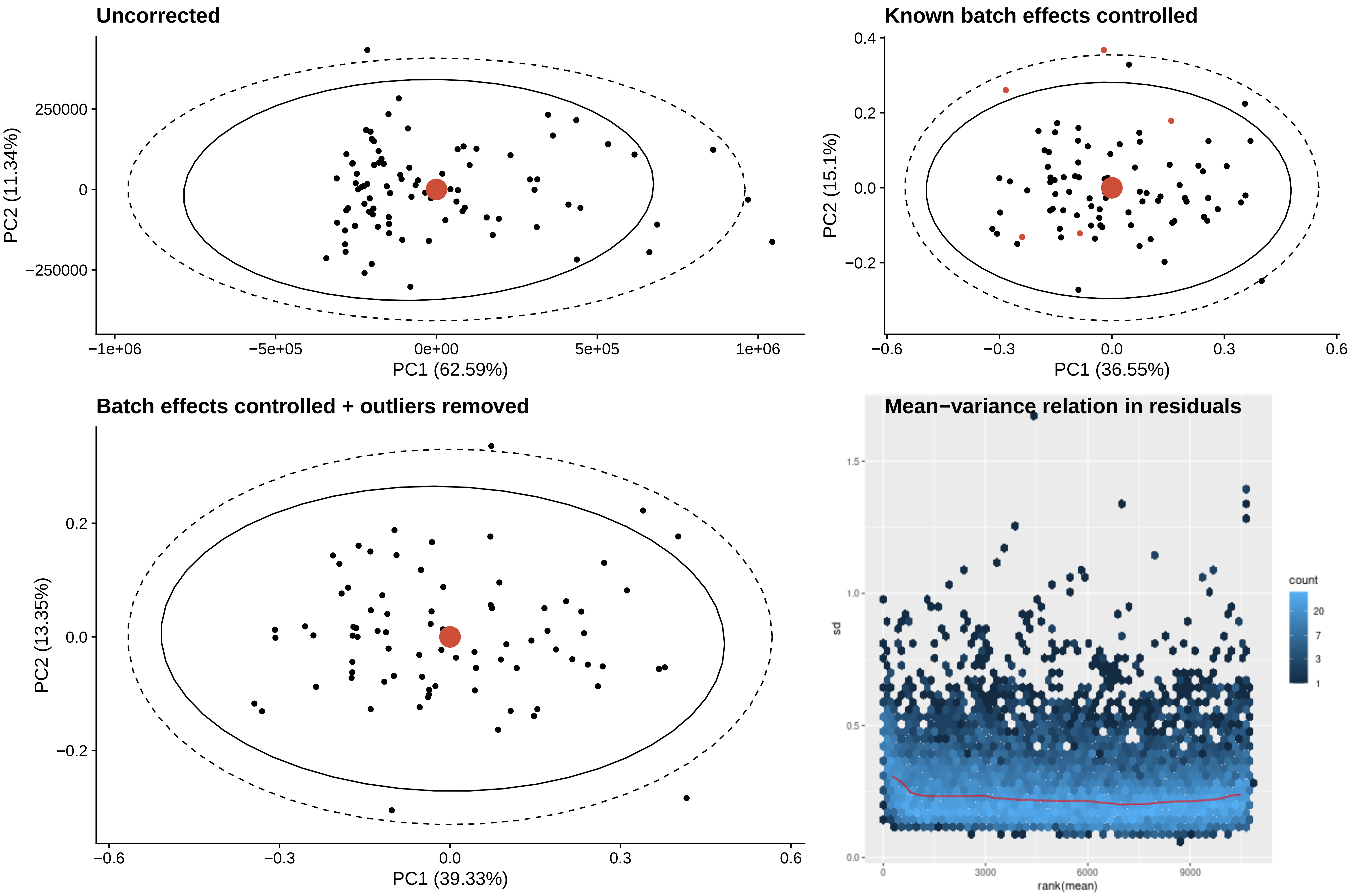

Uncorrected
PC2 (11.34%)
250000
0
−250000
−1e+06
−5e+05
0e+00
5e+05
1e+06
PC1 (62.59%)
Known batch effects controlled
PC2 (15.1%)
0.4
0.2
0.0
−0.2
−0.6
−0.3
0.0
0.3
0.6
PC1 (36.55%)
Batch effects controlled + outliers removed
PC2 (13.35%)
0.2
0.0
−0.2
−0.6
−0.3
0.0
0.3
0.6
PC1 (39.33%)
Mean−variance relation in residuals
sd
1.5
1.0
0.5
0.0
0
3000
6000
9000
rank(mean)
count
20
7
3
1
