## Supporting Information Folder for "Characterizing the landscape of gene expression variance in humans": SI_figures_table1.docx (from another version of this preprint: version 2, 2023-01-09)

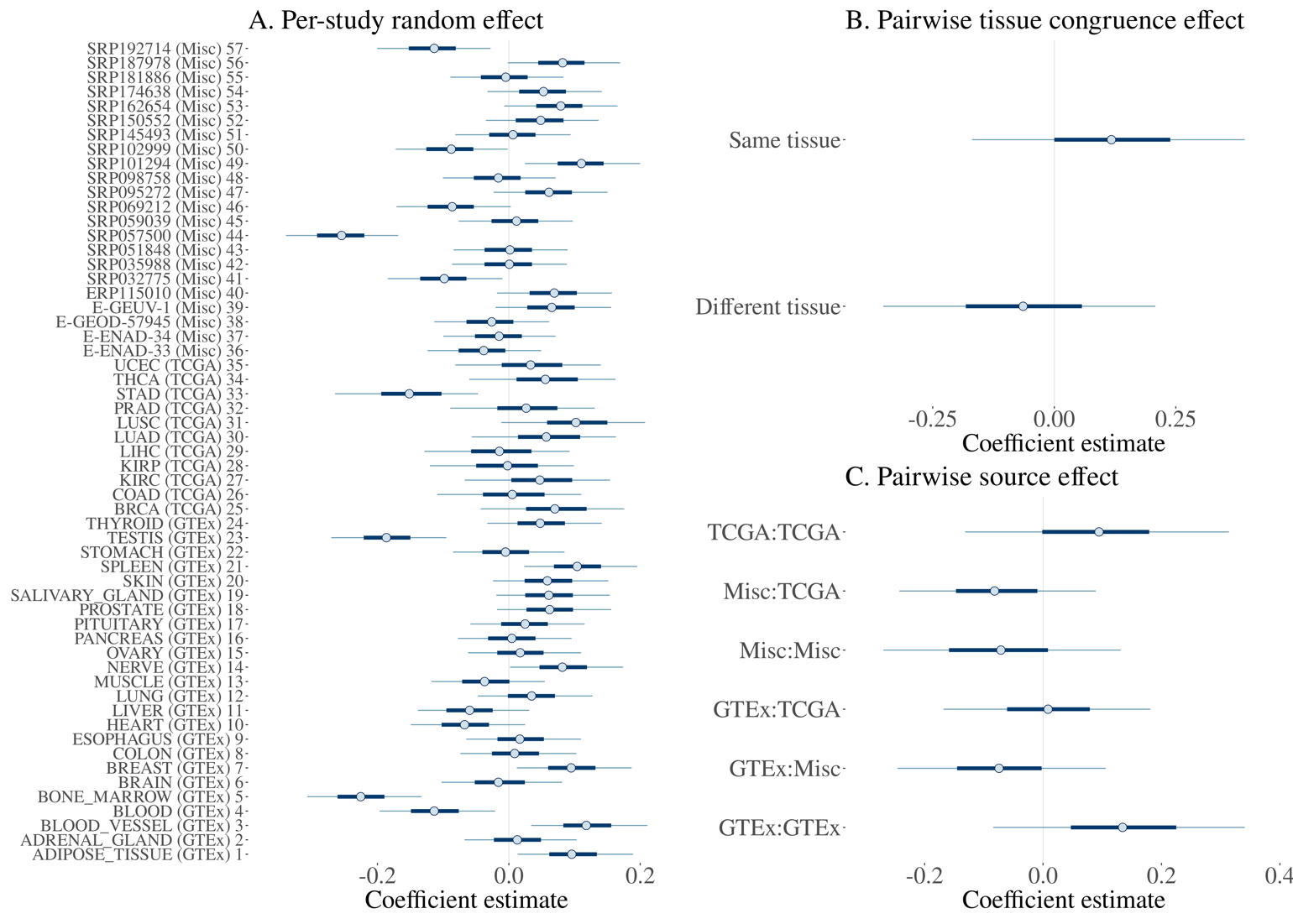

**Supplementary Figure 1**: Modeling the correlations between transcriptional variance across studies. The panels show coefficient estimates from a linear model using the among studies Spearman correlations between gene expression SDs as the response variable. These correlations are shown in fig. 2, panels A and B. In the linear model (see Methods for model equation), correlations are Fisher z-transformed. Study source and tissue are added as fixed effects. Coefficient estimates are shown with 50% and 95% credibility intervals. (A) The per-study random effect which accounts for the non-independence between the pairwise correlation values and estimates the characteristic contribution of each study to these correlations. For example, the lowest estimate among these parameters, which corresponds to the study bone marrow (from GTEx), indicates that correlations involving this study tend to be lower than the others. (B) and (C) Fixed effect estimates for the effects of tissue congruence and study-source effect. In (B) we see that correlations among studies that use the same tissue are slightly higher; and (C) correlations involving studies in the “Misc.” category (non-GTEx and non-TCGA) tend to be lower, while comparisons involving GTEx and TCGA are higher.

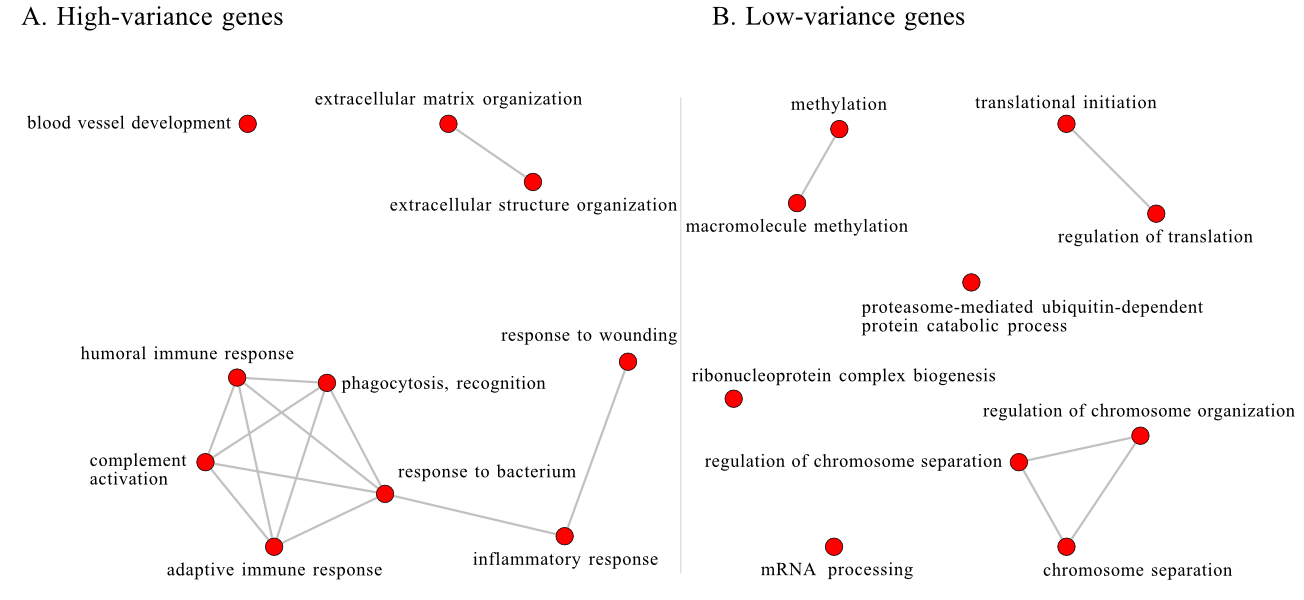

**Supplementary Figure 2.** Gene set enrichment analyses testing for over-representation of gene ontology categories in the upper and lower 5% quantiles of the gene variance rank. (A) High-variance genes are enriched for terms related to immune function, response to wounding, blood vessel morphogenesis, and inflammatory response. In contrast, (B) low-variance genes are associated with translation, control of methylation, RNA processing, chromosome separation, and other cell housekeeping functions. All displayed terms are significant with a 5% FDR corrected p-value below 10^-3^.

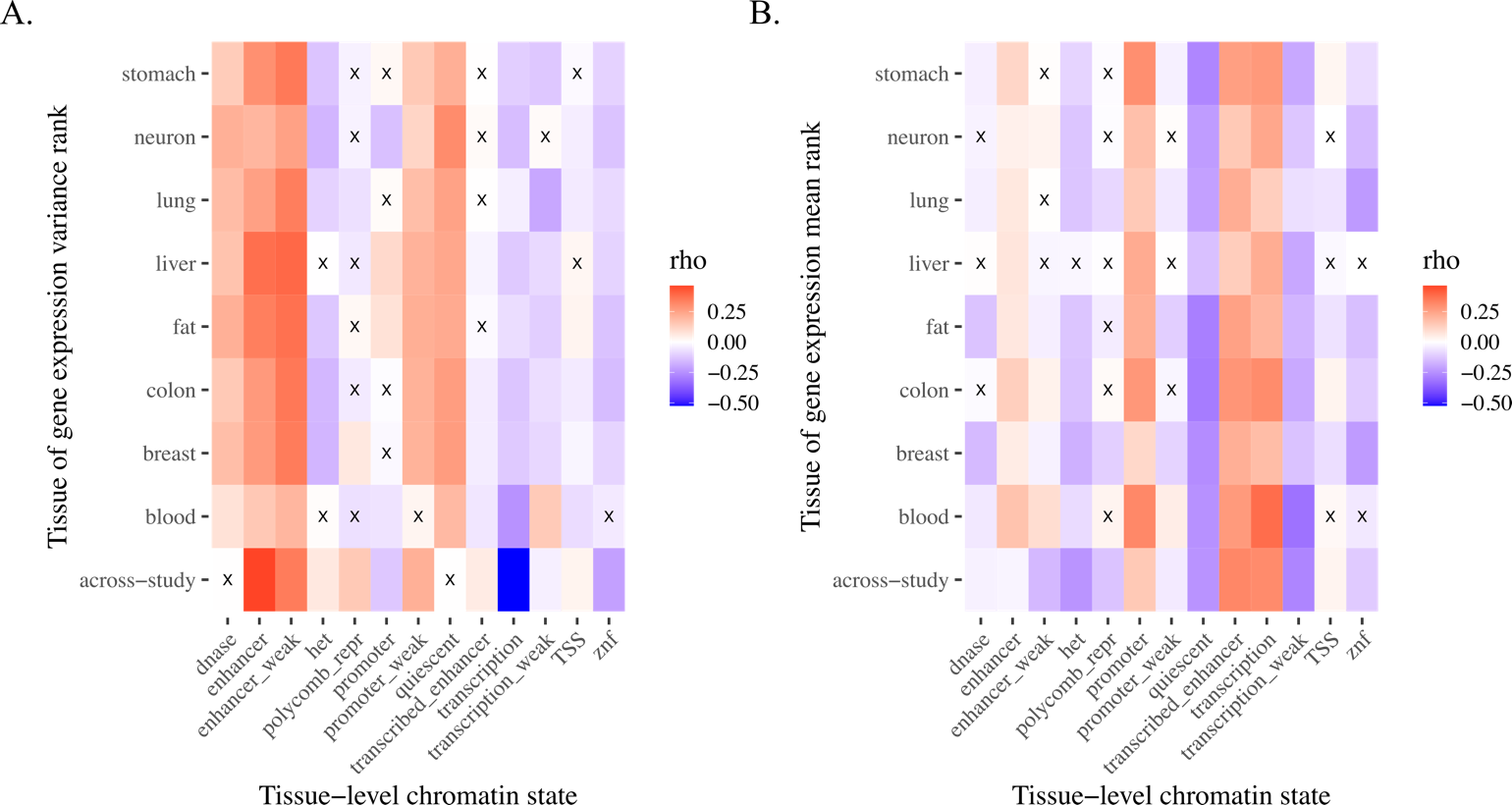

**Supplementary Figure 3. Across-study and tissue-specific gene expression variance and mean correlations with non-overlapping chromatin states through ChromHMM**^1^**.** The across-study variance (A) and mean (B) rank metrics (“across-study” on y-axis) were associated with universal chromatin states^2^ (x-axis). The tissue-level variance (A) and mean (B) rank metrics (see Methods; Supplementary Table 1; named tissues on y-axis) were associated with their respective tissue-specific chromatin states^3^ (x-axis, see Supplementary Table 1). Boxes marked with an “X” are not significantly correlated; all other comparisons are significant (Benjamini-Hochberg adjusted *p* < 0.05). Het indicates heterochromatin; TSS, transcription start sites; znf, zinc finger genes.

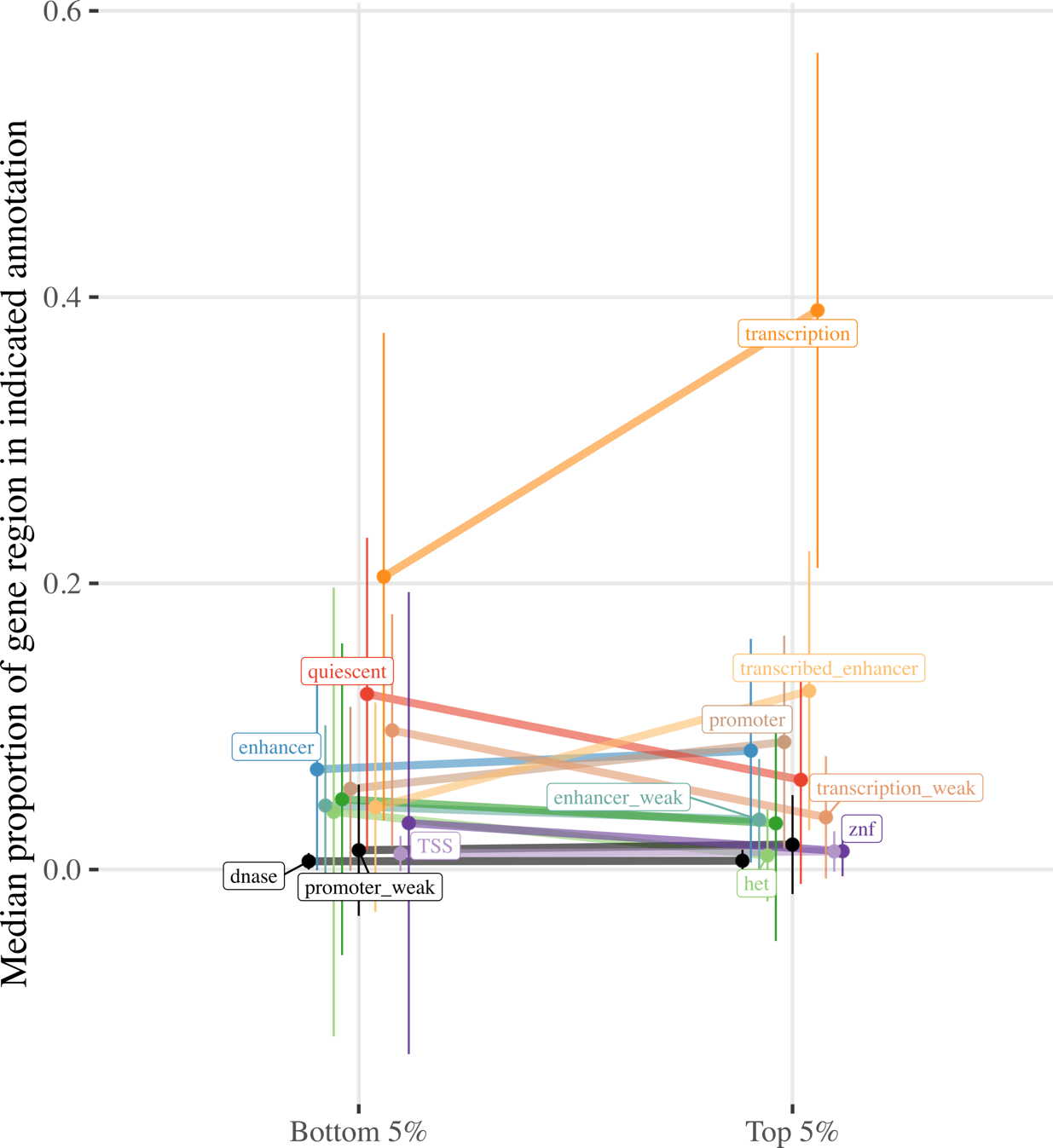

**Supplementary Figure 4. Proportion of gene regions made up of ChromHMM chromatin states for genes in the top and bottom 5% of the across-study mean rank metric.** Line plot contrasts the proportion of gene regions made up of the indicated chromatin states for genes in the top and bottom 5% of the across-study mean rank metric. Ends denote the median proportion of gene regions made up of the chromatin state, and error bars represent the standard error of the mean (SEM). States colored black are not significant, all others exhibit significant differences in gene region made up of the chromatin state for genes in the top and bottom 5% of the mean rank metric (Benjamini-Hochberg adjusted p_Wilcoxon_ < 0.05). Het indicates heterochromatin; TSS, transcription start sites; znf, zinc finger genes.

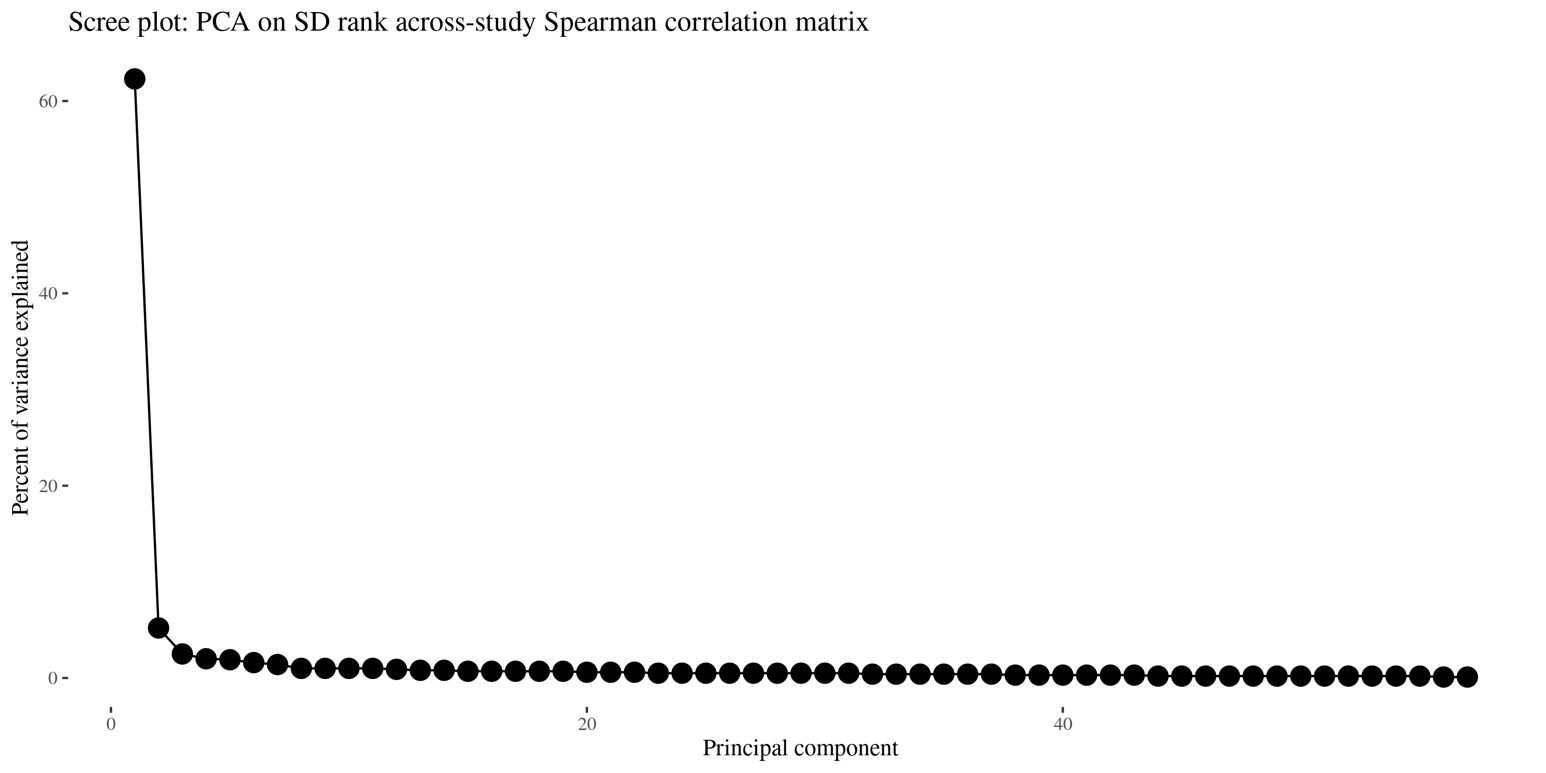

**Supplementary Figure 5. Scree plot on SD rank across-study Spearman correlation matrix.** Scree plot shows the percent of variance explained by each PC. These PCs are calculated on the across-study Spearman correlation matrix of across-study SD expression rank. PC1 accounts for 62.3% of variation and PC2, 5.2%.

**Supplementary Table 1. Variance and mean rank metrics and the corresponding ChromHMM annotations used.**

| **Variance/mean rank metric** | **ChromHMM annotation** | **Roadmap ID** |
| --- | --- | --- |
| Across-study | Universal^2^ |  |
| Blood | Primary mononuclear cells from peripheral blood | E062 |
| Breast | Breast Myoepithelial Primary Cells | E027 |
| Colon | Sigmoid Colon | E106 |
| Fat | Adipose Nuclei | E063 |
| Liver | Liver | E066 |
| Lung | Lung | E096 |
| Neuron | H9 Derived Neuron Cultured Cells | E010 |
| Stomach | Stomach Smooth Muscle | E111 |
